# An updated reference genome sequence and annotation reveals gene losses and gains underlying naked mole-rat biology

**DOI:** 10.1101/2024.11.26.625329

**Authors:** Dustin J Sokolowski, Mihai Miclăuș, Alexander Nater, Mariela Faykoo-Martinez, Kendra Hoekzema, Philip Zuzarte, Simon Monis, Sana Akhtar Alvi, Jason Erdmann, Archana Lal Erdmann, Rathnakumar Kumaragurubaran, Jonathan Bayerl, DongAhn Yoo, Nadia Karimpour, Kyra Ungerleider, Huayun Hou, Fergal J. Martin, Thibaut Hourlier, Zoe Clarke, Heidi E L Lischer, Dragos V Leordean, Yiyue Jiang, Trevor J. Pugh, Ewan St. J. Smith, Leanne Haggerty, Diana J. Laird, Jingtao Lilue, Melissa M. Holmes, Evan E. Eichler, Rémy Bruggmann, Jared T Simpson, Gabriel Balmus, Michael D. Wilson

## Abstract

The naked mole-rat (NMR; *Heterocephalus glaber*) is a eusocial subterranean rodent with a highly unusual set of physiological traits that has attracted great interest amongst the scientific community. However, the genetic basis of most of these traits has not been elucidated. To facilitate our understanding of the molecular mechanisms underlying NMR physiology and behaviour, we generated a long-read chromosomal-level genome assembly of the NMR. This genome was subsequently annotated and incorporated into multiple whole genome alignments in the Ensembl database. Our long-read assembly identified thousands of repeats and genes that were previously unassembled in the NMR and improved the results of routinely used short-read sequencing-based experiments such as RNA-seq, snRNA-seq, and ATAC-seq. We identified several spermatozoa related gene losses that may underlie the unique degenerative sperm phenotype in NMRs (*IRGC*, *FSCB*, *AKAP3*, *MROH2B*, *CATSPER1*, *DCDC2C*, *ATP1A4*, *TEKT5, and ZAN*), and an additional gene loss related to the established NK-cell absence in NMRs (*PILRB*). We resolved several tandem duplications in genes related to pathways underlying unique NMR adaptations including hypoxia tolerance, oxidative stress, and nervous system protection (*TINF2*, *TCP1*, *KYAT1*). Lastly, we describe our ongoing efforts to generate a reference telomere-to-telomere assembly in the NMR which includes the resolution of complex gene families. This new reference genome should accelerate the discovery of the genetic underpinnings of NMR physiology and adaptation.

## Introduction

The naked mole-rat (NMRs; *Heterocephalus glaber*), an African subterranean rodent, exhibits a suite of unique physiological traits, including longevity, cancer protection, hypoxia tolerance, reproductive suppression, poikilothermy, and resistance to multiple types of pain^1–6^. They also exhibit unique and behavioural idiosyncracies for a mammal such as their eusociality and caste system^1–6^. While subsets of these traits and adaptations are shared with other African mole-rats, NMRs are considered to have an extreme implementation of these traits^7,8^. For example, NMRs are the longest lived, display the most pronounced eusociality, and show extreme tolerance to hypoxic environments^7,8^. Further, NMRs thrive in laboratory environments, making them a valuable animal model to study numerous scientific questions in biomedical, ecological, physiological, and behavioural sciences^1–6^. Since the NMR’s designation as Science magazine’s “Vertebrate of the Year” in 2013^9,10^, the wider scientific community has taken a great interest in the molecular underpinnings of their unique set of adaptations and medically relevant traits, with particular interest in its healthy ageing, tumour resistance, and hypoxia tolerance^2,11–13^.

A crucial step towards discovering the genetic basis of unique NMR physiology was the release of the first NMR genome assembly in 2011^13^. This study relied on whole-genome shotgun sequencing and revealed the accumulation of mutations in vision-related genes leading to blindness through neutral evolution and amino acid substitutions in thermoregulatory-relevant genes *UCP1* and *HIF1A*, and has served as a foundation for all NMR researchers ever since^13^. The assembly was updated in 2014 using additional short reads, improving contiguity^14^. In 2020, this genome assembly was again updated by incorporating *in situ* Hi-C sequencing (HiC) and low pass (10x-coverage) long-read PacBio continuous long read (CLR) sequencing to aid in assembly scaffolding^15^. This improved genome helped reveal additional coding variations shared with another long-lived rodent, the Canadian beaver (*Castor canadensis*) and revealed regions of the genome undergoing accelerated evolution^15,16^.

The 2014 update of the 2011 assembly (HetGla1.2) has been the standard reference genome for the species since its release^17,18^. Gene and repeat annotations for HetGla1.2 have also been continually updated in new Ensembl and Refseq gene builds^17,18^, and in multiple sequence alignments (MSA) generated by Ensembl Compara and the Zoonomia project^19,20^. Accelerated genomic region analysis using HetGla1.2 also identified genomic regions related to optical development and vertebral column lengthening, the latter NMR adaptation being proposed to allow queens to produce large healthy litters many times throughout their lives^11^. However, it was noted that more than 66% of NMR accelerated regions are non-coding in origin and difficult to study using a fragmented genome assembly^11,21^.

While the current NMR assembly allows for the study of genic (coding) regions, genome assemblies based on long reads are needed to study repetitive, structural, and non-coding regions^22–24^. Such structural changes (tandem duplications, chromosomal inversions) have been shown to underlie new biological traits, such as the existence of ovotestes in the Iberian mole^25^. Work in other species with unique physiological properties related to their phylogeny (e.g., flight in bats, cancer resistance in elephants, nervous system development in octopus) has also demonstrated that changes in genome structure (i.e., ploidy, tandem duplications, gene loss, and new gene formation) and gene regulation (e.g., enhancers drastically impacting the expression of genes involved in unique NMR physiology) underpin unique lineage/species-specific phenotypes^26–29^.

Here we describe our efforts to create, annotate, align and share a new chromosomal-level scaffold reference that is now serving as the current NMR reference genome in the Ensembl database (https://www.ebi.ac.uk/ena/browser/view/GCA_944319725.1; TOR-NMR2626). We also describe our current progress generating a near telomere-to-telomere (T2T) quality genome that we intend to be the future update of the NMR genome (CAM-845F1). We use our initial T2T data to reveal the strengths and limitations of the current chromosomal-level scaffold genome presented here. We anticipate this resource will be of immediate use to the ever-growing community of researchers studying the unique biology of the naked mole rat.

## Results

### Assembly of HetGlaV3, the TOR-NMR2626 naked mole-rat assembly

To generate a new, diploid chromosomal-level reference NMR reference genome we utilized a trio-binning approach following the data types used in the Vertebrate Genome Project (VGP)^30^. We first generated 10X genomics linked (short) reads from the queen (i.e., “mother”, ID: TOR-NMR2606) and single consort (i.e. “father”, ID: TOR-NMR2624). We also generated short reads (linked reads) from one kin (a male offspring subordinate; ID: TOR-NMR2626) to perform genome polishing and k-mer-based genome assembly quality evaluation^31,32^. We focused our reference diploid assembly on the male subordinate (TOR-NMR2626) and generated Oxford Nanopore Technologies (ONT) R9.4 reads (∼40X coverage) to establish the backbone of our assembly. Lastly, we generated Hi-C sequencing data from the hypothalami of multiple female subordinates originating from the same colony and lineage as TOR-NMR2626. Our genome assembly pipeline also broadly follows the pipeline proposed in VGP^30^ (Materials and Methods. Supplementary Figure S1). Briefly, we used the parental short reads to assign TOR-NMR2626 ONT reads to their haplotype of origin with TrioCanu^31^. We combined haplotyped and unassigned reads to generate the maternal-haplotype and paternal-haplotype assemblies to compensate for the extensive homozygosity found in the NMR colonies (Supplementary Figures S1, S2, Figure 1B). Trio-binned reads were assembled with Flye and polished with racon (two rounds) and medaka before linked-read scaffolding was performed with Scaff10x (Materials and Methods)^33,34^.

**Figure 1.**
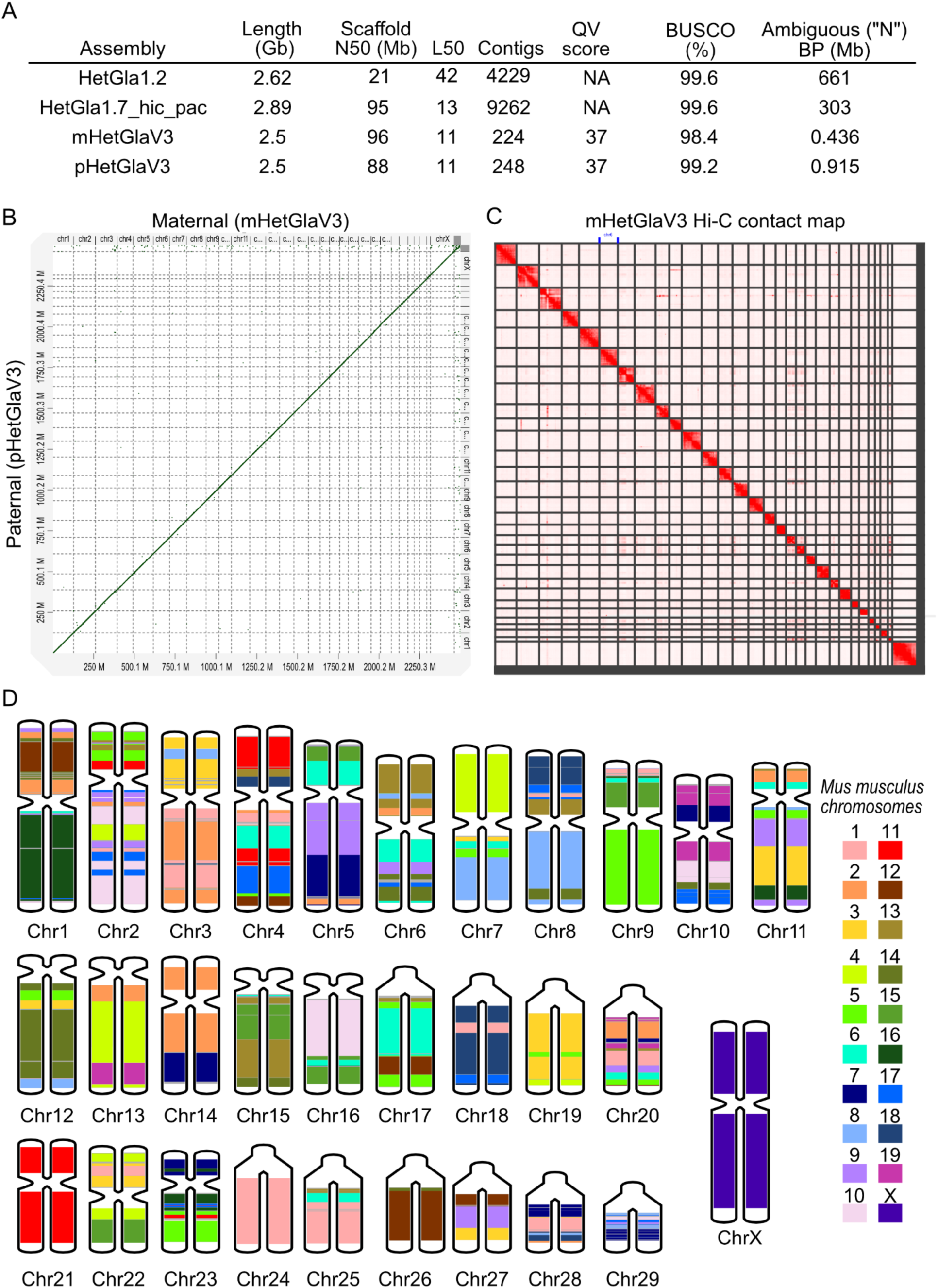
Global view of the de novo naked mole-rat assembly. A) Summary statistics for global assembly evaluation in the short-read based NMR HetGla1.2, the scaffolded short-read based naked mole-rat assembly HetGla1.7_hic_pac, and the diploid assembly presented in this study (mHetGlaV3 and pHetGlaV3). B) Genome-wide dot plot between mHetGlaV3 and pHetGlaV3 calculated using D-GENIES. C) Genome-wide Hi-C contact map of the mHetGlaV3 visualized with Juicebox. D) *in silico* karyotype of the naked mole-rat genome. Chromosome numbers match the physical karyotype. Chromosome painting using the mouse (mm10) genome as a reference, because it is the most commonly studied rodent in genomics. For example, the dark green in the Q arm of naked mole-rat chromosome 1 corresponds to regions with large alignment blocks to mouse chromosome 16.

We performed scaffolding with our Hi-C data using Salsa2 and 3d-DNA and the Juicebox assembly tools and assembly polishing using our short reads with freebayes (Supplementary Figure S1, Figure 1C)^35–37^. Lastly, we assembled a 16.4 kbp linear mitochondrial genome with 36 genes assembled using the mitoVGP pipeline^30^ and 1.5Mbp (predicted length ∼20Mbp) of the Y-chromosome including the *SRY* gene using a custom pipeline (Materials and Methods) (Supplementary Figure S3). Trio binning generated a maternal (mHetGla) and paternal (pHetGla) assembly for autosomes. The assembled X chromosome in mHetGla, Y chromosomal fragments in pHetGla, and mitochondria were added to both haplotypes. These haplotype assemblies were designated HetGlaV2 (mHetGlaV2: GCA_944319715.1, pHetGlaV2: GCA_944319725.1). mHetGlaV2 is currently the NMR reference genome in Ensembl 111 (https://useast.ensembl.org/Heterocephalus_glaber_female/Info/Index), and it is incorporated into “91 eutherian mammals” multiple whole genome alignment^17^.

We discovered a series of intrachromosomal scaffolding errors in HetGlaV2 (Supplementary Figure S1D, Supplementary Figures S4,S5). We fixed these scaffolding errors and placed centromeres using syntenic alignments to the VGP-assembled Canadian Porcupine (*Erethizon dorsatum,* EreDor1)^38^, and a recently published FISH karyotype for the NMR^39^ (Supplementary Figure S5). We designated this scaffold-corrected assembly HetGlaV3. We computed standard assembly quality score statistics between two prior short-read based assemblies (i.e., HetGla1.2, HetGla1.7_hic_pac) and our long-read based assemblies (i.e., HetGlaV2, HetGlaV3) (Figure 1A), where we found a considerable (700-fold) decrease in ambiguous (“N”) bases from HetGla1.2 to mHetGlaV3. We then performed *in silico* chromosome painting of HetGlaV3 to map mouse chromosomal regions to the NMR assembly (Figure 1D). In the subsequent sections of this manuscript our biological analysis of the NMR genome is performed on mHetGlaV3 (Supplementary Figure S6).

### A composite karyogram of diploid naked mole-rat cells as observed by traditional cytogenetics identifies chromosome order, centromeric location, and G-banding

Genomes assembled with long and accurate reads enable researchers to identify the sequence and location of complex, repetitive regions^40^. We generated a composite karyotype of the NMR genome (Figure 2) using over 350 cells from naked mole-rat cell lines including a juvenile male fibroblast cell line derived from a skin biopsy to obtain a physical map of chromosome size, centromere placement, and banding. Our composite karyotype provides examples of each chromosome at multiple banding resolutions. This allowed us to classify each chromosome into one of six groups related to chromosome size and centromere placement (Figure 2, Supplementary Table S1). These groups are broadly split into submetacentric chromosomes of various lengths, metacentric chromosomes of various lengths, and short acrocentric chromosomes. We found that chromosome 29 contains satellites using AgNOR sequential staining which highlights nucleolar organizing regions (Supplementary Figure S7). We assigned the chromosome names in HetGlaV3 to those identified in this composite karyotype by matching chromosome length and centricity.

**Figure 2.**
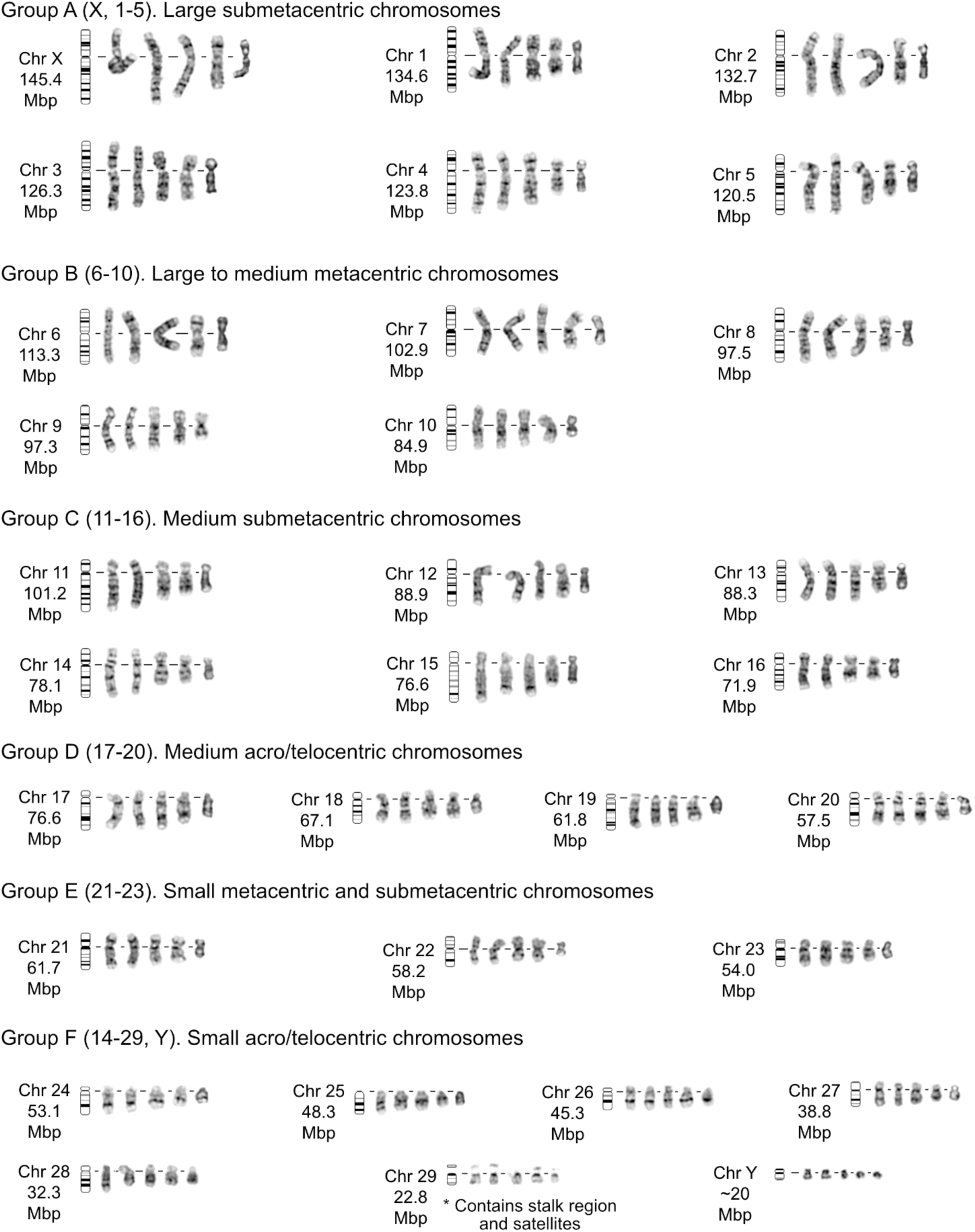
Composite image details the morphological characteristics of each chromosome group, as observed by traditional cytogenetics. Chromosome classes are grouped by centromeric position and ordered by descending sequence length. G-banded chromosome examples show a range of band resolutions. Ideograms illustrate landmark band locations.

### Gene Annotation of the NMR genome assembly

Overall, we identified 35,327 genes in mHetGlaV3 and assigned gene symbols to 21,043 genes. These include 7,500 gene symbols that were missing in Ensembl 111 and that were annotated using a combination of TOGA^41^ and manual curations (Supplementary Table S2,S3). We also found an additional 1,071 pseudogenes through duplication (343 of which were expressed in at least one tissue, 52 of which contained multiple exons) (Supplementary File S1, Supplementary Figure S8).

### Epigenomic annotation of the naked mole-rat hypothalamus

A considerable proportion of how a genome interacts with the environment and how differences in gene abundance can lead to adaptation is through the non-genic, regulatory elements of the genome^42,43^. We built a chromatin state map of the naked mole-rat hypothalamus to systematically profile promoters, enhancers, active regions, repressed regions, and insulators in the NMR hypothalamus epigenome (Supplementary Table S4, ChIP-seq metrics). Specifically, we performed ChIP-seq experiments investigating genome-wide H3K4me2 (promoter-enhancer associated), H3K4me3 (promoter-associated), H3K27Ac (active-enhancer/promoter associated), H3K27me3 (Polycomb repression complex associated), H3K36me3 (transcriptional elongation associated), and H3K9me3 (Silenced-associated) in female subordinate NMRs (N=2 per histone modification). Additionally, we performed ATAC-seq (N=3 female) and CTCF ChIP-seq (N=2 female from the entire brain). We integrated these ChIP-seq and ATAC-seq data with publicly available full-length RNA-seq of the NMR hypothalamus to build a chromatin state map using ChromHMM^44^. We identified the expected repertoire of mammalian chromatin states in a heterogeneous tissue^45,46^ (Figure 3, Supplementary Figure S9).

**Figure 3.**
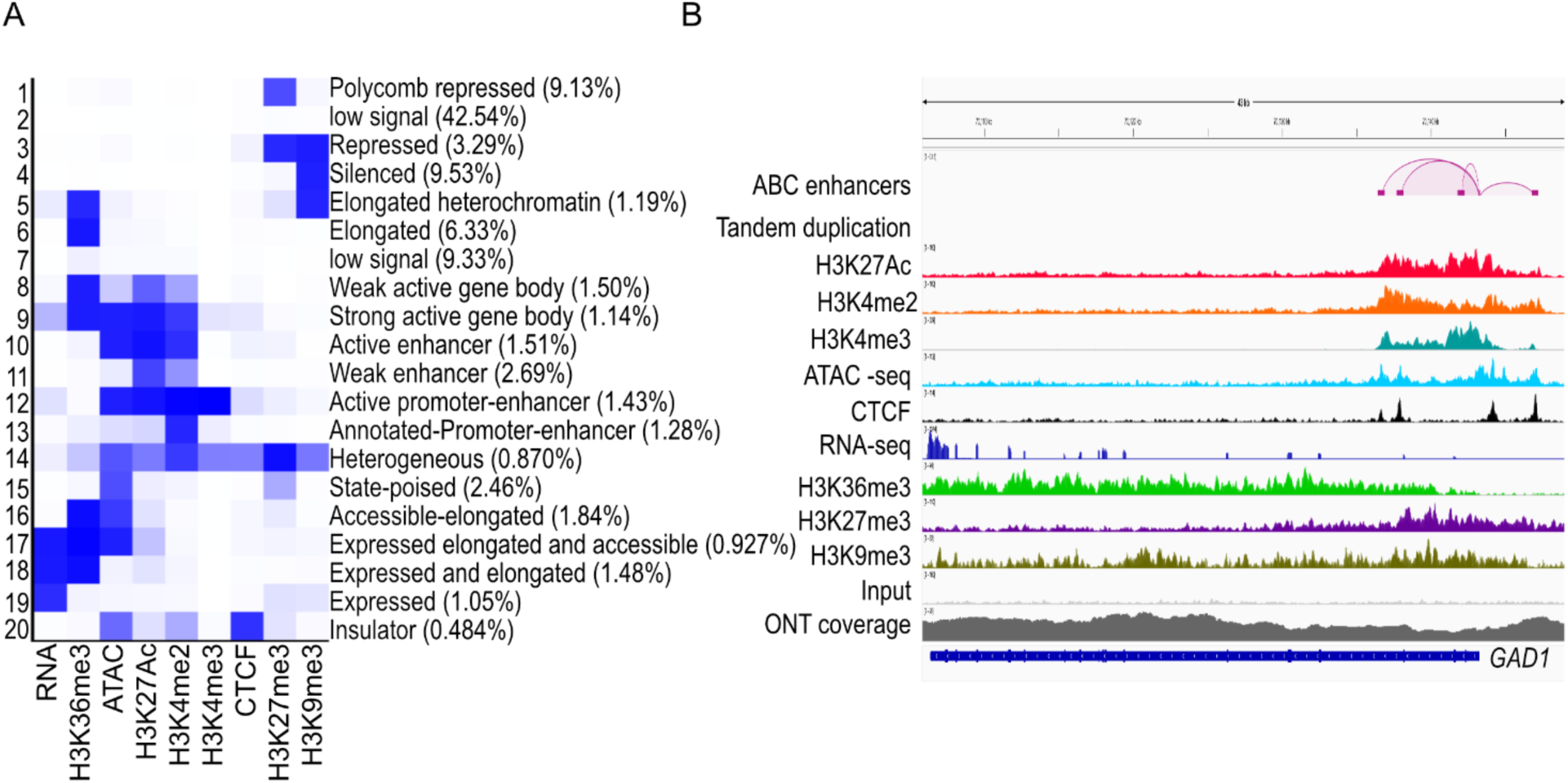
Evaluation of the chromatin state of the hypothalamus in the naked mole-rat genome (mHetGlaV3). A) Chromatin state emissions. Numbers in brackets show the proportion of the genome assigned to each chromatin state. The “heterogeneous chromatin state” (state 14) has activity of both H3K27me3 and H3K27Ac, representing regions with cell-type or cell-state dependent gene regulation. B) Genome browser snapshot of the *GAD1* gene, a canonical marker for GABAergic neurons, with its promoter region representing the “heterogeneous state”.

### Repeat annotation of the naked mole-rat

Transposable elements (TEs) play a critical role in genome evolution and there are several examples where species-specific TEs can influence organismal physiology and gene regulation^42,47^. Previous studies have analyzed the TE landscape of NMRs using HetGla1.2 and revealed NMR-specific TEs containing functional HNF4a and RAR/RXR motifs^42^. To ascertain how HetGlaV3 impacted TE annotations of the NMR we performed TE annotation of mHetGlaV3 and the previous NMR reference assembly (HetGla1.2) using Earl Grey^48^. Our assembly has a higher rate of TEs assigned than the short-read-based assemblies of HetGla1.2 (HetGla1.2 annotated repeats = 805.1Mbp, HetGlaV3 annotated repeats = 1,010.4 Mbp), which is consistent with prior work showing that HetGla1.2 under-represents the repeat landscape of the NMR^49^ (Figure 4A,B). This increase in called TEs is especially noticeable in LINE elements (HetGla1.2 = 16.27% of genome, HetGlaV3 = 22.92% of genome), which are longer on average than other TE classes (Figure 4A,B). We also identified a “peak” of LINE elements containing a Kimura distance from 16-19%, which is missing in the repeat annotation of HetGla1.2. This “peak” of LINE elements is present in other rodents, including mouse (*Mus musculus*, mm10, Figure 4A) further highlighting the importance of a long-read-based assembly backbone when investigating repetitive elements in the genome.

**Figure 4.**
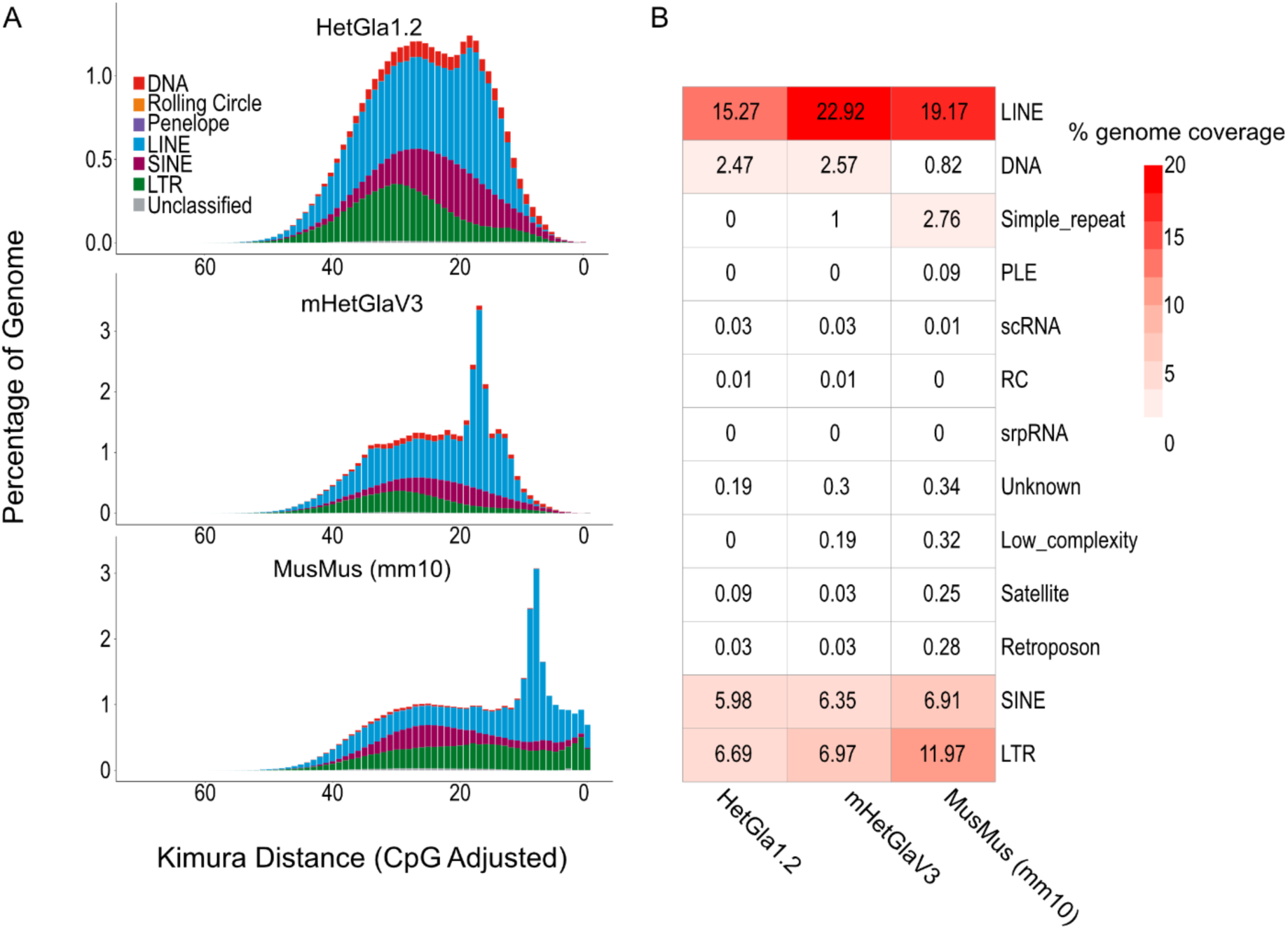
Evaluation of transposable elements (TE) in the naked mole-rat genome. A) Landscape plot displaying the distribution of TEs in HetGla1.2 (top), mHetGlaV3 (middle) and mouse (mm10) (bottom). The Y-axis represents the percentage of genome coverage, and the X-axis represents the Kimura distance of the TE, which reflects the rate of sequence transversion from the TE in the NMR genome and the best matching TE in the Dfam database. The lower the Kimura distance, the lower the rate of sequence transversion, and the more likely that the TE is evolutionarily young. Colour represents the TE class. B) Heatmap of the percentage of total genome coverage for each TE class in HetGla1.2, mHetGlaV3, and mm10.

### Evaluation of genome assembly using RNA-seq and ATAC-seq analysis

With our genome assembled and annotated, we investigated what impact our genome assembly would have when analyzing several commonly used genomics data types and analysis pipelines. We focused on analyzing bulk paired-end RNA-seq with a high read depth (∼150 million reads per sample)^50^ and ATAC-seq of twelve total NMR hypothalamus samples. For each dataset, we compared the differences in aligned reads and detected features (i.e., genes, peaks) between each assembly. We found a higher number of RNA-seq^50^ reads aligned to mHetGlaV3 than HetGla1.2 (range 4.3%-20.3%) and an 18.1% increase in genes with at least 0.5 RPKM in one tissue (Figure 5A-C). We also identified a higher rate of multi-mapping reads in mHetGlaV3 than HetGla1.2 (mean 9.5-fold increase, 3.75-14.9 fold-increase), reducing false alignment rates in HetGla1.2. We found the same pattern in our ATAC-seq data, where we found more reads aligned to HetGlaV3 than to HetGla1.2 (mean 2.6%, 1.98-3.48%) but more peaks called in HetGla1.2 than HetGlaV3 (mean 11.6%, or 5323 more peaks, 7.06-16.8%) (Figure 5E-F). These “HetGla1.2 specific” peaks are mostly aligned to LINE elements (55-70%), with the remaining peaks aligning to other TEs (Figure 5G-H). Therefore, these “HetGla1.2 specific” peaks reflect read pile-ups on misassembled (i.e., collapsed) repeats in HetGla1.2 and are likely false-positive peaks. Improvements in data quality remained consistent with snRNA-seq data in the hypothalamus (42,961 cells, 39 clusters, N=8) (Faykoo-Martinez et al. in preparation), where we saw an increase in gene counting, feature detection, and cell-marker detection and identification (Supplementary Figure S10).

**Figure 5.**
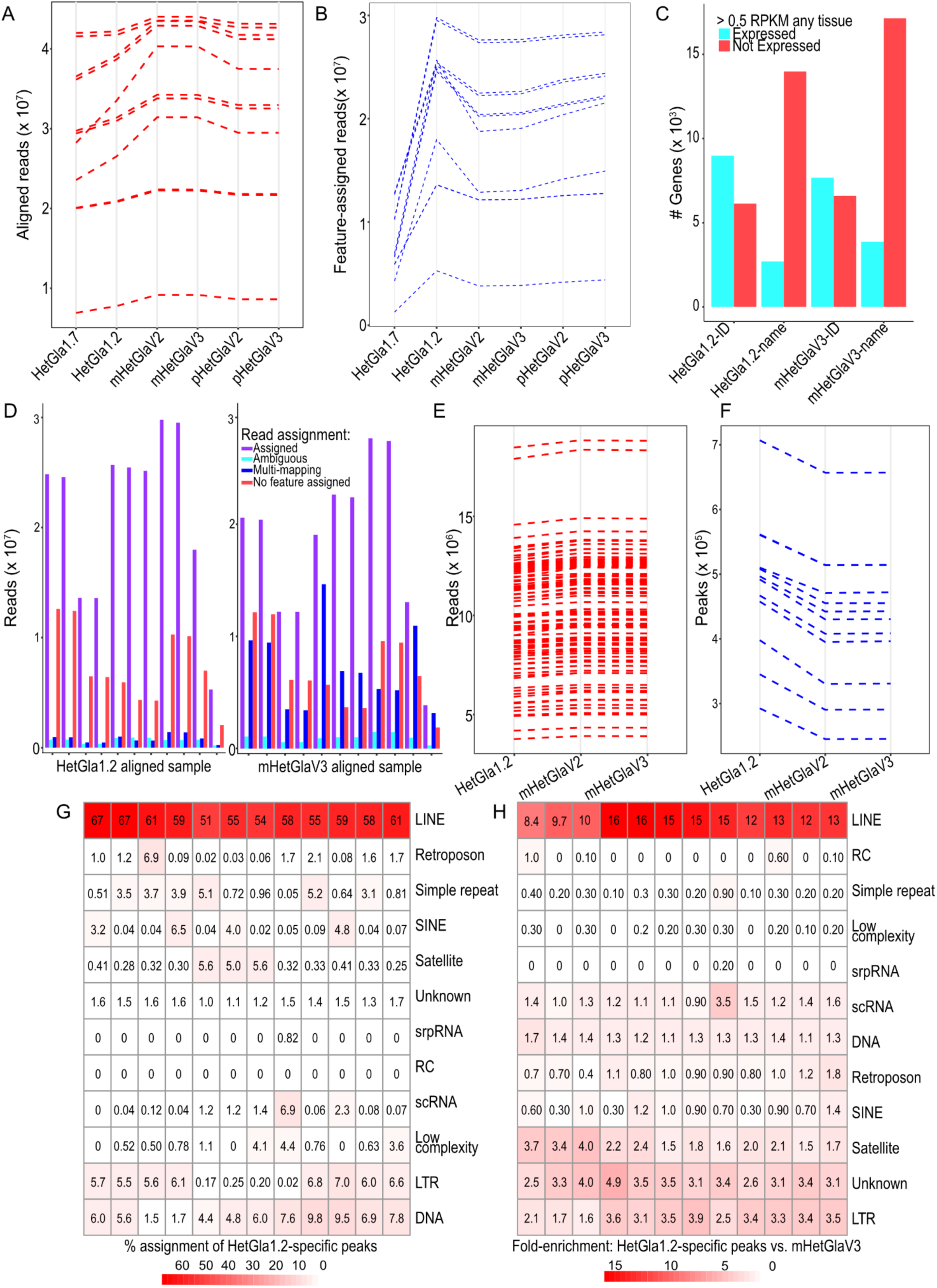
Comparison of bulk RNA-seq and bulk ATAC-seq alignment and annotation between NMR assemblies. A-B) Total number of bulk RNA-seq reads aligned to each publicly available NMR assembly (A) or assigned to a gene feature (B). “HetGla1.2” and “HetGla1.7_hic_pac” (gap-filled HetGla1.2) are short-read based, while the HetGla assemblies are the long-read based assemblies published in this study. Each line represents the same bulk RNA-seq sample from Bens et al., 2018. C) Comparison of the number of expressed and non-expressed genes that are assigned a gene symbol between mHetGlaV3 (our primary assembly) and NMR2011. D) Distribution of reads assigned to features in HetGla1.2 (left) and mHetGlaV3 (right). Each group of bar plots is a different sample from Bens et al 2018. Purple bars are assigned reads. Light blue bars are unassigned due to ambiguity. Blue bars are unassigned due to multimapping. Red bars are unassigned because the read did not map to a feature in the assemblies annotation file. E-F) Comparison of ATAC-seq reads (E) and number of ATAC-seq peaks (F) from 12 samples in the NMR-hypothalamus (this study) when aligned to HetGla1.2, mHetGlaV2, and mHetGlaV3. G-H) Repeat annotation of the HetGla1.2-specific peaks when projected onto the mHetGlaV3 assembly. Each row is a different repeat class and each column is a different NMR sample. G) populates the heatmap with the percentage of HetGla1.2 peaks assigned to each repeat family. H) populates the heatmap with the fold-enrichment of each TE compared to the universe of ATAC-seq peaks using the gat enrichment tool.

### Analysis of the updated NMR genome reveals gene losses related NMR phenotypes

Some of the physiological features of NMRs are driven by trait loss, such as their blindness, which we explored using TOGA^41,51^. We found many inactivated genes (i.e., pseudogenization through point mutation) involved in sensory perception and vision and sperm morphology (Supplementary Figure S11). Compared to other mammals, naked mole-rats have degenerate and atypical sperm morphology^52,53^. This trait is hypothesized to be caused by reduced selection pressure on sperm motility/competition that is prevalent in many rodent species^54^ but absent in NMRs due to their eusociality (a long lived queen who breeds with one or two consorts)^52,53^. Our TOGA analysis identified clear gene disruption events in several sperm morphology and male infertility genes: *IRGC*, *FSCB*, *AKAP3*, *MROH2B*, *CATSPER1*, *DCDC2C*, *ATP1A4*, and *TEKT5*. Indeed, mouse knockout studies of *IRGC*, *AKAP3*, *CATSPER1*, *ATP1A4* show sperm morphology and male infertility phenotypes^55–59^. TEKT5 was recently shown to be a component of the microtubules that form the sperm tail in an unbiased cryo-EM/AlphaFold2 driven study performed in mice^60^. Furthermore, consistent with the other NMR gene losses, knocking out *Tekt5* in mice results in sperm motility defects^60^. FSCB is a post meiotic-expressed sperm/testes specific protein that localizes to mouse mouse sperm flagella^61^ and its phosphorylation increases mouse sperm motility *in vitro^62^*. *MROH2B* was identified as a gene induced during spermatocyte development whose expression was lost in *Hsf5* knockout mice ^63^. *DCDC2C* has been identified as part of the human sperm microtubulome^64^. A *Dcdc2c* mouse knockout phenotype released by the International Mouse Phenotyping Consortium identified an increased heart weight and vertebrae shape phenotypes but no infertility phenotype^65^. However sperm morphology/motility has not yet been assessed in these animals.

For this gene loss analysis we called gene inactivation events conservatively. In our analysis, we required an inactivation in mHetGlaV3 and pHetGlaV3, and that the region was also annotated with the Ensembl pipeline. If we do not require an Ensembl annotation, we find additional gene inactivation events. For example, our analysis using TOGA^41^ identified the pseudogenization of the gene encoding zonadhesin, *ZAN*. We previously showed that *Zan was* needed for species specific sperm egg interactions using a mouse mouse knockout model^66^. These extensive gene inactivations related to sperm morphology echo the prior observations where gene inactivations in vision related genes are related to blindness in NMRs^13^. Further analysis of these gene inactivation events will provide a comprehensive list of sperm-related pseudogenization in the NMR.

Previous work has also shown that NMRs lack canonical natural killer cells and are missing gene family expansions involved in natural killer cell regulation, such as the *LILR, KLR,* and *MHC* gene families^67^. We have replicated these findings in mHetGlaV3 by visualizing the syntenic blocks surrounding these regions in the mouse (Supplementary Figure S12). We also identified an additional gene loss of *PILRB,* a gene which is known to undergo expansions in rodent genomes as part of a host-pathogen evolutionary arms race^68^. In the NMR, this gene loss arose from structural rearrangements (Supplementary Figure 13). Notably, *PILRB* is involved in immune system activation in natural killer cells (which are not present in NMRs)^69^. As the quality of the NMR genome improves, gene losses involving rapidly evolving gene loci and structural rearrangements will become easier to identify^70^.

### Tandem NMR duplications include tumour resistance and hypoxia response genes

Gene expansion and contraction through tandem duplication and gene loss are powerful candidates to drive species-specific adaptation and physiology^26–29^. These events can lead to new gene creation through gene fusion, and paralog formation^26–29^. They can also lead to changes in gene dosage larger than an additive influence of copy number due to the gene being in a new regulatory context^26–29^. However, tandem duplications can be difficult to annotate from a genome assembly and annotation standpoint due to the repetitive nature inherent to these regions^71^. We, therefore, took a conservative approach to identify high-confidence tandem duplications in mHetGlaV3. Briefly, we first identified segmental duplications assigned to different annotated genes or CDS regions within 100kbp from each other. We called these segmental duplications tandem duplication if it 1) had no evidence of misassembly, 2) did not contain many-to-many mapping to the mouse, 3) showed evidence of gene regulation and expression in at least one copy, 4) passed manual curation (Materials and Methods for details). We identified 16 loci (spanning 20 genes) with a highly-confident tandem duplication of genes. Each candidate was investigated manually (Table 1, Supplementary File S2). While many of these could be related to adaptation, we highlighted tandem duplications of regions containing genes with intuitive links to unique NMR physiology, namely *TINF2*, *TCP1*, and *KYAT1* (Figure 6).

**Figure 6.**
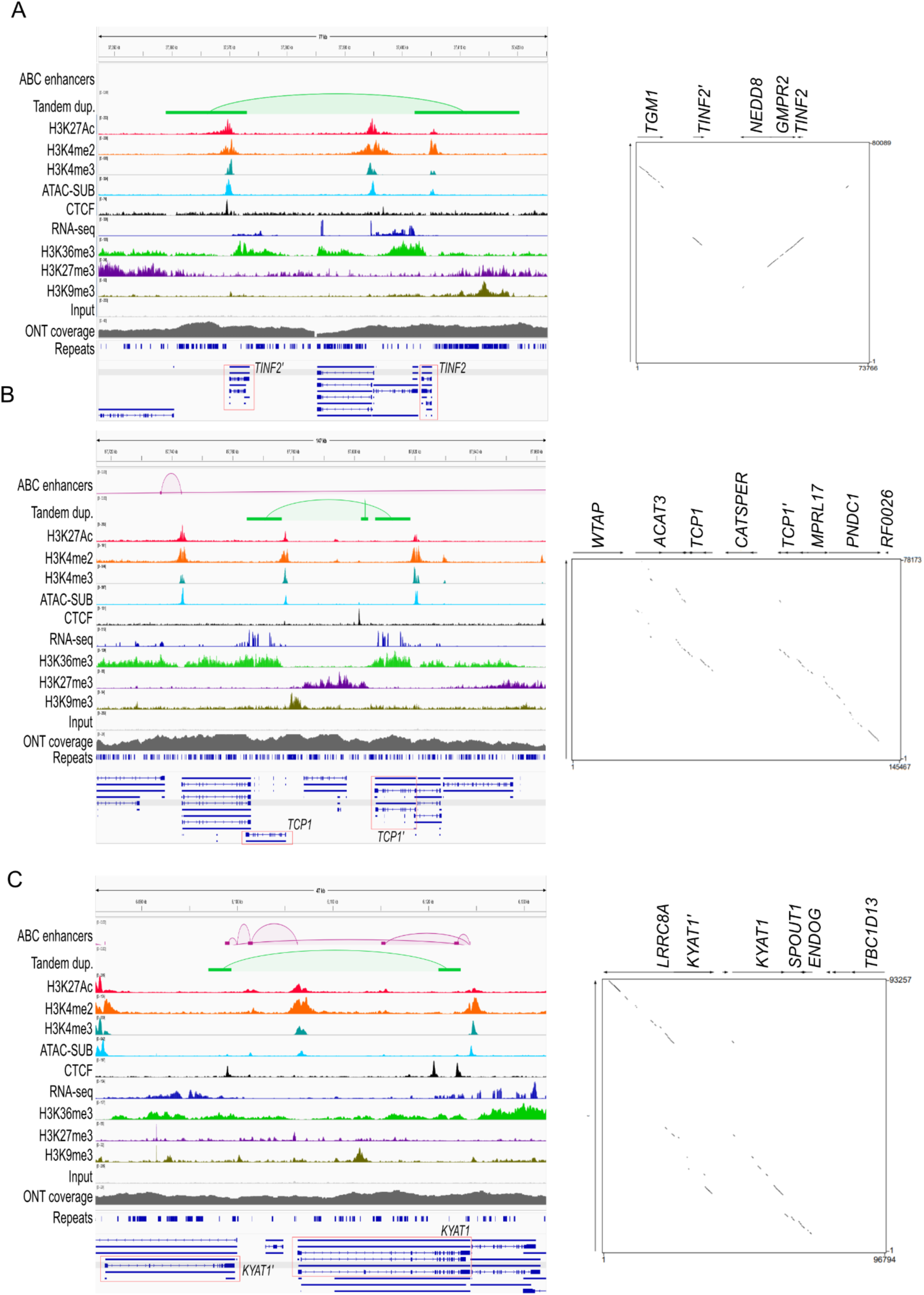
Visualisation of four loci containing tandem duplications where both gene copies contain gene expression and regulation in the NMR. A-D) Genome browser screenshots of loci containing the *TINF2* (A), *TCP1* (B), *KYAT1* (C) genes. Screenshots on the left show NMR data projected onto the mHetGlaV3 assembly, and the right represents dot-plot between the same genomic regions between mHetGlaV3 and mm10.

**Table 1.** Summary of the 20 genes with tandem duplications in mHetGlaV3. Gene expression, the presence of epigenetic regulation, and the presence of an Activity-by-Contact (ABC)-enhancer are specific to gene regulation in the hypothalamus. “Gene expression” reflects hypothalamic gene expression in the parent and the child gene. “Epigenetics” reflects whether the predicted child gene has clear chromatin-state annotations in the hypothalamus. “ABC enhancer reflects” whether the parent and child genes have ABC enhancers between the parent and gene copy or, in the case of *DDT*, from a different segmental duplication. RNA-seq, ChIP-seq, and ATAC-seq datasets in this analysis were restricted to uniquely mapping reads.

| Gene | Genome co-ordinates<br>(mHetGlaV3) | Gene<br>Expression | Epigenetics | ABC enhancer |
| --- | --- | --- | --- | --- |
| *ARF4* | chr6:85,003,219-85,055,214 | Yes | Active | No |
| *CEL* | chr13:3,321,953-3,345,504 | Low | Repressed | No |
| *COG1* | chr21:48,098,201-48,200,795 | Yes | Active | No |
| *CST6* | chr20:23,432,729-23,448,065 | Low | Elongated | No |
| *CTRB1* | chr7:91,436,127-91,492,524 | Low | Low | No |
| *DDT* | chr23:38,197,873-38,313,130 | Yes | Active | Yes |
| *EPHB6* | chr20:46,648,036-46,963,435 | Low | Repressed | Low |
| *FABP1* | chr4:57,701,358-57,769,171 | Low | Low | No |
| *FMO4* | chr24:15,893,506-16,009,869 | Low | Low | No |
| *KCTD14* | chr5:89,138,875-89,170,998 | Yes | Active | Yes |
| *KEL* | chr20:46,648,036-46,963,435 | Low | Low | No |
| *KYAT1* | chr13:6,086,859-6,133,961 | Yes | Active | Yes |
| *LCN2* | chr13:7,349,575-7,373,773 | Low | Low | No |
| *LLCFC1* | chr20:46,648,036-46,963,435 | Low | Low | No |
| *NUDT16* | chr11:32,214,427-32,268,393 | Low | Repressed | No |
| *SPIN2B* | chrX:62,619,182-62,657,958 | Yes | Low | No |
| *TCP1* | chr2:87,715,383-87,860,850 | Yes | Active | No |
| *TINF2* | chr1:37,352,142-37,425,908 | Yes | Active | No |
| *TRPV5* | chr20:46,648,036-46,963,435 | Low | Low | No |
| *TRPV6* | chr20:46,648,036-46,963,435 | Low | Low | No |

MacRae et al., 2015, identified a *TINF2* duplication, a shelterin complex member and a telomere stabilization gene, in HetGla1.2^72^. In NMRs, *TINF2* activates in low-oxygen conditions to aid cell survival and drive responses to hypoxia; however, they were unable to investigate gene expression differences in each *TINF2* copy and only reported an overall increase in *TINF2* expression in NMRs^72,73^. Our assembly also identified the *TINF2* duplication, and we could uniquely map reads to each *TINF2* copy (*TINF2* and *TINF2’*), enabling further investigation into this region. *TCP1* is a chaperonin-containing TCP1 complex (CCT) member and helps regulate protein folding, and it has been shown to impact tumour growth in multiple contexts^74,75^. Lastly, *KYAT1* is important in forming kynurenic acid, which is neuroprotective in the mouse and human brain^76^. We also identify an Activity-by-Contact (ABC) enhancer^77^, described as a promoter-enhancer interaction using complementary evidence from Hi-C-seq, ATAC-seq, H3K27Ac, and RNA-seq data^77^, from one copy of *KYAT1* to the other, suggesting that this tandem duplication may also be influencing gene regulation. NMRs have previously shown a species-specific, strong response to kynurenine compared to mice^78^.

### Primary assembly of a telomere-to-telomere quality naked mole-rat genome assembly

Recent technologies, namely PacBio HiFi, ONT R10.4, and ONT ultra-long reads allow drastic improvements in genome assembly, both at structural and individual base-pair levels^79–81^. These technologies were used to complete the human genome, yielding a telomere-to-telomere (T2T) genome assembly^40^. We are working towards generating a similarly complete genome in the naked mole-rat. We collected state-of-the-art PacBio HiFi reads and ONT ultra-long reads from a new NMR (ID:CAM-845F1), which we assembled with hifiasm^82^ (Figure 7A, “Materials and Methods”). We found a scaffold N50 of 98.0Mb, a total length of 2.63Gb, 0.1 ambiguous (“N”) base pairs per 100Kb, substantially outperforming earlier assemblies (i.e., mHetGlaV3 = 71.3, HetGla1.2 = 11,589.4), and a BUSCO score of 97.9% (Figure 7B). Using Merqury^32^, we found a k-mer completeness score of 98.83, and a QV score of 64.48, which reflect a predicted base-pair error rate of one error in every 2.8Mbp (Figure 7B). The CAM-845F1 and HetGlaV4 show a high degree of structural consistency, with clear syntenic alignments (Figure 7C, upper panel). Conserved syntenic regions emphasise the existing evolutionary relationship between the human(33), mouse^83^, and NMR T2T genome assemblies, but also highlight unique genomic regions in the three species (Figure 7C, lower panel). Our chromosome-level scaffolding is consistent with HetGlaV3, and with the NMR FISH-karyotype^39^. Briefly, our genome annotation of HetGlaV4 involved transferring Ensembl gene models from HetGlaV3 to HetGlaV4 using LiftOff before integrating additional gene models from TOGA, BRAKER3, and StringTie2 using mikado in reference mode^41,84–87^. Here, we identified an additional 3,345 protein coding genes and 6,038 CDS regions. We also identified 2,091 gene symbols that were previously uncharacterized in *HetGlaV3*. These genes are over-represented for the olfactory receptor gene family (29 genes, FDR-adjusted p-value = 4.97 x 10^-12^).

**Figure 7.**
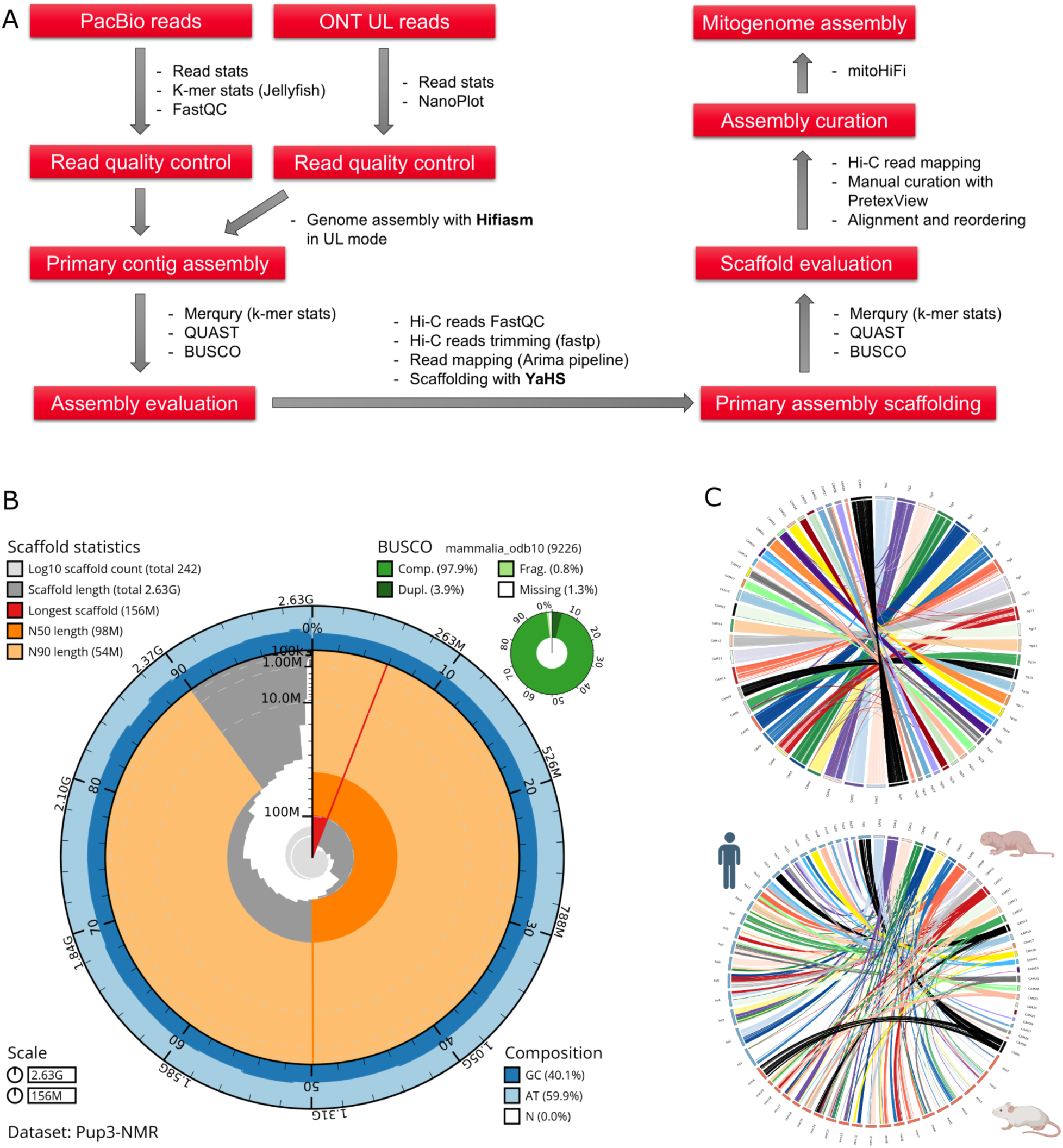
Overview of the naked mole-rat primary genome assembly using sequencing technologies from the telomere-to-telomere genome era. A) Genome assembly and evaluation pipeline of the naked mole-rat using Pacbio HiFi and ONT ultra-long reads. B) Assembly composition plot showing GC content, missingness, assembly and scaffold lengths, and BUSCO score. C) Circos plot comparing CAM-845F1 (HetGlaV4) (left) and HetGlaV3 (right). D) Ciros plot comparing the human, mouse and CAM-845F1 T2T genome assemblies.

The HetGlaV3 assembly failed to resolve several important repeat-dense genomic regions. Four notable examples of these regions are 1: olfactory receptor (*OLFR*) gene families, immunoglobulin gene families, and potential tandem duplications (i.e., *SLC25A2*). We found that HetGlaV4 successfully resolved these regions, finding a 13Mbp OLFR gene expansion on HetGlaV3 that was resolved to 17 Mbp and placed on the correct chromosome (mHetGlaV3 placement: chromosome 2, HetGlaV4 placement: chromosome 12) (Supplementary Figure S14), a 1.4Mbp immunoglobulin region in mHetGlaV3 was 2.4 Mbp in HetGlaV4 (Supplementary Figure S14), that *SLC25A2* does not show a tandem duplication, despite evidence of a gene copy-number variation in HetGlaV3 (Supplementary Figure S14). Lastly, we found evidence that genes surrounding *TINF2* may also be duplicated in HetGlaV3, namely *MDP1*, *CHMP4A*, and *TSSK4*, however, the presence of assembly gaps prevented us from confidently reporting this tandem duplication. In contrast, HetGlaV4 displays a single 40kbp tandem duplication of five genes, *TINF2, MDP1*, *CHMP4A*, *NEDD8*, and *TSSK4*, where each gene has two copies in the NMR (Supplementary Figure S14). With the NMR T2T assembly, designated HetGlaV4, we and other researchers will be able to explore complex regions of the NMR genome and provide a completed genome reference for the community to use for the foreseeable future.

## Discussion

The NMR has a suite of highly unusual physiological traits encoded in their DNA. However, many of the genetic underpinnings of these traits remain unresolved. The short-read-based genome assemblies do not provide sufficient resolution of structural variations, repeat regions, and regulatory regions. Our study helps resolve these challenges by providing a de novo diploid genome assembly of the NMR that integrates long and short reads to optimize assembly structure (i.e., long-reads) and individual base pair accuracy. By integrating complete genome assemblies from related species and a karyotype characterized by fluorescent in situ hybridization^38,39^, we generated karyotype-level chromosomal scaffolding (Figures 1,2). Our genome assembly is paired with a comprehensive genome annotation of genic and repeat elements in our assembly^41,48,50,84^. We also built a chromatin state map^44^ of the NMR hypothalamus to profile regulatory logic surrounding enhancers, genes, and repeat elements (Figures 3,4).

HetGlaV3 showed improvements in contiguity, gene annotation, gene symbol calling, and repeat annotation compared to the previous short-read-based NMR assembly. These improvements lead to improved gene mapping, false-positive peak detection from bulk RNA-seq and snRNA-seq respectively (Figure 5), and improved gene detection and marker identification in single-nuclei RNA-seq data (Figure 5, Supplementary Figure S10). Our primary T2T-level assembly, HetGlaV4, showed improved resolution of complex genic and non-genic regions such as gene family expansions, tandem duplications and centromeric regions. Therefore, it is imperative to use the highest quality genome assembly possible when studying the NMR genome regardless of the sequencing experiment being performed (e.g., differential expression in traditional RNA-seq analysis, ATAC-seq peak calling, etc.). Our discovery of gene inactivation in sperm morphology genes, potentially as a function of reduced selection pressure for highly mobile sperm due to eusociality, shows that NMRs are a relevant model to study the genetics of fertility from the perspective of sperm morphogenesis and evolution^52–54^.

Our analysis of tandem duplications identified genes implicated in hypoxia tolerance (e.g., *TINF2*, *DDT*)^72,88^ or oxidative stress/reactive oxygen species (ROS) (e.g. *TINF2*, *TRPV6*, *LCN2*, *ARF4*, *DDT*, *FMO4*) in model placental organisms including NMRs^72,88–92^. In addition, *TCP1*, which has two expressed gene copies with independent promoters and enhancers, functions as a protein chaperone, and has been linked to tumour progression in humans^75,93^. To our knowledge, *TCP1* has not been implicated in hypoxia tolerance, water retention, or rROS in mammals. However, *TCP1* has been shown to regulate ROS neutralization in the Leishmania donovani parasite^94^, respond to hypoxic environments in *Solea solea* fish^95^, and interact with *Gli1* in hypoxic tumour environments^96^. A number of these genes with tandem duplications in NMRs have also been shown to play a functional role in tumour progression, tumour suppression, or, in the case of *NUDT16*, double-stranded break response^97^. Not every identified tandem duplication contains genes known to influence hypoxia tolerance or oxidative stress. For example, *CEL* encodes a pancreatic enzyme responsible for hydrolyzing fat, cholesterol esters, and fat-soluble vitamins^98^. *CEL* also has copy-number polymorphisms in humans related to both pancreatitis and a form of diabetes named Carboxyl-ester lipase maturity-onset diabetes of the young (CEL-MODY)^99,100^.

The NMR exhibits a suite of traits that are of interest to biomedical scientists, from healthy ageing to cancer resistance. When investigating the genetic underpinnings of these medically relevant traits, it may be valuable to consider them in the evolutionary and ecological context of the NMR. For example, many of the NMR-specific structural, regulatory, and protein changes found within this study and others encode genes whose function has been implicated in hypoxia tolerance, oxidative stress response, or ROS defence, and in tumour growth or longevity^42,101,102^. Other pleiotropic, adaptation-relevant genes have also been seen at the protein level in the NMR. For example, *p14ARF* has been linked to both oxidative stress response and tumour suppression in the NMR^103^, and a gain of function mutation in *Has2* (hyaluronan acid synthase) has been linked to longevity in the NMR^104^. Recently, a NMR-Has2 knock-in model is one of a very short list of genes to increase longevity in mice^104^. From the perspective of the NMR, these medically relevant traits possibly arose from selection pressures resulting from living in arid environments and in low-oxygen microenvironments where for the most part they cannot be reached by predators^105^. These medically relevant traits could have occurred through pleiotropic effects of genes and pathways related to NMR adaptation and ecology (e.g., hypoxia tolerance, oxidative stress response, water retention). These speculations have previously been proposed in NMRs in reference to their cancer resistance^106^. They have also been proposed for the blind mole rat genus, *Spalax*^106,107^, another long-lived, cancer-resistant, subterranean rodent who evolved these traits independently from African mole rats. Accordingly, pathways involved in both medically relevant physiological traits and ecologically relevant adaptations may be strong candidates when studying the genetic underpinnings of longevity and tumour resistance in the NMR.

Our current genome annotation integrates existing published NMR RNA-seq data^50^ with state-of-the-art computational genome annotation approaches^17,41,84^. Our improved assembly and annotation allowed us to identify 3,498 (11.0%) more gene models than the previous NMR assembly annotated by Ensembl (HetGla1.2) and build more complete gene models. We also annotated the naked mole-rat hypothalamus epigenome with a chromatin state-map, annotating promoters, enhancers, transcribed regions, repressive states, and insulators (Figure 3). These efforts will be improved by using long-read RNA-seq (e.g., Iso-Seq) which would also reveal transcript structure that short-read RNA-seq-based transcript models may miss^108^. Iso-Seq data will be released with HetGlaV4. Our genome and annotation will improve comparative genomic analyses that are essential for studying non-coding and structural genomic elements driving physiology and adaptation^12,20,109,110^.

While our updated publicly available HetGlaV3 assembly is a significant improvement over previous reference assemblies, it is not yet complete. For example, our ongoing efforts to create a telomere-to-telomere NMR assembly using additional long-read sequencing technologies, including PacBio HiFi reads, and ONT “ultra-long” reads substantially increase contiguity and individual base-pair accuracy (Figure 7). These long reads, paired with the karyotype-level scaffolding of our NMR, are getting us closer to a telomere-to-telomere assembly (T2T)^40^. These newer, longer reads should resolve difficult to assemble regions such as complex gene families, centromeres and interspersed segmental duplications^111^ that likely underlie NMR-specific biology.

## Materials and Methods

### Animal and tissue collection

#### NMR handling

**Toronto:** Naked mole-rats were bred in-house and maintained in stable colonies (i.e. established breeders were present) housed in large (45.75 cm L × 24 cm W × 15.25 cm H) and small (30 cm L × 18 cm W × 13 cm H) polycarbonate cages connected by plastic tubes (25 cm L × 5 cm D). Animals were fed sweet potato ad libitum supplemented with wet 19% protein mash (Envigo RMS, Inc.) and kept on a 12:12 light/dark cycle at 28–30 C. All procedures were approved by the University Animal Care Committee and performed in accordance with federal and institutional guidelines. **Cambridge:** Naked mole-rats were bred in-house and maintained in an inter-connected network of cages in a humidified (∼55 %) temperature-controlled room at 28-30° C, with red lighting (08:00-16:00) and had access to food *ad libitum*. In addition, a heat cable provided extra warmth under 2-3 cages/colony. Experiments were conducted under the Animals (Scientific Procedures) Act 1986 Amendment Regulations 2012 under a Project License (P7EBFC1B1) granted to E. St. J. Smith by the Home Office and approved by the University of Cambridge Animal Welfare Ethical Review Body. **San Francisco:** Animals were kept on a 7:17 light/dark cycle at 76-86F and at a humidity of 30-70%. Cages (25 cm L × 17.78 cm W × 28 cm H) were made of 1/4" thick polycarbonate walls, hardware was stainless steel and joints made of solvent welded. NMRs had access to heat lamps for a minimum of 7 hours a day during the light cycle. Animals were fed sweet potato ad libitum, and rotational items (ie. Variety of fruits and vegetables, garbanzo beans, wet 19% protein mash (Envigo RMS, Inc), wet baby cereal (Gerber)). All animal care and experimental procedures were in accordance with federal policies and guidelines governing the use of animals and were approved by the University of California–San Francisco’s IACUC. The IACUC follows the guidelines in the 8th edition of The Guide for the Care and Use of Laboratory Animals^112^.

### NMR sample collection

#### Linked short-read and nanopore long-read sequencing (Toronto)

For linked short-read sequencing, we collected gonad tissue from a female breeder (four years old) (TOR-NMR2606) and a male breeder (5.5 years old) (TOR-NMR2624), and two of their offspring, namely a female subordinate (1 year old) (TOR-NMR2625) and a male subordinate (1 year old) (TOR-NMR2626). The same male subordinate sample was used for the nanopore long-read sequencing. Animals were anesthetized with isoflurane and rapidly decapitated before a single gonad was harvested and flash-frozen in isopentane on dry ice.

#### Hi-C sequencing (Toronto)

Twenty subordinate (wild-type) female NMRs between 1-4 years of age were anaesthetised with isoflurane and rapidly decapitated. The brain was sliced coronally into 1 mm sections on an acrylic block. Sections were transferred to cold PBS, and the hypothalamus was removed using a scalpel blade and flash-frozen whole in isopentane on dry ice.

#### ChIP sequencing (Toronto)

The hypothalamus for eighteen NMR subordinates (14 female, 4 male) were collected for ChIP-sequencing using the same approach as in Hi-C sequencing. For ChIP-seq of CTCF, we collected the whole brain of two NMRs.

#### ATAC sequencing (Toronto)

Twelve NMR hypothalamus samples between two and four years of age were collected using the same method as in Hi-C and ChIP-sequencing. These twelve NMRs were divided into subordinates (SUB), and subordinates were removed from the colony for one week (EXSUB-wk1), in both males, and females (N=3 per developmental stage/sex).

#### Nanopore ultra long read sequencing (Cambridge)

Ultra-high molecular weight DNA was extracted from the spleen tissue of a female, male, and their male pup using a modified phenol/chloroform DNA extraction method.

#### PacBio HiFi long read sequencing (Cambridge)

High molecular weight DNA was extracted from frozen whole blood with the NEB Monarch cells and tissue kit, following manufacturer’s instructions.

### Naked mole-rat sequencing

#### Linked short-read sequencing (Toronto)

Tissue for DNA extraction was homogenized in cell lysis buffer [10mM TRIS (pH8.0), 25mM EDTA, 100mM NaCl, 0.5% SDS, RNase A] and digested with proteinase K overnight at 37C. The resulting lysate was extracted twice with TRIS-saturated phenol (pH8.0) followed by a single extraction with chloroform. DNA was precipitated from the solution by adding 1/10th volume 3M Sodium Acetate and 2.5 volumes of 95% ethanol, followed by gentle inversion. Precipitated DNA was spooled onto a glass rod and washed with excess 70% ethanol, and allowed to partially air dry before being resuspended in an appropriate volume of EB [10mM TRIS (pH8.0)]. We followed The Genome Reagent Kits v2 User Guide, Protocol Step 4 (https://support.10xgenomics.com/genome-exome/index/doc/user-guide-chromium-genome-reagent-kit-v2-chemistry) library construction protocol with no changes. Then, The Genome Reagent Kits v2 User Guide, Protocol Step 5 (https://support.10xgenomics.com/genome-exome/index/doc/user-guide-chromium-genome-reagent-kit-v2-chemistry) nucleic acid sequencing protocol was followed exactly with no changes to perform genome sequencing.

#### Nanopore (R9.4) long-read sequencing (Toronto)

High molecular weight DNA was extracted from the same sample and tissue as the male-linked short-reads sample using a standard phenol/chloroform DNA extraction protocol. DNA for library preparation was sheared to an average size of 10kb on a Covaris g-TUBE. Libraries were prepared with the Oxford Nanopore SQK-LSK109 library prep by ligation kit according to the manufacturer’s instructions. The resulting libraries were quantified by fluorometry using a Qubit High Sensitivity DNA quantification kit (ThermoFisher), and 100 fmol of each resulting library was used to load an R9.4 PromethION flow cell.

#### Hi-C sequencing (Toronto)

Hi-C experiments were performed using the Arima Hi-C kit (Arima Genomics) according to the manufacturer’s instructions. Briefly, pooled flash-frozen brain tissues were cross-linked with 2% formaldehyde for 20 min at RT in buffer containing 100mM NaCl, 1mM EDTA, 0.5mM EGTA, 50mM HEPES (pH8.0) in 1X PBS. DNA content was estimated, and 3ug was used as input per biological replicate. Chromatin was digested with an enzyme cocktail included in the kit. Ends were filled-in in the presence of biotinylated nucleotides, followed by subsequent ligation. Ligated DNA was sonicated using the Diagenode Bioruptor Pico (10 Cycles of 30 Secs On/Off) to an average fragment size of 400 bp DNA was then subjected to a double-size selection to retain DNA fragments between 200 and 600 bp using Ampure XP beads (Beckman Coulter). Biotin-ligated DNA was precipitated with streptavidin-coupled magnetic beads (included in the kit). The library was prepared using Swift Biosciences Accel-NGS 2s Plus library kit (Cat # 21024). Final amplification before sequencing was done with KAPA Library Amplification Kit ( Cat # KK2620). The cDNA library was sequenced on the Illumina NovaSeq 6000 S4 flow cell to generate paired-end 150 bp reads at The Centre for Applied Genomics (TCAG).

#### ChIP sequencing (Toronto)

Tissue was dounced in freshly prepared solution A ( 1% formaldehyde, 50mM Hepes-KOH, 100mM NaCl, 1mM EDTA, 0.5mM EGTA) and incubated for 20 mins at room temperature. The formaldehyde was then quenched with 1/20 volume of 2.5M glycine and incubated for 5 min. Tissue was then rinsed with ice cold PBS twice and centrifuge at 4 C at 2500 x rcf for 5 min.

Crosslinked tissue was then lysed using Chromatin Easy Shear Low SDS Kit (Diagenode #C01020013). Chromatin was sheared into 100–500 bp DNA fragments by sonication (Diagenode Bioruptor Pico) for 8 cycles of 30s ON and 30s OFF. Approximately 1.5% of chromatin was used for input DNA extraction. For each ChIP, chromatin lysates were combined with 10ug of anti-H3K27ac (Active Motif #39133), anti-H3K4me2 (Millipore # 07-030), H3K27me3 (Millipore #07-449), H3K36me3 (Abcam #ab9050) and H3K9me2 (Abcam #ab8898) antibodies and incubated overnight rotating at 4C.

For library preparation, all the ChIP DNA and input DNA were mixed with reagents from NEBNext Ultra II Library Prep Kit (NEBNext #E7645S) according to the manufacturer’s protocol.This DNA was then amplified with barcoded primers for Illumina sequencing (NEBNext® Multiplex Oligos for Illumina® #E7335S). PCR amplifications were carried out as [98 °C 30 s, (98 °C 10 s, 65 °C 75s) ×9cycles, 65 °C 5 min, 4 °C hold]. The amplified and barcoded library was then purified and selected for 200–350-bp fragments using AMPure XP beads (BECKMAN COULTER #A63881) and sequenced on IlluminaNOVAseq S4 flowcell with a 150-bp run to obtain 50 million reads paired-end per sample.

#### ATAC sequencing (Toronto)

NMR hypothalamus were dissected, flash frozen and stored at -80C for up to four months. Tissue was homogenised in a dounce homogenizer in PBS, then washed 2x with ice-cold PBS. Cells were counted with a Countess^TM^ using 1:1 dilution of Trypan Blue. ATAC-Seq was performed according to Buenrostro, et al. (2015)^113^. Briefly, 100000 cells were used for each sample and resuspended in 50uL of lysis buffer (10 mM Tris-HCl, pH 7.4, 10 mM NaCl, 3 mM MgCl2, 0.1% IGEPAL CA-630). Cells were spun for 10 mins at 600xg and 4C to prepare a crude nuclei extract. Nuclear pellets were then resuspended in transposition mix (25uL 2X TD buffer, 2.5uL TD [Illumina #20034198], and 23uL nuclease-free water) and incubated in a thermomixer at 37C for 30 mins shaking at 800rpm. DNA was isolated using the DNA Clean & Concentrator-5 kit (Zymo #D4013) and eluted in 11.5uL of 10mM Tris-HCl, pH 8.0. Libraries were prepared using 10uL of each sample. To generate libraries 50uL PCR reactions were made using 25uL NEBNext® High-Fidelity 2X PCR Master Mix(NEB #M0541L), 2.5uL of universal Ad1_noMx primer, 2.5uL of Ad2.x barcoded primer and 10uL nuclease-free water. Reactions were run according to recommended conditions with an annealing temperature of 63C and amplified for a total of 5 cycles. qPCR was subsequently run using 5uL of each 50uL PCR reaction to determine the number of additional cycles choosing a cycle number which provided ⅓ maximum fluorescence value. After subsequent PCR cycles, libraries were cleaned using a two-sided AMPureXP bead clean up using 0.5X then 1.3X beads. Libraries were eluted in a final volume of 21uL of 10mM Tris-HCl, pH 8.0, and quantified with the DNA-HS Qubit assay. Libraries were sequenced on Illumina NOVAseq S4 flowcell with a 150-bp run to obtain 50-100 million paired-end reads per sample.

#### Nanopore ultra long read sequencing (Cambridge)

Libraries were prepared with the Oxford Nanopore SQK-ULK001 ultra-long library prep kit, following the manufacturer’s protocol, with modifications to DNA input and library clean up. The resulting libraries were loaded onto an R9.4 PromethION flow cell and sequenced for 72 hours with two washes and reloads 24 and 48 hours.

#### PacBio HiFi long read sequencing

A starting amount of 4-5 ug HMW gDNA was sheared to a target size of 20-30 kbp using a Megaruptor 3 instrument (Diagenode). The sheared DNA underwent size selection using a Pippin HT instrument (Sage Science) to target a size range of 15-22 kbp. Following size selection, the DNA was used for CCS (circular consensus sequencing) library preparation using the SMRTbell Express Template Prep Kit 2.0 and Enzyme Cleanup Kit 1.0 (PacBio). Each library was barcoded using PacBio Barcoded Overhang Adapters. After library preparation, the concentration of the DNA stock was measured using the DNA-HS Qubit assay, and the DNA size was estimated using the Femto Pulse. Sequencing was conducted on a PacBio Sequel IIe instrument, using version 2.0 sequencing reagents and operating on control software version 10.1.0.119549, with a movie collection time of 40 hours per 8M SMRT Cell.

### Naked mole-rat *HetGlaV3* genome assembly *(TOR-NMR2626)*

#### Autosomal and X chromosome assembly

Our genome assembly pipeline blends the Vertebrate Genome Project and DNA-zoo pipelines^38,114^ (Supplementary Figure S1). First, we partitioned individual long-reads to maternal, paternal, or ambiguous (unassigned) haplotypes using TrioCanu (canu/2.1) with a provided genome size of 2.7 gigabases ^31^. Furthermore, the eusocial nature of naked mole-rats paired with the inbreeding that comes with an in-lab animal leads to considerable homozygosity and, as a byproduct, a larger proportion of reads that cannot be haplotyped when compared to wild animals or humans (Supplementary Figure S2). Accordingly, the maternal *Hetercephalus glaber* (mHetGla) assembly included maternal + unassigned reads, while the paternal *Hetercephalus glaber* (pHetGla) assembly included paternal + unassigned reads. mHetGla and pHetGla nanopore reads were assembled with the flye/2.8.2 using the –nano-raw parameter^115^. Assembled contigs were then polished three times, twice with racon/1.4.21 and once with medaka/1.4.3 (https://github.com/nanoporetech/medaka)^33,34^. For each round of racon polishing, trio binned reads were aligned to each assembly with bwa/0.7.17 (-x ont2d) before racon was performed (-m 8 -x -6 -g -8 -w 500)^116^. We completed Medaka polishing with default parameters using the “r941_prom_hac_g507” model. We then performed an initial round of scaffolding for the mHetGla and pHetGla with linked short reads. Briefly, assembled contigs longer than 50kbp were indexed with longranger/2.2.2 (https://github.com/10XGenomics/longranger) before linked short-reads were aligned to the contigs. Linked short reads were then used to connect contigs using Scaff10x/4.1 (https://github.com/wtsi-hpag/Scaff10X) with default parameters.

Next, we used Hi-C seq to scaffold contigs into chromosomes. We merged Hi-C replicates and aligned them to the Scaff10x-level mHetGla and pHetGla using juicer 2.0 using default parameters^117^. Then, we used juicer “sam_to_pre” and Juicebox assembly tools (JBAT) to visualize our Hi-C data at the Scaff10X level mHetGla and pHetGla^36^. We converted these Hi-C alignments into bam files and sorted bed files with samtools/1.9 and bedtools/2.27.1 respectively^118,119^, for Salsa2 scaffolding. We then performed 10 iterations of the salsa2 (version 2.2) algorithm using the Arima restriction enzyme mixture (GATC, GAATC, GATTC, GACTC, GAGTC). Hi-C data were aligned to the Salsa2-level mHetGla and pHetGla using juicer 2.0 and JBAT/2.17.00 for visualization and input into the next assembly step^35,36,114,117^. A second round of scaffolding was performed on the Salsa2-level mHetGla and pHetGla using 3d-DNA (version 180922), using the “haploid” module and only including contigs greater than 10Kb. Hi-C data were then aligned to the 3d-DNA-level mHetGla and pHetGla using juicer 2.0 and visualized with JBAT^36^. Chromosomal boundaries in the 3d-DNA-level mHetGla and pHetGla were adjusted and evaluated using JBAT^35,36,114,117^.

The 3d-DNA-level mHetGla and pHetGla were aligned with minimap2/2.1, using parameters for intra-species assembly alignment (-cx asm5)^120^. Alignment between 3d-DNA-level mHetGla and pHetGla identified inconsistent chromosomal boundaries across haplotypes. We selected the assembly with a more confident chromosomal boundary and adjusted the other assembly using JBAT^36^. For example, if two pHetGla “chromosomes” aligned evenly to one mHetGla “chromosome”, and the pHetGla had one more “chromosome” than expected given previously published NMR karyotypes (i.e. 31 scaffolds)^39^, the pHetGla “chromosomes” were merged. After scaffolding and visualization, we completed additional genome polishing with long and short reads. Specifically, the scaffolded mHetGla and pHetGla were polished twice with racon and once with medaka using the same parameters as the contig-level assembly^33,34^. Then, we aligned the 10X-short reads from the male proband to the scaffolded assemblies using longranger. Haplotype-aware variant calling of aligned short read data was completed with freebayes/1.3.1( --skip-coverage $((mean_cov*12)))^37^ and filtered (’QUAL>1 && (GT="AA" || GT="Aa")’) using bcftools/1.11^121^. The filtered variants were applied to modify the sequence of each assembly using bcftools consensus^121^. Intra-chromosomal assemblies and inversions between the final mHetGla and pHetGla (mHetGlaV2 and pHetGlaV2) were identified by aligning each haplotype with minimap2^120^ and visualizing alignments with an interactive dot plot using D-Genies^122^.

#### Mitochondrial and Y chromosome assembly

We used the mitoVGP/2.0 pipeline to identify an array of MT-derived nanopore reads, polish them using nanopore and 10X short read data, and trim and overlap the MT-derived array to generate a single, complete, and polished MT genome^30^. MT-derived tRNAs were identified with tRNAscan-SE^30,33,34,120,123,124^.

We used four versions of our genome assemblies to identify Y-chromosome-derived contigs. Specifically, we used mHetGlaV2 and pHetGlaV2, a contig-level assembly using paternal and unassigned reads, and a contig-level assembly using paternal reads only. Both contig-level assemblies were polished with two rounds of racon, one round of medaka, and one round of freebayes with the same parameters as in mHetGlaV2 and pHetGlaV2^33,34,38^. We designated these additional two assemblies as paternal-unassigned-contig (PUC) and paternal-contig (PC). We aligned 10X short reads from each NMR sample (mother, father, male subordinate, and female subordinate) to each of these assemblies using longranger (https://github.com/10XGenomics/longranger). The number of reads from each sample assigned to each contig within each assembly was assigned using samtools idxstats^118^. Within each assembly, we measured sex differences in reads aligned to each contig using DESeq2^125^. Contigs with a strong paternal bias of aligned reads were designated as candidates for Y chromosomes (log2-fold change > 2, FDR-adjusted p-value < 0.05). We then aligned Y chromosome contig candidates from pHetGlaV2, PUC, and PC to mHetGlaV2 using minimap2 (-cx asm5)^120^. Contigs with greater than >15% alignment to mHetGlaV2 were filtered. This filtering procedure also results in pseudo-autosomal contigs being assigned to the X chromosome instead of the Y chromosome. Lastly, we overlapped Y-chromosome contigs from PF, PUC, and PC to evaluate the amount of Y-chromosome captured. Y-chromosome Contigs from PC was then scaffolded with one round of scaff10X, one round of Salsa2, and one round of 3d-DNA^35,38,114^. For Hi-C scaffolding, we used publicly available Hi-C-seq reads from a male NMR fibroblast cell line^15^ because our Hi-C dataset was from a female NMR hypothalamus. Scaffolded Y contigs were then polished with two rounds of racon, one round of medaka and two rounds of freebayes using the same cutoffs as the chromosome-level assembly^33,34,38^. Y chromosome contigs and the MT genome were included in mHetGlaV2 and pHetGlaV2.

We employed syntenic analysis, chromosome painting, and a publicly available FISH-Karyotype^39^ to fix scaffolding errors and place centromeres in our genome assembly (Figure 1F-G). First, we split the chromosomes in mHetGlaV2 and pHetGlaV2 into draft scaffolds computed by 3d-DNA^114^. We then computed syntenic blocks between this assembly and the Canadian Porcupine (EreDor1) genome assembly (https://www.ncbi.nlm.nih.gov/datasets/genome/GCA_028451465.1/)^38^. Syntenic blocks were completed using a custom script that filters Blastn matches for repeats, random matching, and multi-matching. Adjacent matches were then grouped based on alignment quality and density. Large draft scaffolds aligning to the same syntenic block were automatedly placed adjacently. Small draft scaffolds were placed and oriented manually, followed by the manual correction of small misassemblies. Next, we used the FISH-Karyotype of the naked mole-rat completed by Romanenko et al., 2023^39^ to address misassemblies and place centromeres. We re-aligned Hi-C sequencing data to these painted assemblies. Regions where syntenic alignments could not decide the orientation of a chromosomal segment were checked by visually inspecting the Hi-C contact map and manually “flipping” fragments if the opposite orientation creates a more consistent Hi-C contact map. These final assemblies were designated mHetGlaV3 and pHetGlaV3 (maternal/paternal, *Heterocephalus glaber*, Version 3). Annotations were lifted over from HetGlaV2 to HetGlaV3 using LiftOff/1.6.3^84^. Additionally, mHetGlaV2 assembly was included in Ensembl 111 (October 2023) as the NMR main build^17^. These gene symbols from Ensembl 111 were integrated into metGlaV3.

### Autosomal, sex chromosome, and mitochondrial assembly of HetGlaV4 (CAM-845F1)

We performed quality control on the raw read data using FastQC/0.12.1^126^ and NanoPlot/1.42.0^127^. To assess expected genome size and heterozygosity, we evaluated the k-mer profile of the PacBio HiFi reads using Jellyfish/2.3.1 ^128^ and GenomeScope/2.0^129^. We assembled PacBio HiFi and ONT ultra long reads using hifiasm/0.19.8^82^ in ultra long mode to generate a highly-contiguous primary assembly of NMR. After scaffolding, we aligned the assembly to mHetGlaV3 (GCA_944319715.1) using Minimap2/2.28^120^. Based on 1-to-1 alignments of scaffolds to chromosomes, we reordered, reoriented, and renamed scaffolds according to mHetGlaV3. Additionally, we assembled the mitogenome with MitoHiFi/3.2.2^130^ using the HiFi reads of *CAM-845F1* and the mitogenome NC_015112.1 as mapping reference.

### Sequence data processing with assembled genomes

#### Linked short-read sequencing

Reads were aligned to the NMR assemblies using the same alignment protocol as in the genome assembly. Briefly, reads were evaluated with fastqc^126^. Linked reads were then aligned and QC’d with the longranger pipeline. Genomes were indexed for longranger/2.2.2 with “longranger mkref” and aligned using “longranger align” with default parameters. This pipeline also generates alignment summaries https://github.com/10XGenomics/longranger.

#### Nanopore (R9.4) long-read sequencing

Fast5 files were basecalled using Guppy (v4.4.1) (https://community.nanoporetech.com) using the dna_r9.4.1_450bps_hac_prom.cfg configuration. FASTQ files were aligned to each assembly using minimap2^120^. Assemblies were indexed with minimap -d and then reads were aligned with minimap2 (-x map-ont -a -- secondary=no) and filtered with samtools view (-S -b -F 4 -F 0x800)^118,120^.

#### Hi-C sequencing

Paired-end Hi-C-seq reads were aligned to each assembly using juicer (2.0-encode-CPU)^117^. Briefly, reads were aligned to the bwa indexed assembly using the “juicer.sh” script with default parameters^117^. The generated *.sam file was then converted to the *.hic file using samtools view (-O SAM -F 1024), the sam_to_pre.awk script provided in the juicer package, and then juicer pre in the juicer tools (1.22.01) package^117,118^. Generated *.hic files were then visualized using Juicebox (1.11.08) and looping data was measured using HICCUPS within juicer tools (1.22.01)^36,117^.

#### ChIP- and ATAC-sequencing

Paired-end FASTQ files were initially visualized with fastqc^126^. TruSeq3-PE adaptors were trimmed with trimmomatic/0.32, and paired-end reads with a minimum read length of 36bp were kept^131^. Reads were aligned to HetGla1.2, mHetGlaV2, and mHetGlaV3 using the paired-end module of bwa-mem^13,116^. Aligned reads were converted to a sorted bam file with a q > 30 and with unique reads using samtools (1.9) view and samtools sort^118^. Then, mitochondrial reads were removed using samtools view^118^. These files were used for downstream analysis. As an additional level of QC, we removed duplicates using Picard (http://broadinstitute.github.io/picard/) with a lenient filter. Both sets of *.bam files were then evaluated with fastqc^126^. In the mouse, ChIP-seq sequencing reads were analyzed with the same parameters, with the additional step of filtering reads aligning to the ENCODE blacklist region (https://sites.google.com/site/anshulkundaje/projects/blacklists). For both the mouse and naked mole-rat Quality control of the datasets processed from the raw sequencing reads was assessed following the ENCODE ChIP-seq guidelines (Supplementary Table S3)^45^.

#### RNA sequencing

We re-processed publicly available full-length RNA-seq^50^ (Bens et al., 2018) to evaluate and annotate our NMR assemblies. These full-length RNA-seq data were the same datasets used to identify gene structures in our annotations^19,50^. Briefly, reads were downloaded from SRP061363 using stratoolkit/2.8.0 fastq-dump before Illumina adaptors were trimmed^132^. Unpaired reads were filtered and average base pair quality (q) < 30 using trim_galore/0.4.4^133^. Paired reads were aligned to each existing NMR assembly (Table 2) (HetGla1.2,HetGla1.7_hic_pac, mHetGlaV2, pHetGlaV2, mHetGlaV3, and pHetGlaV3)^13,15^ using STAR 2.7.10^134^ with the *.gtf of each associated assembly. PCR duplicates were then marked with Picard/1.133 (http://broadinstitute.github.io/picard/) using a lenient validation strategy, and aligned reads were again filtered using samtools view with a mapping quality q < 30^118^. These full-length RNA-seq data were processed using the same pipeline as in “Gene structure identification”.

**Table 2.**
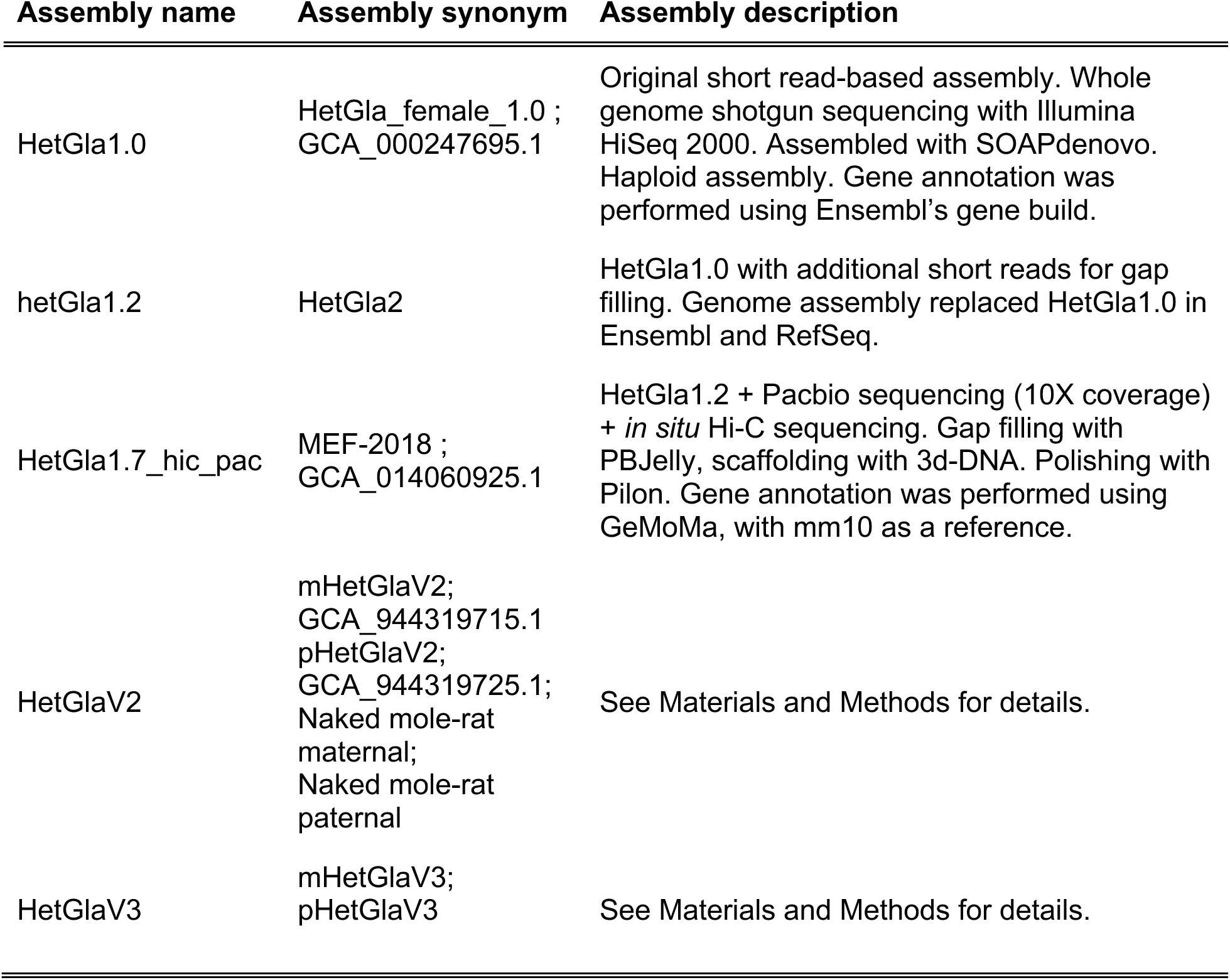
Summary of the complete genome assemblies compared in this study.

We measured basic assembly statistics and gene-level completeness of each assembly using QUAST/5.0 and BUSCO/5.2.2 (rodentia database), respectively^135,136^. We then measured genomic gaps using “bedtools nuc”^119^. We then completed a k-mer-based assembly evaluation of the mHetGlaV3 and pHetGlaV3 with Merqury (Meryl v1.4 release) using a database from linked reads from TOR-NMR2606 (maternal) and TOR-NMR2624 (paternal) NMR^32^. Next, we investigated full-length RNA-seq counting and gene detection differences in all assemblies (see Read alignment and processing for details).

### Single-nuclei RNA sequencing

We re-processed publicly available snRNA-seq data from the naked mole-rat hypothalamus (Faykoo-Martinez et al., in prep) previously generated in our lab. Briefly, the HetGla1.2 and mHetGlaV3 assemblies were indexed with cellranger (7.0.1) (https://github.com/10XGenomics/cellranger) mkref using the HetGla1.2 Ensembl^13^ gene build, and our liftOff-applied ^84^ annotation to mHetGlaV3 gtf files respectively. Reads were then aligned and quantified using cellranger count with default parameters. Aligned and quantified reads and cells were then normalized and clustered using “process_dgTMatrix_lists”, a wrapper for Seurat V5 and SCTransform using the scMappR package^137,138^. Cell-types were identified in Faykoo-Martinez et al., and maintained here. Briefly, they used a combination of reference markers and publicly available snRNA-seq datasets in the mouse^139–142^. Genes describing neuronal markers (e.g., CART neurons) reflect previously discovered cell-types in the mouse or NMR^143^. We then plotted the ratio of counts (RNA, SCT), features (RNA, SCT), and percent mitochondria adjusted for each cluster identified in Faykoo-Martinez et al., using a boxplot. We looked at differences in cell-type marker expression between assemblies for matching clusters by first taking the top 200 (by adjusted p-value, log2FC > 0) markers for the matched cell type in each assembly and comparing the fold-changes between the markers in each assembly. We identified genes with a >3 standard deviation difference in fold-change between assemblies, and we performed gene set enrichment of genes specific to mHetGlaV3 using gprofiler2 (using all cell-type markers from all cell-types as the statistical background)^144^. Markers between assemblies were plotted using ggplot2^144^. For snRNA-seq, we compared reads mapped to cells, total cells, genes identified per cell, cluster identification, and differences in marker genes per cluster between HetGla1.2 and mHetGlaV3.

### Naked mole-rat genome annotation

#### Repeat Region annotation

We annotated the repeat regions of mHetGlaV3, pHetGlaV3, and HetGlaV4 using “EarlGrey”^48^. We extracted the Kimura distances from the “prep_divFromAln” output of Earl Grey to annotate each TE found in “.filteredRepeats.bed” to its estimated evolutionary age^48,145,146^.

#### Gene structure annotation of mHetGlaV3 (TOR-NMR2626)

Our primary annotation of gene structures in the maternal and paternal assemblies was completed through submission and integration into Ensembl rapid release when gene annotations were completed using the RNA-seq pipeline within Ensembl gene build (https://rapid.ensembl.org/info/genome/genebuild/full_genebuild.html)^19^. These gene structures incorporated publicly available short-read-based RNA-seq data from Bens et al., 2018, which included the hypothalamus, pituitary, thyroid, ovary, skin, and kidney in subordinate and breeder NMR tissues^50^. We further evaluated and annotated gene structures using the “tool to infer orthologs from genome alignments” (TOGA/1.1.0), with the mm10 genome as a reference^41^. Briefly, we used the CACTUS aligner (version 2.5.1) to generate a .hal file between the mHetGlaV2 to mouse and pHetGlaV2 to mouse respectively^147^. Then, we used the hal2fasta, halStats, halLiftover, and finally, axtChain functions within CACTUS to convert the .hal file to a .chain file and genome .2bit files compatible with TOGA^41,147^.

#### Gene symbol annotation of mHetGlaV3 (TOR-NMR2626)

We integrated multiple approaches to call gene identities. Firstly, Ensembl called gene symbols on their annotations using the gene symbol transformer (GST) (https://github.com/Ensembl/gene_symbol_transformer)^19^. Briefly, the GST is a neural network transformer model trained on the protein sequences in the Ensembl Main Release, which learns higher-dimensional features of the CDS of each gene structure to predict the most likely gene symbol based on that gene structure. GST annotates gene structures with a confidence score >0.9 (range of 0-1). Secondly, we incorporated the gene symbols from the Ensembl 111 release, which incorporated the mHetGlaV2 into its stable release^17^. Thirdly, we generated a protein-level BLAST from UniProt^148^. Then, we input the translated CDS regions from the Rapid Ensembl into blastp (blast+/2.12.0)-evalue 1e-6 -max_hsps 6 -max_target_seqs 6 -outfmt 6)^148^. We identified the top protein IDs (UniProt IDs) and covered them with human gene symbols using Ensembl Biomart^148,149^. Similarly, we build a nucleotide blastdb from the RFam database^150^. Then, we aggregated Rapid-Ensembl gene structures annotated to “lncRNA”, “miRNA”, “scaRNA”, “snoRNA”, or “snRNA” into a fasta file of non-coding RNA’s. We annotated these regions’ non-coding RNAs within the RFam database using blastn, with the same cutoffs as for protein-coding genes, before mapping them to their RNA ID with Infernal^150–152^. Lastly, orthologous and paralogous regions identified by TOGA are mapped to mouse and human gene symbols^41^.

After gene symbols were called using each tool, we used a decision tree to identify the final gene symbol for each Ensembl gene structure. We used evidence from TOGA and Ensembl-111 (the stable release including mHetGlaV2) as the primary sources of evidence and gene symbols from Rapid-Ensembl, BLAST, and liftOff (from the annotation of HetGla1.2) approaches as secondary sources of evidence^17,19,41,84,148^. Specifically, if a gene had its symbol exclusively called by a primary evidence source, then the gene symbol would get the TOGA call. Similarly, if a gene symbol was called by TOGA and another method, then the gene was assigned the primary source annotation. If a gene symbol was called by one secondary annotation but no other tool (e.g., BLAST only), then the symbol was excluded. Afterwards, gene symbols were called with a majority voting method where a primary source was worth two votes, and a secondary source was worth one vote. Genes whose symbols did not have a clear majority vote (e.g., TOGA = Uncalled and Ensembl 111 = geneA, rapid ensembl=geneA, Ensembl-BLAST=geneB, HetGla1.2 LiftOver=geneC), then the gene region was flagged for manual inspection^17,19,41,84,148^. These ambiguous genes were almost always pseudonyms for the same gene (e.g., *Pilr-b* and. *Pilr-beta*). In these instances, we defaulted our gene symbols to the pseudonym found in the mouse genome, as it was our reference species in TOGA and downstream evolutionary analyses. If a gene symbol remained ambiguous, then we did not assign it a gene symbol. All gene symbol calls from each method and the consensus gene symbols were stored in Supplementary Tables S2 and S3.

#### Gene structure and symbol annotation of HetGlaV4 (CAM-845F1)

Gene models were transferred from the Ensembl 112 annotation of HetGlaV2 (Heterocephalus_glaber_female.Naked_mole-rat_maternal.112.gtf) using “liftoff -copy” to annotate any duplicated genes that were not assembled in HetGlaV2 or HetGlaV3^17,84^. Liftoff also transferred gene symbols from mHetGlaV3^17,84^.

We added additional gene structures to these annotations by performing TOGA using the mouse as a reference^41^, BRAKER3 in RNA and protein mode using publicly available RNA-seq data and protein sequences from the universal protein resource (UniProt)^50,85,148^, and *de novo* gene structures from the sample RNA-seq data using StringTie2 for model building and Mikado for model evaluation using TransDecoder (https://github.com/TransDecoder/TransDecoder) Portcullis, BLASTp, and the mikado mammalian scoring file for gene model evaluation^86,87,153,154^. These additional models were integrated into the gene models from Liftoff using Mikado in reference mode, thereby supplementing but not replacing gene models annotated by Ensembl 112^17,87^. Additional gene symbols were populated from alignments from TOGA and reciprocal alignments against the UniProt database^41,148,154^.

### Epigenetic annotations

#### Peak-calling and annotations

We called peaks for each antibody using MACS/2.1.1 (BAMPE)(150). ChIP-seq experiments used inputs within the same experimental batch as a reference for peak calling. Peaks were called from antibodies representing histone modifications (H3K4me3, H3K4me2, H3K27Ac, H3K27me3, H3K36me3, and H3K9me2) using the “--broad” parameter, while narrow peaks were called for CTCF and ATAC-seq experiments. Peaks were called for the ATAC-seq experiments without experimental inputs(150).

Genomic regions (e.g., peaks) were annotated to genomic features using the annotatePeak function “ChIPseeker” R package (v 1.34.1)(149). For the NMR, ChIPseeker-compatible genomic annotations were generated using applying makeTxDbFromGFF function in the GenomicFeatures package to the mHetGlaV3 annotations (see “Naked mole-rat genome annotations”) for details(152).

Genomic regions assigned to genes were then subject to gene-level functional enrichment To test for the enrichment of hypothalamus cell types within these gene lists, we applied the “tissue_scMappR_custom” to signature matrices generated from hypothalamic snRNA-seq data(134). In the NMR, we generated a signature matrix from the reprocessed snRNA-seq data in Faykoo-Martinez et al. (see Naked mole-rat snRNA-seq processing for details), using the “seurat_to_generes” and “generes_to_heatmap” functions in scMappR(134). Lastly, we identified enriched and de novo transcription-factor motifs of these genomic regions using HOMER (10-24-2019 release)(53,155).

#### Enhancer-gene mapping

We used the activity-by-contact (ABC) model to map genes to enhancers in the female subordinate NMR hypothalamus^77^ (https://github.com/broadinstitute/ABC-Enhancer-Gene-Prediction, version 2.0). First, we called ATAC-seq peaks in the female NMR subordinates with macs2 callpeaks (BAMPE)^77,155^. Peaks were then sorted and merged with samtools^77,118^. We then used the makeCandidateRegions.py^77^ with our annotations and “blacklist region” (i.e., assembly breakpoints between mHetGlaV2 and mHetGlaV3) (Supplementary File S3). Neighbourhoods of the top 100,000 peaks were measured using the run.neighborhoods.py using our H3K27Ac and ATAC-seq data, and the average gene expression from the full-length hypothalamus RNA-seq from Bens et al., 2018^50^ adding the --use_secondary_counting_method flag^77^. Then, the *.hic file in the NMR hypothalamus was converted into a format compatible with the ABC model using the juicebox_dump.py script (--resolution 5000 --include_raw)^77,156^. These files were all integrated to compute ABC enhancers using the predict.py script (--threshold 0.2 – scale_hic_using_powerlaw –make_all_putative)^77^.

#### Chromatin state mapping

We used ChromHMM/1.12^44^ to build chromatin state maps of the hypothalamus in mHetGlaV3. For the NMR, we inputted the .bam files for the H3K4me2, H3K4me3, H3K27Ac, H3K27me3, H3K36me3, H3K9me3, CTCF, ATAC-seq, and RNA-seq data into ChromHMM.jar BinarizeBam and LearnModel using default parameters^44^. We used these data to compute 20 chromatin states with a 200bp window. We manually annotated chromatin states following guidelines from the Encyclopedia of DNA Elements (ENCODE) project^45^.

### Naked mole-rat structural element mapping

#### Chromosome painting and comparison to other species and FISH-karyotype

The scaffolds are first assembled into pseudo-chromosomes with genome homology. The chromosome level assembly closest to *Heterocephalus glaber* is *Erethizon dorsatum* (GCA_028451465.1)^38^, with both belonging to infraorder *Hystricognathi*. The homology of both species to repeat-masked house mouse genome has been generated by BLASTN ^154^ (v2.14.1+), with parameters -evalue 1e-5 -outfmt 6. The results are analysed with SynBuild (v2019, https://github.com/igcbioinformatics/Syn2Chr), and manually adjusted when the paternal and maternal scaffolds conflict. 307 scaffolds, (2,460,490,714bp, 98.5% total length) from maternal assembly and 331 scaffolds, (2460249682bp, 98.5% total length) from paternal assembly have been assigned to *Heterocephalus glaber* pseudo chromosomes. Between scaffolds, 5000 bp ambiguous sites (N) were added.

#### Classification of tandem duplications

We computed segmental duplications within the soft-masked mHetGlaV3 and pHetGlaV3 using BISER, the successor to SeDef, with default parameters^157^. We filtered these segmental duplications to be within 100Kb of each other to select for potential tandem duplications. Then, we further filtered these tandem duplications such that each end of the duplication must overlap with a different annotated CDS from our gene annotation. Each candidate tandem duplication was evaluated manually by investigating syntenic blocks to the mouse mm10 genome and functional annotation of RNA-seq and epigenetic data. Candidate segmental duplications were manually annotated into whole-gene duplications, exon duplications, regulatory region duplications, gene-family expansions and paralogs, and false positives and processed for further filtering.

We defined gene, exon, and regulatory duplications as genes (or genomic regions) with multiple tandem copies in the NMR and for that same region to be expected to be a 1-1 ortholog between the mouse and other rodents (1-1 orthology key from Ensembl BioMart)^149^. We filtered BISER-detected^157^ self-alignment to different CDSs’ of gene clusters with variable copy numbers between species, as these regions reflect gene family expansions rather than random duplications (e.g., olfactory receptors, zinc finger TFs, keratin genes, histone complex genes etc.). We also filtered segmental duplication by BISER^157^ annotating to paralogs present in both the mouse and NMR. Lastly, we filtered possible false-positive tandem duplications, which we defined as tandem duplications in regions with a high rate of spurious alignment, either driven by low complexity or a potential assembly error. These regions were characterized by highly variable nanopore or ChIP-seq read depth, deviating from the assumption that the genome is covered uniformly. Identified genic, exonic, and regulatory duplications were visualized using RNA-seq, ChIP-seq, ATAC-seq data and with genome annotations (e.g., gene and repeat regions), and functional annotations (e.g., active enhancers) with the IGV^158^. They were also manually compared to the mm10 assembly using a dot plot^159^. We predicted the parental vs. child copy based on the location of each gene copy relative to mouse (mm10) and the number of exons vs. introns in each gene copy relative to mouse. If gene location and gene structure were ambiguous, we predicted parental vs. child gene copy using our RNA-seq, ChIP-seq, and ATAC-seq data, under the assumption that the child copy would be more likely to have a low or repressed signal than the parental copy.

#### Classification of candidate pseudogenes through duplication

We extracted Ensembl 111 one-to-many mapped genes between the mouse mm10 and mHetGlaV2 genomes using Ensembl BioMart to identify additional candidate pseudogenes to those classified by Ensembl^17,149^. We used mHetGlaV2 because this assembly was incorporated into the Ensembl 111 gene build. We then applied a cutoff of a 75% reciprocal alignment between overlapping regions in the mm10 and mHetGlaV2 genes. For each one-to-many mapped gene, we identified the “primary” gene with the largest length, gene order conservation, orthology confidence, and genome alignment coverage scores stored in BioMart^149^. From this pool, we identified gene copies with their “Gene order conversion score” < 50 and a gene length less than half the length of the primary gene copy as candidate pseudogenes. If the NMR gene was comparable in length to the mouse gene but was a low-confidence homolog, it was classified as “other.” Lastly, we tested whether these genes were expressed (>0.5 reads per kilobase per million expression in any tissue) to flag whether these candidate pseudogenes were processed. These annotations were then lifted over to mHetGlaV3.

#### Classification of gene inactivation and gene loss

We annotated protein-coding orthologs and paralogs of the mHetGlaV3 and pHetGlaV3 against the mm10 mouse genome assembly using TOGA with default parameters^41^. Briefly, TOGA classifies transcripts as “intact”, “paralogous projections”, “lost”, “partially intact”, and “missing” (i.e., a substantial proportion of the gene spans a genomic gap)^41^. “Lost” transcripts are identified as containing gene-inactivating mutations (frameshifting, stop codon or splice site mutations, exon or gene deletions)^41^. We extracted these inactivated regions and overlaid them with the full-length RNA-seq data. If an inactivated gene did not contain an expression level of 0.5 reads per kilobase per million (RPKM) in any available tissue, we considered it a candidate for gene loss. Gene-set enrichment of candidate-loss genes was completed with g:Profiler^144^. Gene family loss between the naked mole-rat and mouse were visualized using a linear plot between mHetGlaV3 and mm10 in the WashU Comparative Epigenome Browser. Regions with a missing gene despite an intact surrounding syntenic regions were considered candidates for gene loss^160^.

### Naked mole-rat genome evaluation

#### HetGlaV3 (TOR-NMR2626) genome evaluation

We measured basic assembly statistics and gene-level completeness of each assembly using QUAST/5.0 and BUSCO/5.2.2 (rodentia database), respectively^135,136^. We then measured genomic gaps using “bedtools nuc”^119^. We then completed a k-mer-based assembly evaluation of the mHetGlaV3 and pHetGlaV3 with Merqury (Meryl v1.4 release) using a database from linked reads from TOR-NMR2606 (maternal) and TOR-NMR2624 (paternal) NMR^32^. Next, we investigated full-length RNA-seq counting and gene detection differences in all assemblies (see Read alignment and processing for details). For snRNA-seq, we compared reads mapped to cells, total cells, genes identified per cell, cluster identification, and differences in marker genes per cluster between HetGla1.2 and mHetGlaV3.

We used minimap2 to align and compare assemblies within the NMR species^120^. We aligned assembly drafts using minimap2 (-cx asm5 --cs) to generate .paf files^120^. We visualized pairwise genome alignments using a genome-wide dot plot with D-GENIES^122^. Alignments and misassembly boundaries were identified by parsing the alignment “Association Table” outputted from D-Genies^122^. Regions of miscaffolding between the HetGlaV2 and HetGlaV3 assemblies (i.e., mHetGlaV2 vs. mHetGlaV3, pHetGlaV2 vs. pHetGlaV3) were annotated to the mHetGlaV3 genome using ChIPSeeker (v 1.34.1) and visualized against both to HetGlaV3 assemblies using the interactive genome viewer (IGV)^158,161^. For local alignments (e.g., visualization of an individual tandem duplication), we used pip-maker to generate dotplots between the naked mole-rat and mouse assemblies^159^.

#### HetGlaV4 (CAM-845F1) genome evaluation

We evaluated the resulting assembly by running quast/5.2.0^136^, assessed k-mer-based statistics with Merqury/1.3^32^, and evaluated the completeness of the gene space with BUSCO/5.7.1 ^135^. We then mapped Hi-C reads to the contig assembly using the Arima Genomics Hi-C mapping pipeline (https://github.com/ArimaGenomics/mapping_pipeline, commit b2b2457). We scaffolded the contigs using YaHS/1.2a ^162^ and converted name-sorted and duplicate marked BAM files with PretextMap/0.1.9 (https://github.com/sanger-tol/PretextMap) into a genome contact map to manually curate the assembly with PretextView/0.2.5 (https://github.com/sanger-tol/PretextView).

### Naked mole-rat cytogenetics

#### Development of naked mole-rat fibroblast cell-lines

Postnatal and adult subordinate NMRs were euthanized according to IACUC-approved methods and AVMA Guidelines for Euthanasia of Animals. A 2-3 cm2 skin biopsy harvested and transferred into NMR fibroblast medium: alphaMEM (Gibco™ 12571063) containing 15% FBS (Cytiva, SH3007103), 100 ug/ml VitaminC (Fujifilm Wako, 321-44823), 1x Glutamax (Gibco™ 35050061), 1 mM Sodium Pyruvate (Gibco™ 11360070), 1x non-essential amino acids (Gibco™ 11140050), 1x Penicillin/Streptomycin (Gibco™ 15140122) and 50 ug/ml Primocin (InvivoGen, ant-pm-05). Specimens were washed 3x in HBSS -/- containing 1x Penicillin/Streptomycin and 50 ugl/ml Primocin and the dermis layer was separated from the epidermis with Tweezers. Dermal sheets were washed in 1xHBSS -/- and incubated in 0.5% Collagenase B (Roche) in HBSS -/- with 3mM CaCl2 at 37 degrees with repeated pipetting with wide-bore pipette tips for 20-30 min when digestion was stopped with NMR fibroblast medium. After centrifugation cells and tissue pieces were seeded on Fibronectin- (42 ug/ml, FisherScientific, 356009) and Laminin-coated plates (21 ug/ml, Corning, CB-40239) and cultured at 32 degrees in 3% hypoxia in primary NMR fibroblast medium (50:50 alphaMEM with Ham’s-F12, 15% FBS, 100 ug/ml VitaminC, 1x Glutamax, 1 mM Sodium Pyruvate, 1x non-essential amino acids, 1x Penicillin/Streptomycin and 50 ug/ml Primocin) as passage 0. Cells were passaged with 0.05% trypsin and routinely checked for mycoplasma.

#### Cytogenetics of naked mole-rat cell-lines

Cultures were harvested following three hour mitotic arrest with 0.1 μg/mL Colcemid® and 30 minute addition of 10 μg/mL ethidium bromide using standard methodologies of hypotonic shock with 0.075 M KCl and fixation with methanol:acetic acid fixative (3:1)^163^. Metaphase preparations were made by dropping fixed cell suspension onto wet microscope slides, flooding with fixative and air-drying. Slides were aged at 90°C for 45 minutes and stained with the GTW banding method, G-banding with trypsin and Wright stain. G-banded slides were destained and subsequently stained with silver nitrate, AgNOR^164^. Metaphase cells were imaged and analyzed with a Zeiss Imager Z2 microscope (Zeiss Corp., Jena, Germany) with 100x planapochromatic objective and GenASIs™ imaging system (Applied Spectral Imaging Inc., Carlsbad, CA).

The 31 NMR chromosome classes were assigned numbers and arranged by sequence length and centromeric position. More than 350 metaphase cells were investigated. Sequence length was linked to in situ chromosome data by comparison of a G-banded metaphase image with whole-chromosome FISH sequentially performed on the same cell^39^. Morphologically similar chromosomes were grouped and given general descriptions. An ideogram was drawn for each chromosome class to illustrate centromeric position as well as landmark band positions and sizes. Example chromosome images demonstrate morphologies seen in each class, with resolutions ranging from approximately 300 to 500 bands per haploid cell.

## Acknowledgments

We would like to thank the genome centres who provided genome, RNA and epigenome sequencing: The Centre for Applied Genomics, Ontario Institute for Cancer Research, University of Washington Long Reads Sequencing Center. We would also like to thank Jane Reznick, Jay Zussman, and Phoebe Edwards for their helpful discussions and for sharing useful information while they worked with HetGlaV3, and Katherine M. Munson for sequencing of PacBio reads.

This work was supported by; a Natural Sciences and Engineering Research Council of Canada (NSERC) [RGPIN-2019–07014 to MDW; CIHR [202003PJT-437197 to MMH, MDW]; Canada First Research Excellence Fund (CFREF) Medicine by Design to MDW and TJP; NSERC CGS M, PGS D and Ontario Graduate Scholarships (to DJS) and the Ontario Genomics-CANSSI Ontario Postdoctoral Fellowship in Genome Data Science; NSERC PGS D, the association computing machinery special interest group on high performance computing (ACM/SIGHPC) Intel Computational and Data Science Fellowship (to MFM.); the Chan Zuckerberg Biohub (DJL), WM Keck Foundation (DJL); JTS is supported by the Ontario Institute for Cancer Research through funds provided by the Government of Ontario, the Government of Canada through Genome Canada and Ontario Genomics (OGI-136 and OGI-201) and the National Human Genome Research Institute (NHGRI 561 project 5R01HG009190); NSERC [RGPIN 2018–04780, RGPAS 2018–522465 to MMH; This work was supported, in part, by US National Institutes of Health (NIH) grant HG002385 to E.E.E; work was in part supported by a Dunhill Medical Trust grant RPGF2002\188 to EStJS. Research in the GB lab is supported by the UK Dementia Research Institute, which receives contributions from UK DRI, Ltd. and the UK MRC and by the Romanian Ministry of Research, Innovation and Digitization (no. PNRR-III-C9-2022-I8-66; contract 760114). Ensembl receives majority funding from Wellcome Trust [222155/Z/20/Z] with additional funding for specific project components. Ensembl receives further funding from The Biotechnology and Biological Sciences Research Council [BB/W019108/1, BB/T015608/1, BB/X018695/1]; UK Medical Research Council [MR/S000453/1]; Wellcome Trust [226458/Z/22/Z, 226083/Z/22/Z].

## Author contributions

MDW, GB, JTS, RB, EEE, MHM, DJL, TJP, ESt.JS obtained funding, designed the study, and supervised the work. DJS, MM, AN, MFM, KH, PZ, SM, SAA, JE, ALE, RK, JB, DY, NK, KU, FM, TH, HH, ZC, HELL, DVL, YJ, JL performed the research and analysed the data. DS, MDW wrote the manuscript. All authors read and reviewed the manuscript.

## Data availability

1. Genome assemblies can be found at ENA (mHetGlaV2(https://www.ebi.ac.uk/ena/browser/view/GCA_944319715.1), pHetGlaV2(https://www.ebi.ac.uk/ena/browser/view/GCA_944319725.1), mHetGlaV3 (https://www.ebi.ac.uk/ena/browser/view/GCA_964261345.1), pHetGlaV3 (https://www.ebi.ac.uk/ena/browser/view/GCA_964261355.1)). At the time of submission, mHetGlaV2 is also the current naked mole-rat reference genome on Ensembl (https://useast.ensembl.org/Heterocephalus_glaber_female/Info/Index).
2. Gene, repeat, and epigenome annotations aligned to mHetGlaV3 are stored on a Zenodo repository (https://zenodo.org/records/14194953).
3. Oxford nanopore long reads (R9.4) (TOR-NMR2626) (ERX9397318), 10X-linked reads (TOR-NMR 2626, TOR-NMR2606, TOR-NMR2624, TOR-NMR2625) (PRJEB53592), and Hi-C sequencing (PRJEB53711), data used in mHetGlaV3 assembly were stored in ENA.
4. ChIP-seq (E-MTAB-14648) and ATAC-seq (E-MTAB-14649) data are stored in array express and will be released with the completion of the CAM-845F1 (HetGlaV4) assembly. For data inquiries please contact Dustin Sokolowski, Michael Wilson, and Gabriel Balmus.

## Bibliography

1. Pamenter, M. E. Adaptations to a hypoxic lifestyle in naked mole-rats. J. Exp. Biol. 225, (2022).

2. Buffenstein, R. et al. The naked truth: a comprehensive clarification and classification of current “myths” in naked mole-rat biology. Biol. Rev. Camb. Philos. Soc. 97, 115–140 (2022).

3. Jarvis, J. U., O’Riain, M. J., Bennett, N. C. & Sherman, P. W. Mammalian eusociality: a family affair. Trends Ecol. Evol. 9, 47–51 (1994).

4. Keane, M. et al. The Naked Mole Rat Genome Resource: facilitating analyses of cancer and longevity-related adaptations. Bioinformatics 30, 3558–3560 (2014).

5. Brieño-Enríquez, M. A. et al. Postnatal oogenesis leads to an exceptionally large ovarian reserve in naked mole-rats. Nat. Commun. 14, 670 (2023).

6. Holmes, M. M. et al. Social control of brain morphology in a eusocial mammal. Proc Natl Acad Sci USA 104, 10548–10552 (2007).

7. Buffenstein, R. & Jarvis, J. U. M. The naked mole rat--a new record for the oldest living rodent. Sci. Aging Knowledge Environ. 2002, pe7 (2002).

8. Amoroso, V. G., Zhao, A., Vargas, I. & Park, T. J. Naked Mole-Rats Demonstrate Profound Tolerance to Low Oxygen, High Carbon Dioxide, and Chemical Pain. Animals (Basel) 13, (2023).

9. Tian, X. et al. High-molecular-mass hyaluronan mediates the cancer resistance of the naked mole rat. Nature 499, 346–349 (2013).

10. Azpurua, J. et al. Naked mole-rat has increased translational fidelity compared with the mouse, as well as a unique 28S ribosomal RNA cleavage. Proc Natl Acad Sci USA 110, 17350–17355 (2013).

11. Ferris, E., Abegglen, L. M., Schiffman, J. D. & Gregg, C. Accelerated evolution in distinctive species reveals candidate elements for clinically relevant traits, including mutation and cancer resistance. Cell Rep. 22, 2742–2755 (2018).

12. Lewis, K. N. et al. Unraveling the message: insights into comparative genomics of the naked mole-rat. Mamm. Genome 27, 259–278 (2016).

13. Kim, E. B. et al. Genome sequencing reveals insights into physiology and longevity of the naked mole rat. Nature 479, 223–227 (2011).

14. Fang, X. et al. Adaptations to a subterranean environment and longevity revealed by the analysis of mole rat genomes. Cell Rep. 8, 1354–1364 (2014).

15. Zhou, X. et al. Beaver and naked mole rat genomes reveal common paths to longevity. Cell Rep. 32, 107949 (2020).

16. Lok, S. et al. De Novo Genome and Transcriptome Assembly of the Canadian Beaver (Castor canadensis). G3 (Bethesda) 7, 755–773 (2017).

17. Martin, F. J. et al. Ensembl 2023. Nucleic Acids Res. 51, D933–D941 (2023).

18. O’Leary, N. A. et al. Reference sequence (RefSeq) database at NCBI: current status, taxonomic expansion, and functional annotation. Nucleic Acids Res. 44, D733–45 (2016).

19. Cunningham, F. et al. Ensembl 2022. Nucleic Acids Res. 50, D988–D995 (2022).

20. Christmas, M. J. et al. Evolutionary constraint and innovation across hundreds of placental mammals. Science 380, eabn3943 (2023).

21. Dengler-Crish, C. M. & Catania, K. C. Cessation of reproduction-related spine elongation after multiple breeding cycles in female naked mole-rats. Anat Rec (Hoboken) 292, 131–137 (2009).

22. Greenhalgh, R. et al. Trio-binned genomes of the woodrats Neotoma bryanti and Neotoma lepida reveal novel gene islands and rapid copy number evolution of xenobiotic metabolizing genes. Mol. Ecol. Resour 22, 2713–2731 (2022).

23. Flynn, J. M., Ahmed-Braimah, Y. H., Long, M., Wing, R. A. & Clark, A. G. High quality genome assemblies reveal evolutionary dynamics of repetitive DNA and structural rearrangements in the Drosophila virilis sub-group. BioRxiv (2023) doi:10.1101/2023.08.13.553086.

24. Marx, V. Method of the year: long-read sequencing. Nat. Methods 20, 6–11 (2023).

25. M Real, F., et al. The mole genome reveals regulatory rearrangements associated with adaptive intersexuality. Science 370, 208–214 (2020).

26. Albertin, C. B. et al. The octopus genome and the evolution of cephalopod neural and morphological novelties. Nature 524, 220–224 (2015).

27. Sulak, M. et al. TP53 copy number expansion is associated with the evolution of increased body size and an enhanced DNA damage response in elephants. eLife 5, (2016).

28. Booker, B. M. et al. Bat accelerated regions identify a bat forelimb specific enhancer in the hoxd locus. PLoS Genet. 12, e1005738 (2016).

29. Eckalbar, W. L. et al. Transcriptomic and epigenomic characterization of the developing bat wing. Nat. Genet. 48, 528–536 (2016).

30. Formenti, G. et al. Complete vertebrate mitogenomes reveal widespread repeats and gene duplications. Genome Biol. 22, 120 (2021).

31. Koren, S. et al. De novo assembly of haplotype-resolved genomes with trio binning. Nat. Biotechnol. (2018) doi:10.1038/nbt.4277.

32. Rhie, A., Walenz, B. P., Koren, S. & Phillippy, A. M. Merqury: reference-free quality, completeness, and phasing assessment for genome assemblies. Genome Biol. 21, 245 (2020).

33. Vaser, R., Sović, I., Nagarajan, N. & Šikić, M. Fast and accurate *de novo* genome assembly from long uncorrected reads. Genome Res. 27, 737–746 (2017).

34. Lee, J. Y. et al. Comparative evaluation of Nanopore polishing tools for microbial genome assembly and polishing strategies for downstream analysis. Sci. Rep. 11, 20740 (2021).

35. Ghurye, J. et al. Integrating Hi-C links with assembly graphs for chromosome-scale assembly. PLoS Comput. Biol. 15, e1007273 (2019).

36. Dudchenko, O., et al. The Juicebox Assembly Tools module facilitates *de novo* assembly of mammalian genomes with chromosome-length scaffolds for under $1000. BioRxiv (2018) doi:10.1101/254797.

37. Garrison, E. & Marth, G. Haplotype-based variant detection from short-read sequencing. arXiv (2012).

38. Rhie, A. et al. Towards complete and error-free genome assemblies of all vertebrate species. Nature 592, 737–746 (2021).

39. Romanenko, S. A. et al. Integration of fluorescence in situ hybridization and chromosome-length genome assemblies revealed synteny map for guinea pig, naked mole-rat, and human. Sci. Rep. 13, 21055 (2023).

40. Nurk, S. et al. The complete sequence of a human genome. Science 376, 44–53 (2022).

41. Kirilenko, B. M. et al. Integrating gene annotation with orthology inference at scale. Science 380, eabn3107 (2023).

42. Parey, E. et al. Phylogenetic modeling of enhancer shifts in African mole-rats reveals regulatory changes associated with tissue-specific traits. Genome Res. 33, 1513–1526 (2023).

43. Ballinger, M. A., Mack, K. L., Durkin, S. M., Riddell, E. A. & Nachman, M. W. Environmentally robust cis-regulatory changes underlie rapid climatic adaptation. Proc Natl Acad Sci USA 120, e2214614120 (2023).

44. Ernst, J. & Kellis, M. ChromHMM: automating chromatin-state discovery and characterization. Nat. Methods 9, 215–216 (2012).

45. Landt, S. G. et al. ChIP-seq guidelines and practices of the ENCODE and modENCODE consortia. Genome Res. 22, 1813–1831 (2012).

46. Hoffman, M. M. et al. Integrative annotation of chromatin elements from ENCODE data. Nucleic Acids Res. 41, 827–841 (2013).

47. Jangam, D., Feschotte, C. & Betrán, E. Transposable element domestication as an adaptation to evolutionary conflicts. Trends Genet. 33, 817–831 (2017).

48. Baril, T., Galbraith, J. & Hayward, A. Earl Grey: A Fully Automated User-Friendly Transposable Element Annotation and Analysis Pipeline. Mol. Biol. Evol. 41, (2024).

49. Platt, R. N., Blanco-Berdugo, L. & Ray, D. A. Accurate transposable element annotation is vital when analyzing new genome assemblies. Genome Biol. Evol. 8, 403–410 (2016).

50. Bens, M. et al. Naked mole-rat transcriptome signatures of socially suppressed sexual maturation and links of reproduction to aging. BMC Biol. 16, 77 (2018).

51. Zheng, Z., Hua, R., Xu, G., Yang, H. & Shi, P. Gene losses may contribute to subterranean adaptations in naked mole-rat and blind mole-rat. BMC Biol. 20, 44 (2022).

52. van der Horst, G., Maree, L., Kotzé, S. H. & O’Riain, M. J. Sperm structure and motility in the eusocial naked mole-rat, Heterocephalus glaber: a case of degenerative orthogenesis in the absence of sperm competition? BMC Evol. Biol. 11, 351 (2011).

53. van der Horst, G., Kotzé, S. H., O’Riain, M. J. & Maree, L. Testicular Structure and Spermatogenesis in the Naked Mole-Rat Is Unique (Degenerate) and Atypical Compared to Other Mammals. Front. Cell Dev. Biol. 7, 234 (2019).

54. Firman, R. C. Of mice and women: advances in mammalian sperm competition with a focus on the female perspective. Philos. Trans. R. Soc. Lond. B Biol. Sci. 375, 20200082 (2020).

55. Xu, K., Yang, L., Zhang, L. & Qi, H. Lack of AKAP3 disrupts integrity of the subcellular structure and proteome of mouse sperm and causes male sterility. Development 147, (2020).

56. Li, J. et al. The immunity-related GTPase IRGC mediates interaction between lipid droplets and mitochondria to facilitate sperm motility. FEBS Lett. 597, 1595–1605 (2023).

57. Kaneda, Y. et al. IRGC1, a testis-enriched immunity related GTPase, is important for fibrous sheath integrity and sperm motility in mice. Dev. Biol. 488, 104–113 (2022).

58. Ren, D. et al. A sperm ion channel required for sperm motility and male fertility. Nature 413, 603–609 (2001).

59. Jimenez, T., McDermott, J. P., Sánchez, G. & Blanco, G. Na,K-ATPase alpha4 isoform is essential for sperm fertility. Proc Natl Acad Sci USA 108, 644–649 (2011).

60. Zhou, L. et al. Structures of sperm flagellar doublet microtubules expand the genetic spectrum of male infertility. Cell 186, 2897–2910.e19 (2023).

61. Li, Y.-F. et al. FSCB, a novel protein kinase A-phosphorylated calcium-binding protein, is a CABYR-binding partner involved in late steps of fibrous sheath biogenesis. J. Biol. Chem. 282, 34104–34119 (2007).

62. Zhang, X. et al. FSCB phosphorylation regulates mouse spermatozoa capacitation through suppressing SUMOylation of ROPN1/ROPN1L. Am. J. Transl. Res. 8, 2776–2782 (2016).

63. Luo, C. et al. Meiotic chromatin-associated HSF5 is indispensable for pachynema progression and male fertility. Nucleic Acids Res. 52, 10255–10275 (2024).

64. Jumeau, F. et al. Defining the human sperm microtubulome: an integrated genomics approach. Biol. Reprod. 96, 93–106 (2017).

65. Groza, T. et al. The International Mouse Phenotyping Consortium: comprehensive knockout phenotyping underpinning the study of human disease. Nucleic Acids Res. 51, D1038–D1045 (2023).

66. Tardif, S. et al. Zonadhesin is essential for species specificity of sperm adhesion to the egg zona pellucida. J. Biol. Chem. 285, 24863–24870 (2010).

67. Hilton, H. G. et al. Single-cell transcriptomics of the naked mole-rat reveals unexpected features of mammalian immunity. PLoS Biol. 17, e3000528 (2019).

68. Wilson, M. D., Cheung, J., Martindale, D. W., Scherer, S. W. & Koop, B. F. Comparative analysis of the paired immunoglobulin-like receptor (PILR) locus in six mammalian genomes: duplication, conversion, and the birth of new genes. Physiol. Genomics 27, 201–218 (2006).

69. Shiratori, I., Ogasawara, K., Saito, T., Lanier, L. L. & Arase, H. Activation of natural killer cells and dendritic cells upon recognition of a novel CD99-like ligand by paired immunoglobulin-like type 2 receptor. J. Exp. Med. 199, 525–533 (2004).

70. Sharma, V. et al. A genomics approach reveals insights into the importance of gene losses for mammalian adaptations. Nat. Commun. 9, 1215 (2018).

71. Chaisson, M. J. P., Sulovari, A., Valdmanis, P. N., Miller, D. E. & Eichler, E. E. Advances in the discovery and analyses of human tandem repeats. Emerg. Top. Life Sci. 7, 361–381 (2023).

72. MacRae, S. L. et al. Comparative analysis of genome maintenance genes in naked mole rat, mouse, and human. Aging Cell 14, 288–291 (2015).

73. Augereau, A. et al. Naked mole rat TRF1 safeguards glycolytic capacity and telomere replication under low oxygen. Sci. Adv. 7, (2021).

74. Tang, N., Cai, X., Peng, L., Liu, H. & Chen, Y. TCP1 regulates Wnt7b/β-catenin pathway through P53 to influence the proliferation and migration of hepatocellular carcinoma cells. Signal Transduct. Target. Ther. 5, 169 (2020).

75. Chang, Y.-X. et al. Chaperonin-Containing TCP-1 Promotes Cancer Chemoresistance and Metastasis through the AKT-GSK3β-β-Catenin and XIAP-Survivin Pathways. Cancers (Basel) 12, (2020).

76. Zhen, D., Liu, J., Zhang, X. D. & Song, Z. Kynurenic acid acts as a signaling molecule regulating energy expenditure and is closely associated with metabolic diseases. Front Endocrinol (Lausanne) 13, 847611 (2022).

77. Fulco, C. P. et al. Activity-by-contact model of enhancer-promoter regulation from thousands of CRISPR perturbations. Nat. Genet. 51, 1664–1669 (2019).

78. Kawamura, Y. et al. Cellular senescence induction leads to progressive cell death via the INK4a-RB pathway in naked mole-rats. EMBO J. 42, e111133 (2023).

79. Hon, T. et al. Highly accurate long-read HiFi sequencing data for five complex genomes. Sci. Data 7, 399 (2020).

80. Jain, M. et al. Nanopore sequencing and assembly of a human genome with ultra-long reads. Nat. Biotechnol. 36, 338–345 (2018).

81. Sereika, M. et al. Oxford Nanopore R10.4 long-read sequencing enables the generation of near-finished bacterial genomes from pure cultures and metagenomes without short-read or reference polishing. Nat. Methods 19, 823–826 (2022).

82. Cheng, H., Concepcion, G. T., Feng, X., Zhang, H. & Li, H. Haplotype-resolved de novo assembly using phased assembly graphs with hifiasm. Nat. Methods 18, 170–175 (2021).

83. Adams, D., et al. The structural diversity of telomeres and centromeres across mouse subspecies revealed by complete assemblies. BioRxiv (2024) doi:10.1101/2024.10.24.619615.

84. Shumate, A. & Salzberg, S. L. Liftoff: accurate mapping of gene annotations. Bioinformatics 37, 1639–1643 (2021).

85. Gabriel, L., et al. BRAKER3: Fully automated genome annotation using RNA-seq and protein evidence with GeneMark-ETP, AUGUSTUS and TSEBRA. BioRxiv (2024) doi:10.1101/2023.06.10.544449.

86. Kovaka, S. et al. Transcriptome assembly from long-read RNA-seq alignments with StringTie2. Genome Biol. 20, 278 (2019).

87. Venturini, L., Caim, S., Kaithakottil, G. G., Mapleson, D. L. & Swarbreck, D. Leveraging multiple transcriptome assembly methods for improved gene structure annotation. Gigascience 7, (2018).

88. Pasupuleti, V. et al. Dysregulated D-dopachrome tautomerase, a hypoxia-inducible factor-dependent gene, cooperates with macrophage migration inhibitory factor in renal tumorigenesis. J. Biol. Chem. 289, 3713–3723 (2014).

89. Maruggi, M., Nguyen, A. T. & Resta, S. Oxidative stress inhibits Klotho activation of TRPV5 and TRPV6 in intestinal epithelial cells. FASEB J. 24, (2010).

90. Song, E. et al. Holo-lipocalin-2-derived siderophores increase mitochondrial ROS and impair oxidative phosphorylation in rat cardiomyocytes. Proc Natl Acad Sci USA 115, 1576–1581 (2018).

91. Huang, S., Howington, M. B., Dobry, C. J., Evans, C. R. & Leiser, S. F. Flavin-Containing Monooxygenases Are Conserved Regulators of Stress Resistance and Metabolism. Front. Cell Dev. Biol. 9, 630188 (2021).

92. Bime, C. et al. Reactive oxygen species-associated molecular signature predicts survival in patients with sepsis. Pulm. Circ. 6, 196–201 (2016).

93. Li, L.-J. et al. Chaperonin containing TCP-1 subunit 3 is critical for gastric cancer growth. Oncotarget 8, 111470–111481 (2017).

94. Yadav, S., Ali, V., Singh, Y., Kanojia, S. & Goyal, N. Leishmania donovani chaperonin TCP1γ subunit protects miltefosine induced oxidative damage. Int. J. Biol. Macromol. 165, 2607–2620 (2020).

95. Mazurais, D. et al. Identification of hypoxia-regulated genes in the liver of common sole (Solea solea) fed different dietary lipid contents. Mar. Biotechnol. 16, 277–288 (2014).

96. Park, S. H. et al. Activating CCT2 triggers Gli-1 activation during hypoxic condition in colorectal cancer. Oncogene 39, 136–150 (2020).

97. Zhang, F. et al. Nudix Hydrolase NUDT16 Regulates 53BP1 Protein by Reversing 53BP1 ADP-Ribosylation. Cancer Res. 80, 999–1010 (2020).

98. Fjeld, K. et al. Characterization of CEL-DUP2: Complete duplication of the carboxyl ester lipase gene is unlikely to influence risk of chronic pancreatitis. Pancreatology 20, 377–384 (2020).

99. Johansson, B. B. et al. The role of the carboxyl ester lipase (CEL) gene in pancreatic disease. Pancreatology 18, 12–19 (2018).

100. Ræder, H. et al. Carboxyl-ester lipase maturity-onset diabetes of the young is associated with development of pancreatic cysts and upregulated MAPK signaling in secretin-stimulated duodenal fluid. Diabetes 63, 259–269 (2014).

101. Freire Jorge, P., Goodwin, M. L., Renes, M. H., Nijsten, M. W. & Pamenter, M. Low Cancer Incidence in Naked Mole-Rats May Be Related to Their Inability to Express the Warburg Effect. Front. Physiol. 13, 859820 (2022).

102. Munro, D., Baldy, C., Pamenter, M. E. & Treberg, J. R. The exceptional longevity of the naked mole-rat may be explained by mitochondrial antioxidant defenses. Aging Cell 18, e12916 (2019).

103. Tian, X. et al. INK4 locus of the tumor-resistant rodent, the naked mole rat, expresses a functional p15/p16 hybrid isoform. Proc Natl Acad Sci USA 112, 1053–1058 (2015).

104. Zhang, Z. et al. Increased hyaluronan by naked mole-rat Has2 improves healthspan in mice. Nature 621, 196–205 (2023).

105. Hadj-Moussa, H., Eaton, L., Cheng, H., Pamenter, M. E. & Storey, K. B. Naked mole-rats resist the accumulation of hypoxia-induced oxidative damage. Comp Biochem Physiol, Part A Mol Integr Physiol 273, 111282 (2022).

106. Seluanov, A., Gladyshev, V. N., Vijg, J. & Gorbunova, V. Mechanisms of cancer resistance in long-lived mammals. Nat. Rev. Cancer 18, 433–441 (2018).

107. Schmidt, H. et al. Hypoxia tolerance, longevity and cancer-resistance in the mole rat Spalax - a liver transcriptomics approach. Sci. Rep. 7, 14348 (2017).

108. Sharon, D., Tilgner, H., Grubert, F. & Snyder, M. A single-molecule long-read survey of the human transcriptome. Nat. Biotechnol. 31, 1009–1014 (2013).

109. Ballester, B. et al. Multi-species, multi-transcription factor binding highlights conserved control of tissue-specific biological pathways. eLife 3, e02626 (2014).

110. Herrero, J. et al. Ensembl comparative genomics resources. Database (Oxford) 2016, (2016).

111. Logsdon, G. A. et al. The variation and evolution of complete human centromeres. Nature 629, 136–145 (2024).

112. National Research Council (US) Committee for the Update of the Guide for the Care and Use of Laboratory Animals. Guide for the care and use of laboratory animals. (National Academies Press (US), 2011). doi:10.17226/12910.

113. Buenrostro, J. D., Wu, B., Chang, H. Y. & Greenleaf, W. J. ATAC-seq: a method for assaying chromatin accessibility genome-wide. Curr. Protoc. Mol. Biol. 109, 21.29.1–21.29.9 (2015).

114. Dudchenko, O. et al. De novo assembly of the Aedes aegypti genome using Hi-C yields chromosome-length scaffolds. Science 356, 92–95 (2017).

115. Kolmogorov, M., Yuan, J., Lin, Y. & Pevzner, P. A. Assembly of long, error-prone reads using repeat graphs. Nat. Biotechnol. 37, 540–546 (2019).

116. Li, H. Aligning sequence reads, clone sequences and assembly contigs with BWA-MEM. arXiv (2013).

117. Durand, N. C. et al. Juicer Provides a One-Click System for Analyzing Loop-Resolution Hi-C Experiments. Cell Syst. 3, 95–98 (2016).

118. Li, H. et al. The Sequence Alignment/Map format and SAMtools. Bioinformatics 25, 2078–2079 (2009).

119. Quinlan, A. R. & Hall, I. M. BEDTools: a flexible suite of utilities for comparing genomic features. Bioinformatics 26, 841–842 (2010).

120. Li, H. Minimap2: pairwise alignment for nucleotide sequences. Bioinformatics 34, 3094–3100 (2018).

121. Danecek, P. et al. Twelve years of SAMtools and BCFtools. Gigascience 10, (2021).

122. Cabanettes, F. & Klopp, C. D-GENIES: dot plot large genomes in an interactive, efficient and simple way. PeerJ 6, e4958 (2018).

123. Lowe, T. M. & Eddy, S. R. tRNAscan-SE: a program for improved detection of transfer RNA genes in genomic sequence. Nucleic Acids Res. 25, 955–964 (1997).

124. Langmead, B. & Salzberg, S. L. Fast gapped-read alignment with Bowtie 2. Nat. Methods 9, 357–359 (2012).

125. Love, M. I., Huber, W. & Anders, S. Moderated estimation of fold change and dispersion for RNA-seq data with DESeq2. Genome Biol. 15, 550 (2014).

126. Brown, J., Pirrung, M. & McCue, L. A. FQC Dashboard: integrates FastQC results into a web-based, interactive, and extensible FASTQ quality control tool. Bioinformatics 33, 3137–3139 (2017).

127. De Coster, W. & Rademakers, R. NanoPack2: population-scale evaluation of long-read sequencing data. Bioinformatics 39, (2023).

128. Marçais, G. & Kingsford, C. A fast, lock-free approach for efficient parallel counting of occurrences of k-mers. Bioinformatics 27, 764–770 (2011).

129. Ranallo-Benavidez, T. R., Jaron, K. S. & Schatz, M. C. GenomeScope 2.0 and Smudgeplot for reference-free profiling of polyploid genomes. Nat. Commun. 11, 1432 (2020).

130. Uliano-Silva, M. et al. MitoHiFi: a python pipeline for mitochondrial genome assembly from PacBio high fidelity reads. BMC Bioinformatics 24, 288 (2023).

131. Bolger, A. M., Lohse, M. & Usadel, B. Trimmomatic: A flexible trimmer for Illumina sequence data. Bioinformatics 30, 2114–2120 (2014).

132. Leinonen, R., Sugawara, H., Shumway, M. & International Nucleotide Sequence Database Collaboration. The sequence read archive. Nucleic Acids Res. 39, D19–21 (2011).

133. Martin, M. Cutadapt removes adapter sequences from high-throughput sequencing reads. EMBnet j. 17, 10 (2011).

134. Dobin, A. et al. STAR: ultrafast universal RNA-seq aligner. Bioinformatics 29, 15–21 (2013).

135. Simão, F. A., Waterhouse, R. M., Ioannidis, P., Kriventseva, E. V. & Zdobnov, E. M. BUSCO: assessing genome assembly and annotation completeness with single-copy orthologs. Bioinformatics 31, 3210–3212 (2015).

136. Gurevich, A., Saveliev, V., Vyahhi, N. & Tesler, G. QUAST: quality assessment tool for genome assemblies. Bioinformatics 29, 1072–1075 (2013).

137. Hao, Y. et al. Dictionary learning for integrative, multimodal and scalable single-cell analysis. Nat. Biotechnol. 42, 293–304 (2024).

138. Sokolowski, D. J., et al. Single-cell mapper (scMappR): using scRNA-seq to infer the cell-type specificities of differentially expressed genes. NAR Genom. Bioinform. 3, lqab011 (2021).

139. Hajdarovic, K. H. et al. Single-cell analysis of the aging female mouse hypothalamus. Nat. Aging 2, 662–678 (2022).

140. Morris, M. E. et al. A single-cell atlas of the cycling murine ovary. eLife 11, (2022).

141. Zhang, X. et al. CellMarker: a manually curated resource of cell markers in human and mouse. Nucleic Acids Res. 47, D721–D728 (2019).

142. Diaz-Mejia, J. J. et al. Evaluation of methods to assign cell type labels to cell clusters from single-cell RNA-sequencing data [version 3; peer review: 2 approved, 1 approved with reservations]. F1000Res. 8, 296 (2019).

143. Lau, J. et al. CART neurons in the arcuate nucleus and lateral hypothalamic area exert differential controls on energy homeostasis. Mol. Metab. 7, 102–118 (2018).

144. Kolberg, L., Raudvere, U., Kuzmin, I., Vilo, J. & Peterson, H. gprofiler2 -- an R package for gene list functional enrichment analysis and namespace conversion toolset g:Profiler. F1000Res. 9, (2020).

145. Kimura, M. A simple method for estimating evolutionary rates of base substitutions through comparative studies of nucleotide sequences. J. Mol. Evol. 16, 111–120 (1980).

146. Chalopin, D., Naville, M., Plard, F., Galiana, D. & Volff, J.-N. Comparative analysis of transposable elements highlights mobilome diversity and evolution in vertebrates. Genome Biol. Evol. 7, 567–580 (2015).

147. Armstrong, J. et al. Progressive Cactus is a multiple-genome aligner for the thousand-genome era. Nature 587, 246–251 (2020).

148. UniProt Consortium. The universal protein resource (uniprot) in 2010. Nucleic Acids Res. 38, D142-8 (2010).

149. Kinsella, R. J. et al. Ensembl BioMarts: a hub for data retrieval across taxonomic space. Database (Oxford) 2011, bar030 (2011).

150. Griffiths-Jones, S., Bateman, A., Marshall, M., Khanna, A. & Eddy, S. R. Rfam: an RNA family database. Nucleic Acids Res. 31, 439–441 (2003).

151. Nawrocki, E. P., Kolbe, D. L. & Eddy, S. R. Infernal 1.0: inference of RNA alignments. Bioinformatics 25, 1335–1337 (2009).

152. Nawrocki, E. P. Annotating functional RNAs in genomes using Infernal. Methods Mol. Biol. 1097, 163–197 (2014).

153. Mapleson, D., Venturini, L., Kaithakottil, G. & Swarbreck, D. Efficient and accurate detection of splice junctions from RNA-seq with Portcullis. Gigascience 7, giy131 (2018).

154. Müller, A., MacCallum, R. M. & Sternberg, M. J. Benchmarking PSI-BLAST in genome annotation. J. Mol. Biol. 293, 1257–1271 (1999).

155. Zhang, Y. et al. Model-based analysis of ChIP-Seq (MACS). Genome Biol. 9, R137 (2008).

156. Robinson, J. T. et al. Juicebox.js Provides a Cloud-Based Visualization System for Hi-C Data. Cell Syst. 6, 256–258.e1 (2018).

157. Išerić, H., Alkan, C., Hach, F. & Numanagić, I. Fast characterization of segmental duplication structure in multiple genome assemblies. Algorithms Mol. Biol. 17, 4 (2022).

158. Thorvaldsdóttir, H., Robinson, J. T. & Mesirov, J. P. Integrative Genomics Viewer (IGV): high-performance genomics data visualization and exploration. Brief. Bioinformatics 14, 178–192 (2013).

159. Schwartz, S. et al. PipMaker--a web server for aligning two genomic DNA sequences. Genome Res. 10, 577–586 (2000).

160. Zhuo, X. et al. Comparing genomic and epigenomic features across species using the WashU Comparative Epigenome Browser. Genome Res. 33, 824–835 (2023).

161. Yu, G., Wang, L.-G. & He, Q.-Y. ChIPseeker: an R/Bioconductor package for ChIP peak annotation, comparison and visualization. Bioinformatics 31, 2382–2383 (2015).

162. Zhou, C., McCarthy, S. A. & Durbin, R. YaHS: yet another Hi-C scaffolding tool. Bioinformatics 39, (2023).

163. Houck, M. L., Lear, T. L. & Charter, S. J. Animal cytogenetics. in The AGT cytogenetics laboratory manual (eds. Arsham, M. S., Barch, M. J. & Lawce, H. J.) 1055–1102 (Wiley, 2017). doi:10.1002/9781119061199.ch24.

164. Lawce, H. J. Chromosome stains. in The AGT cytogenetics laboratory manual (eds. Arsham, M. S., Barch, M. J. & Lawce, H. J.) 213–300 (Wiley, 2017). doi:10.1002/9781119061199.ch6.

