## Supplementary Figures for "An updated reference genome sequence and annotation reveals gene losses and gains underlying naked mole-rat biology"

#### Overview of Supplementary Information

**Supplementary Files:**

**Supplementary File S1**. Tab-separated table proving the Ensembl IDs and gene coordinates for additional candidate pseudogenes.
**Supplementary File S2**. PDF file showing interactive genome viewer screenshots of each tandem duplication identified on mHetGlaV3 (associated with Table 2).
**Supplementary File S3**. Tab-separated file describing breakpoints between mHetGlaV2 and mHetGlaV3 using mHetGlaV3 coordinates.

**Supplementary Tables:**

**Supplementary Table S1.** Description of chromosome number and its relationship to mHetGlaV3, chromosome numbers identified by a karyotype in Romanenko et al., and centricity.
**Supplementary Table S2.** Tab-separated table listing gene symbols annotated to each source on mHetGlaV3.
**Supplementary Table S3.** Tab-separated table listing gene symbols annotated to each source on pHetGlaV3.
**Supplementary Table S4.** Tab-separated table listing aligned reads and peaks for each ChIP-seq and ATAC-seq sample assigned to mHetGlaV3.

**Supplementary Figures:**

***Figure S1.*** *Schematic overview of our TOR-NMR2626 (HetGlaV3) assembly pipeline*
***Figure S2.*** *Pedigree of the samples used in our TOR-NMR2626 assembly.*
***Figure S3.*** *Identification of Y chromosome contigs and SRY gene in the NMR assembly.*
***Figure S4.*** *Example of NMR chromosome 1 scaffolding using the Canadian porcupine (Eredor1) as a reference genome.*
***Figure S5.*** *Evaluation of assembly breakpoints between mHetGlaV2 and mHetGlaV3.*
**Figure S6.** Summary and timeline of current NMR assemblies and technologies. Figure generated with Biorender (https://www.biorender.com/).
***Figure S7.*** *AgNOR sequential staining showed that Chromosome 29 is the only chromosome in the NMR genome with satellites and NOR-positive stalk regions.*
***Figure S8.*** *Interactive Genome Viewer screenshot of regions we identified as mis-annotated in mHetGlaV3.*
***Figure S9.*** *Peak set enrichment of hypothalamus ATAC-seq peaks binned into chromatin states in the naked mole-rat.*
***Figure S10.*** *Improvement of gene-cell and cell-type maker mapping of hypothalamus snRNA-seq data when aligned to HetGla1.2 and mHetGlaV3.*
***Figure S11.*** *Summary of gene inactivation and an example of gene loss in the NMR.*
***Figure S12.*** *Example of the expanded Mhc gene family in Mus musculus (mm10) that is not present in the naked mole-rat (NMR) (mHetGlaV3).*
***Figure S13.*** *Example of a gene loss event of PILRB genes in the naked mole-rat.*
***Figure S14.*** *Four examples of where a naked mole-rat genome assembly using PacBio Hifi and ONT ultra-long reads (HetGlaV4) could resolve regions that were misassembled in mHetGlaV3.*

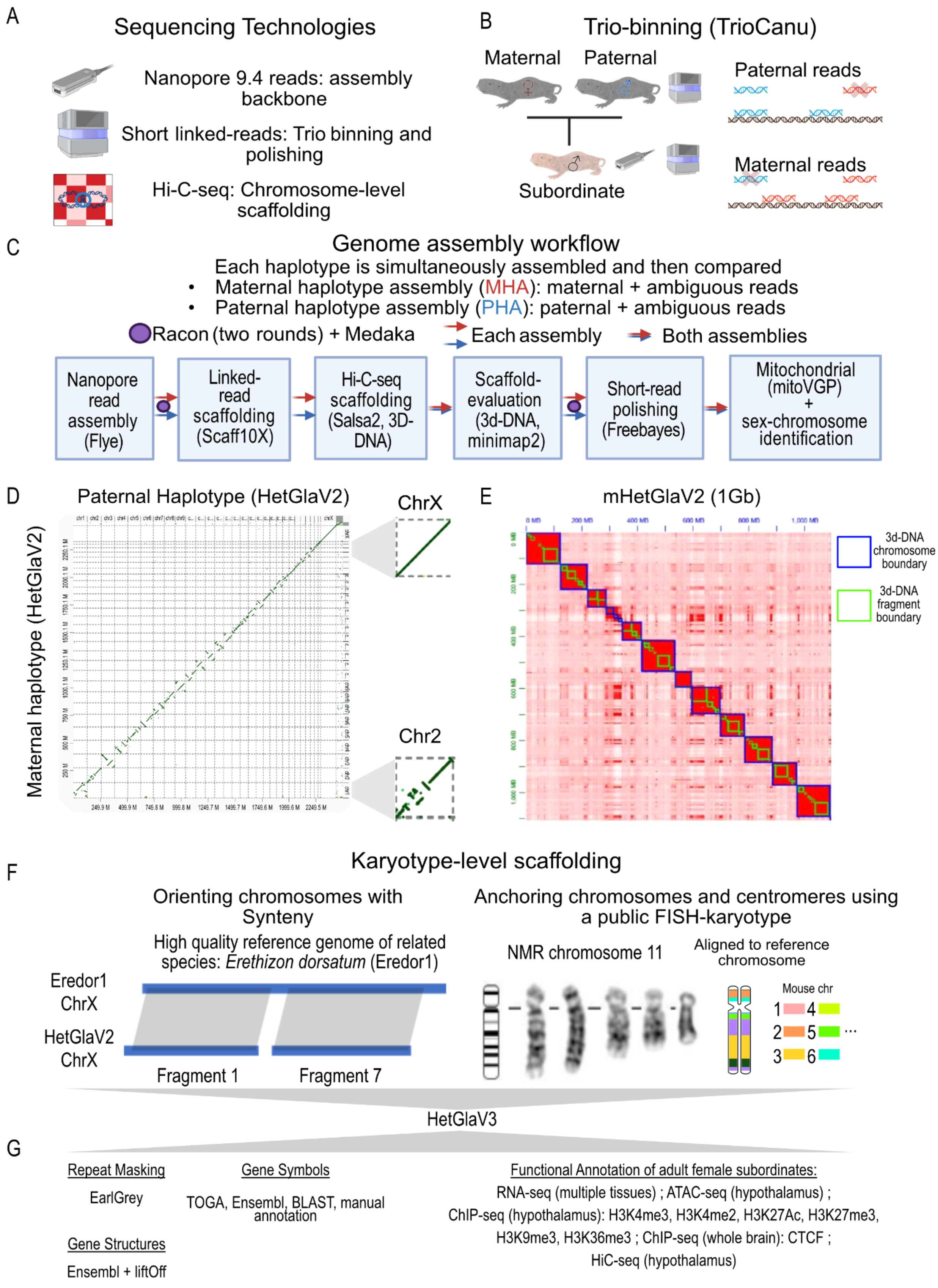
A
Sequencing Technologies
Nanopore 9.4 reads: assembly backbone
Short linked-reads: Trio binning and polishing
Hi-C-seq: Chromosome-level scaffolding
B
Trio-binning (TrioCanu)
Maternal
Paternal
Subordinate
Paternal reads
Maternal reads
C
Genome assembly workflow
Each haplotype is simultaneously assembled and then compared
• Maternal haplotype assembly (MHA): maternal + ambiguous reads
• Paternal haplotype assembly (PHA): paternal + ambiguous reads
Racon (two rounds) + Medaka
Each assembly
Both assemblies
Nanopore read assembly (Flye)
Linked-read scaffolding (Scaff10X)
Hi-C-seq scaffolding (Salsa2, 3D-DNA)
Scaffold-evaluation (3d-DNA, minimap2)
Short-read polishing (Freebayes)
Mitochondrial (mitoVGP) + sex-chromosome identification
D
Paternal Haplotype (HetGlaV2)
Maternal haplotype (HetGlaV2)
ChrX
Chr2
E
mHetGlaV2 (1Gb)
3d-DNA chromosome boundary
3d-DNA fragment boundary
F
Karyotype-level scaffolding
Orienting chromosomes with Synteny
High quality reference genome of related species: Erethizon dorsatum (Eredor1)
Eredor1 ChrX
HetGlaV2 ChrX
Fragment 1
Fragment 7
Anchoring chromosomes and centromeres using a public FISH-karyotype
NMR chromosome 11
Aligned to reference chromosome
Mouse chr
1 4
2 5 ...
3 6
HetGlaV3
G
Repeat Masking
EarlGrey
Gene Symbols
TOGA, Ensembl, BLAST, manual annotation
Gene Structures
Ensembl + liftOff
Functional Annotation of adult female subordinates:
RNA-seq (multiple tissues) ; ATAC-seq (hypothalamus) ;
ChIP-seq (hypothalamus): H3K4me3, H3K4me2, H3K27Ac, H3K27me3, H3K9me3, H3K36me3 ; ChIP-seq (whole brain): CTCF ;
HiC-seq (hypothalamus)

***Supplementary Figure S1.*** *Schematic overview of our TOR-NMR2626 (HetGlaV3) assembly pipeline.* A) Summary of the sequencing technologies used to assemble the naked mole-rat genome, Oxford Nanopore R9.4 reads (40x coverage), 10X linked reads of the TOR-NMR2626 same animal with long reads and their parents (maternal “queen”: TOR-NMR2606, paternal “consort”: TOR-NMR2624), and Hi-C sequencing reads. B) Schematic of trio-binning approach to haplotype individual reads using TrioCanu. Each long read is assigned to a maternal haplotype, paternal haplotype, or unknown haplotype based on k-mer occurrence patterns in the parents. C) Summary of the genome assembly workflow for the maternal haplotype mHetGlaV2 and the paternal haplotype (pHetGlaV2). These are the current Ensembl 112 release assemblies (mHetGlaV2: GCA_944319715, pHetGlaV2; GCA_944319725). Due to the considerable homozygosity in the wild-derived NMRs, each haplotype was assembled using the parental and ambiguous reads. D) Full genome dot plot between mHetGlaV2 and pHetGlaV2 generated with dGenies. The X chromosome is from mHetGlaV2 and provides an example of a self-alignment (i.e., a perfect match). Chromosome 2 provides an example of intrachromosomal misassemblies between mHetGlaV2 and pHetGlaV2. E) Hi-C heatmap of the first gigabase of mHetGlaV2 labelled with chromosome boundaries (blue boxes) and chromosome fragments (green boxes) assigned by 3D-DNA. F) Schematic of how we used the Canadian porcupine genome assembly (EreDor1) to connect the fragments of an assembly. F) After synteny-based chromosome assembly, we assigned structural regions and chromosome identification using our own *de novo* karyotype and data and FISH-karyotype data from Romanenko et al., 2023. G) List of the sequencing technologies and methods used in genome annotation.

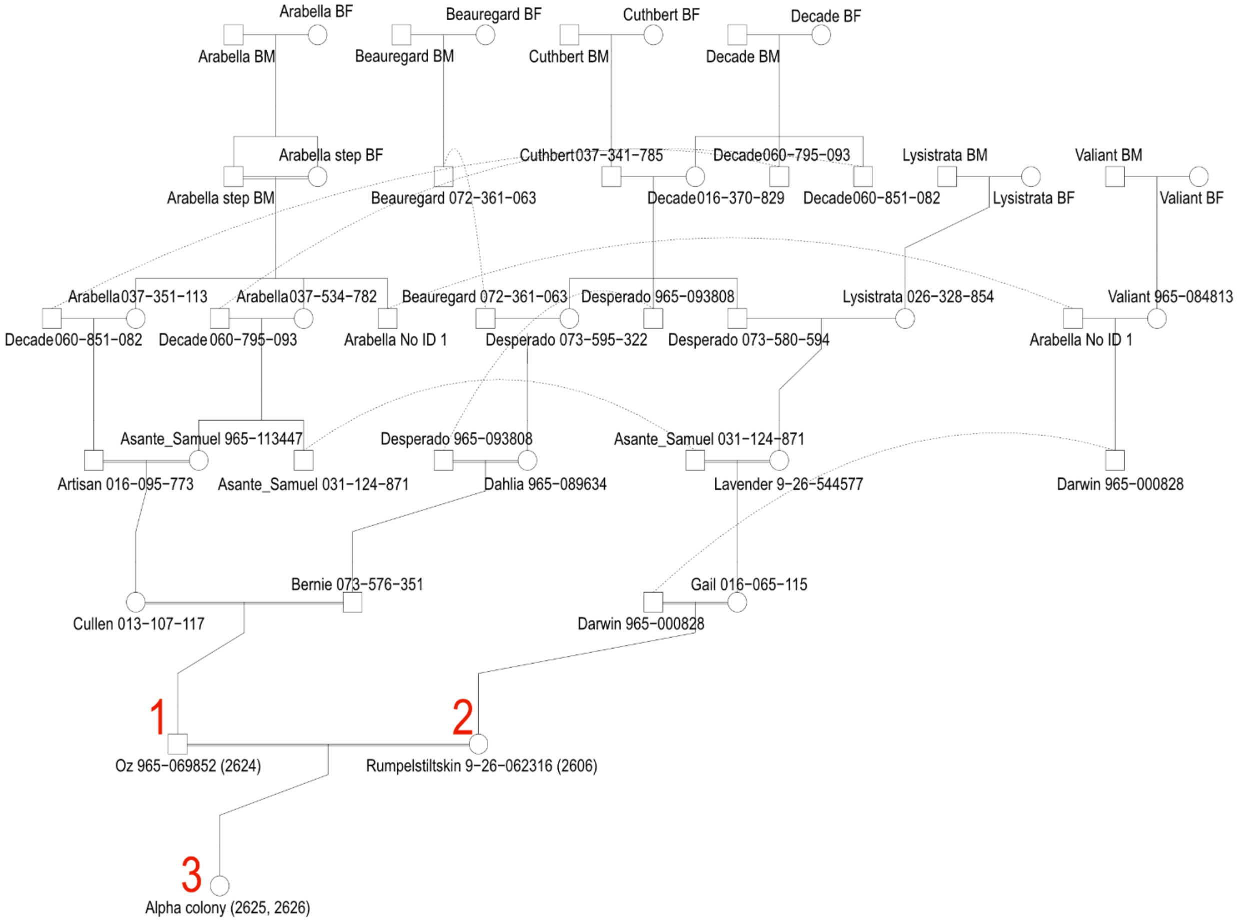


***Figure S2.*** *Pedigree of the samples used in our TOR-NMR2626 assembly.* 1) represents the paternal sample (TOR-NRM2624), 2) represents the maternal sample (TOR-NMR2606), and 3) represents the sample whose assembly we built our reference on (TOR-NMR-2626).

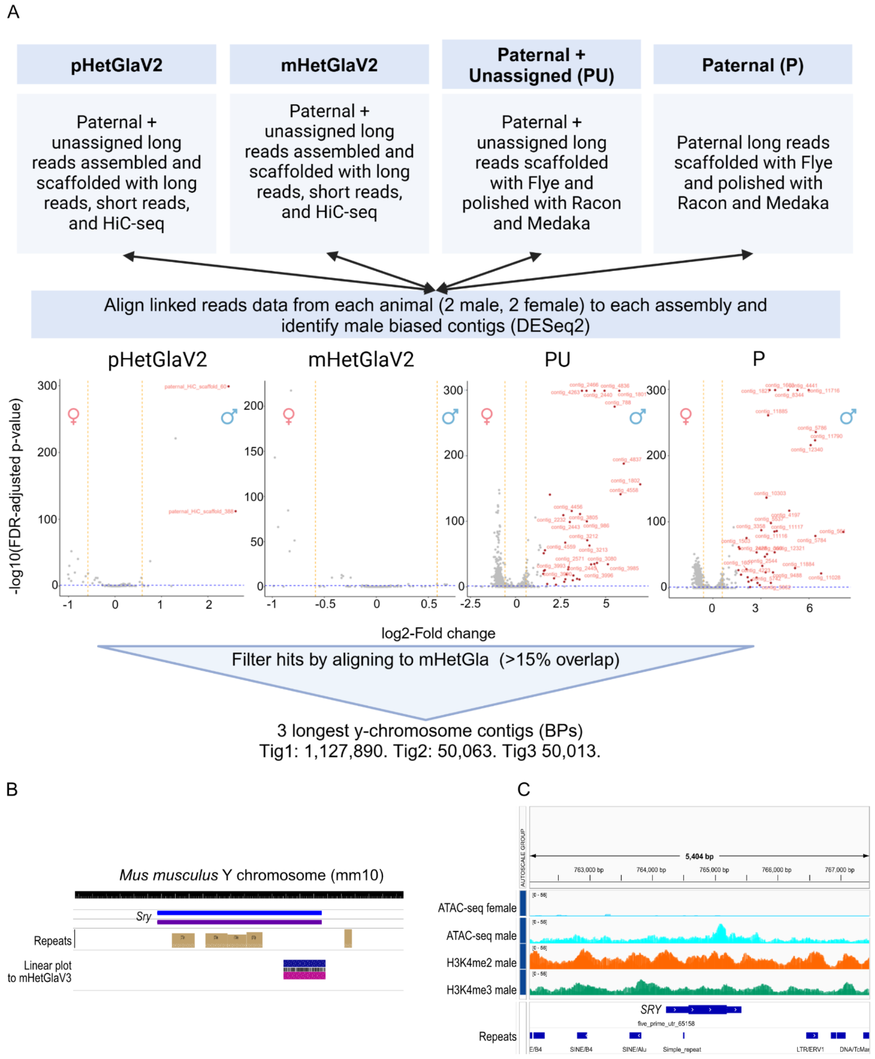


***Figure S3.*** *Identification of Y chromosome contigs and SRY gene in the NMR assembly.* A) Workflow to identify Y chromosome contigs. We first generated four assemblies: mHetGlaV2, pHetGlaV2, and a polished contigged assembly using paternal and unassigned reads, and paternal-only reads. We then aligned our male (N=2) and female (N=2) 10X-linked read data to each of these assemblies, and identified sex-biased contigs (FDR < 0.05 and log2Fold-change > 2). We then filtered contigs with a 15% overlap with mHetGlaV2 to get a candidate set of Y-chromosomal contigs. B) WashU genome browser of the mm10 assembly looking at the *Sry* locus. C) Interactive genome viewer of the aligned region in (B) (ENSHGLG00000045866), which was then identified as *SRY*.

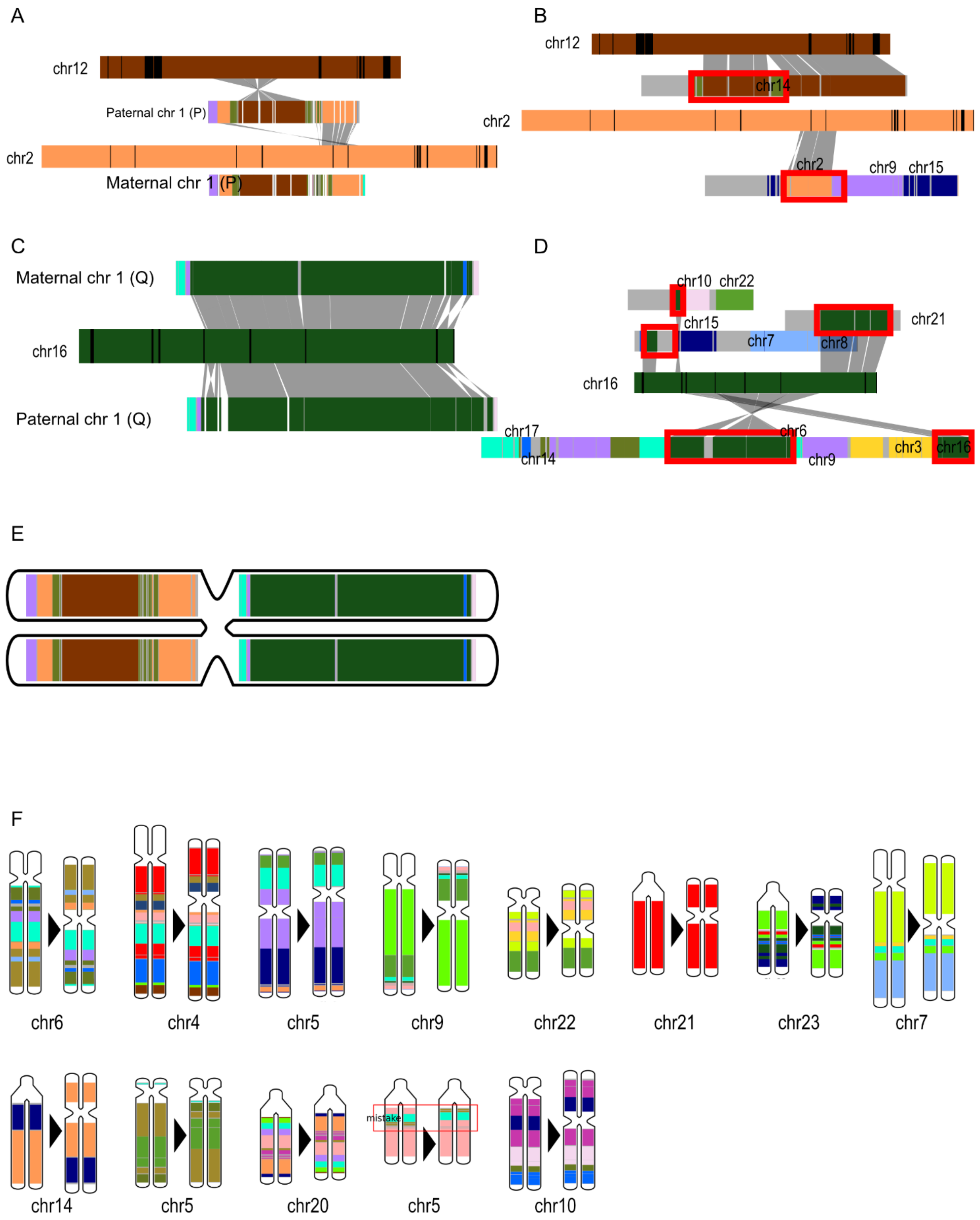
A
chr12
Paternal chr 1 (P)
chr2
Maternal chr 1 (P)
B
chr12
chr14
chr2
chr2
chr9
chr15
C
Maternal chr 1 (Q)
chr16
Paternal chr 1 (Q)
D
chr10
chr22
chr21
chr15
chr7
chr8
chr16
chr17
chr14
chr6
chr9
chr3
chr16
E
F
chr6
chr4
chr5
chr9
chr22
chr21
chr23
chr7
chr14
chr5
chr20
mistake
chr5
chr10

***Figure S4.*** *Example of NMR chromosome 1 scaffolding using the Canadian porcupine (Eredor1) as a reference genome.* All labelled chromosomes on this plot are from EreDor1 other than “Maternal chr1” and “Paternal chr1” A) Example of major alignment blocks between the Canadian porcupine’s chromosome 2 and chromosome 12 and the P arm of HetGlaV3. B) zooming in on (A) to show smaller alignments of other EreDor1 chromosomes onto the P arm of HetGlaV3 chromosome 1. C) Example of major alignment blocks chromosome 16 of *EreDor1* and the Q arm. D) Zooming in on (C) to show smaller alignment blocks between other *EreDor1* chromosomes and the Q arm of HetGlaV3 chromosome 1. E) Full chromosome painting of NMR chromosome 1 with centromere placed. F) NMR chromosomes whose centromeres were re-arranged after integrating FISH karyotype data from Romanenko et al., 2023. We also identified a misfolded inversion in chromosome 5, which we corrected.

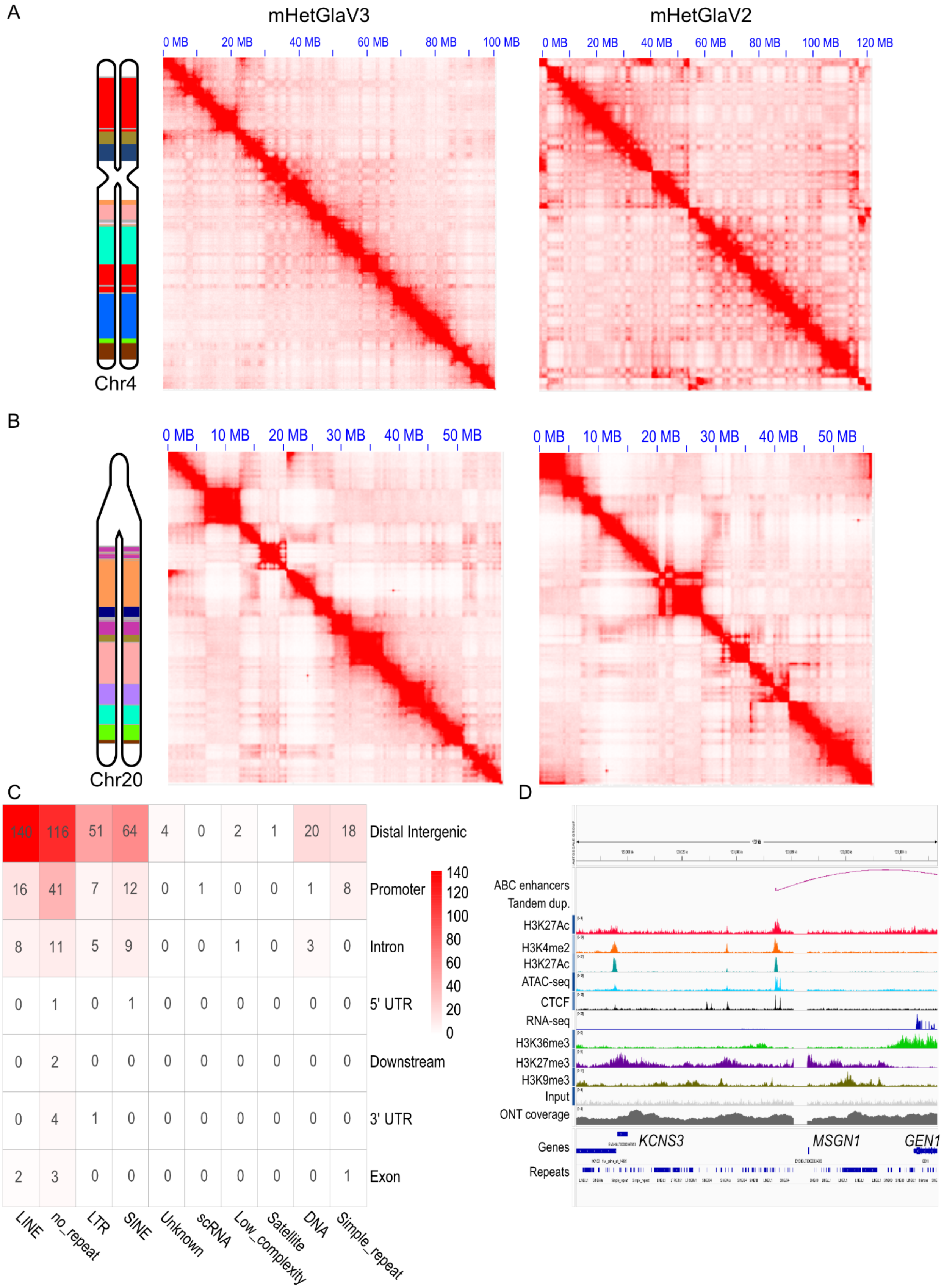
A
mHetGlaV3
mHetGlaV2
0 MB
20 MB
40 MB
60 MB
80 MB
100 MB
120 MB
Chr4
B
0 MB
10 MB
20 MB
30 MB
40 MB
50 MB
Chr20
C
140 116 51 64 4 0 2 1 20 18 Distal Intergenic
16 41 7 12 0 1 0 0 1 8 Promoter
8 11 5 9 0 0 1 0 3 0 Intron
0 1 0 1 0 0 0 0 0 0 5' UTR
0 2 0 0 0 0 0 0 0 0 Downstream
0 4 1 0 0 0 0 0 0 0 3' UTR
2 3 0 0 0 0 0 0 0 1 Exon
140
120
100
80
60
40
20
0
LINE
no_repeat
LTR
SINE
Unknown
scRNA
Low_complexity
Satellite
DNA
Simple_repeat
D
ABC enhancers
Tandem dup.
H3K27Ac
H3K4me2
H3K27Ac
ATAC-seq
CTCF
RNA-seq
H3K36me3
H3K27me3
H3K9me3
Input
ONT coverage
Genes
Repeats
KCNS3
MSGN1
GEN1

***Figure S5.*** *Evaluation of assembly breakpoints between mHetGlaV2 and mHetGlaV3.* A) Hi-C contact maps for a telocentric chromosome (chromosome 4) (A) and an acrocentric chromosome (chromosome 20) (B) before and after additional scaffolding from evolutionary and FISH karyotype data. In A-B, the left is the final painted chromosome, the middle is a Hi-C contact map for that chromosome in mHetGlaV3, and the right is a Hi-C contact map for that chromosome in mHetGlaV2. C) Heatmap identifying the gene and TE annotation of assembly breakpoints between mHetGlaV2 and mHetGlaV3. Heat and numbers both represent the number of regions annotating to that breakpoint. D) Example of an assembly breakpoint that lies outside of a TE and within the *MSGN1* gene promoter on the interactive genome viewer of the mHetGlaV3 genome assembly.

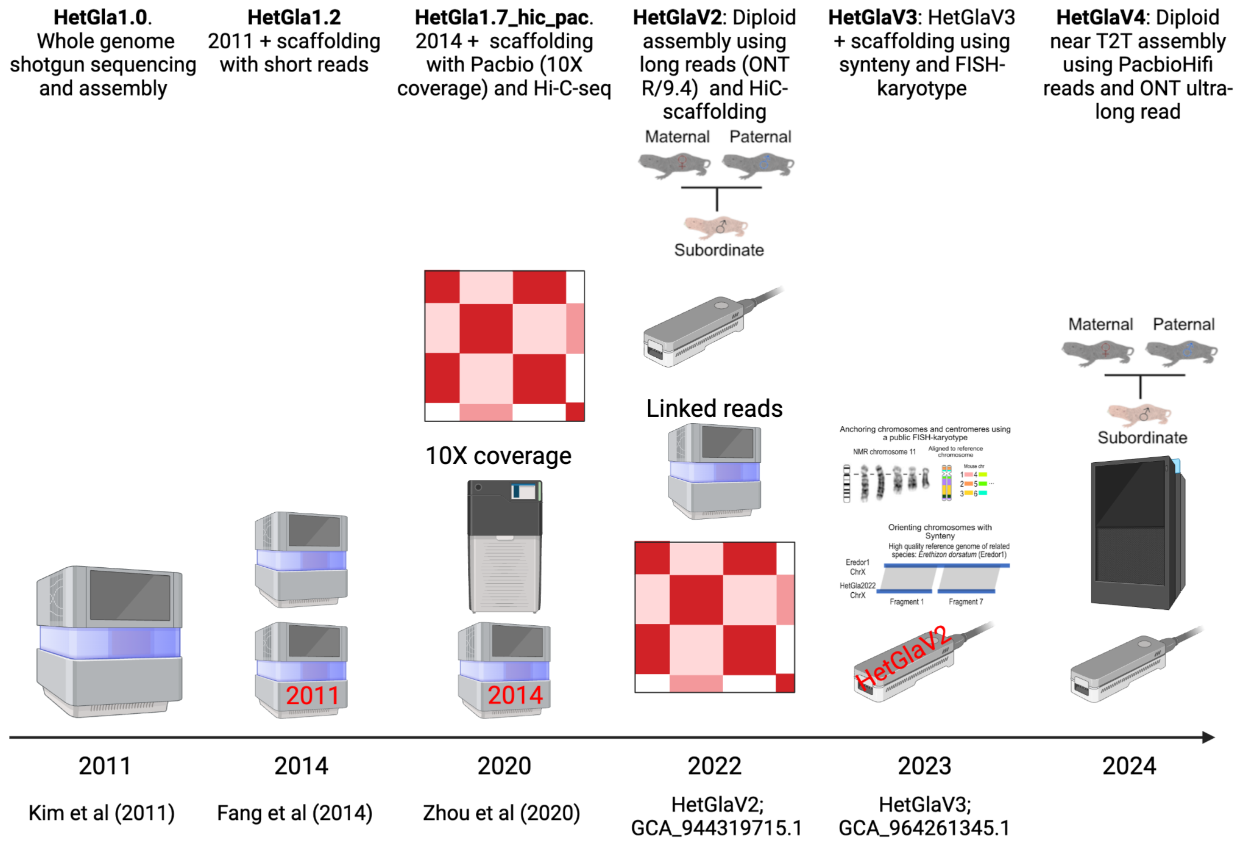


***Figure S6.*** *Summary and timeline of current NMR assemblies and technologies.* Figure generated with Biorender (https://www.biorender.com/).

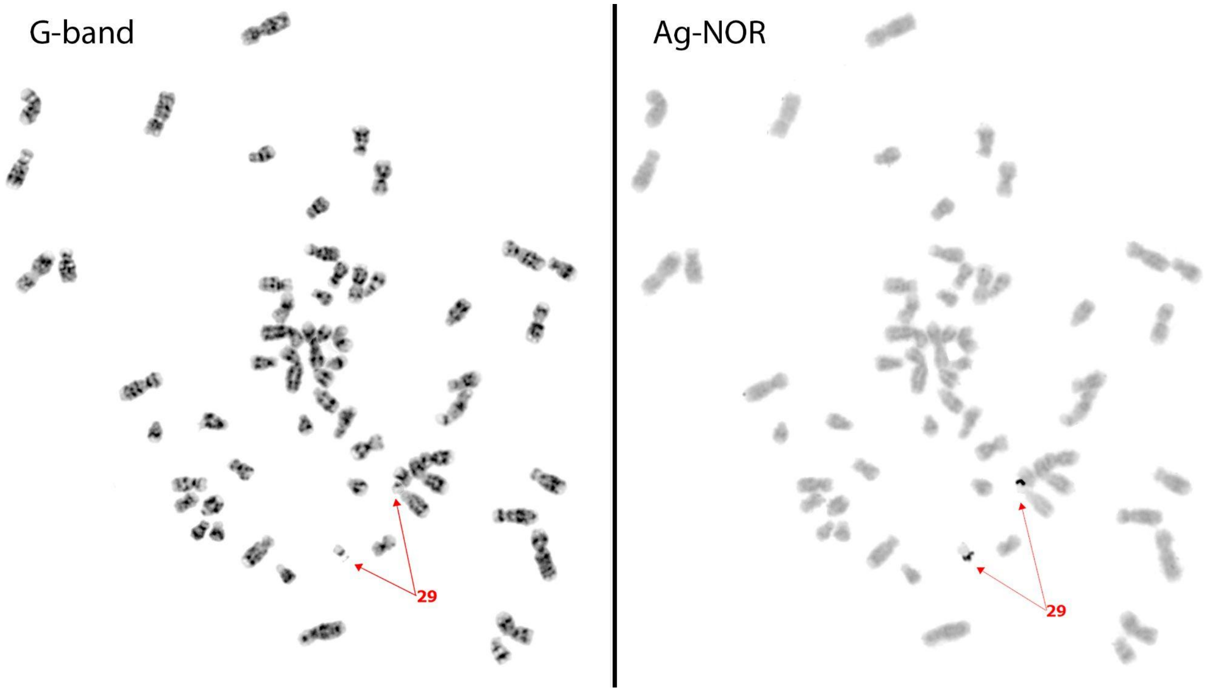


***Figure S7.*** *AgNOR sequential staining showed that Chromosome 29 is the only chromosome in the NMR genome with satellites and NOR-positive stalk regions.*

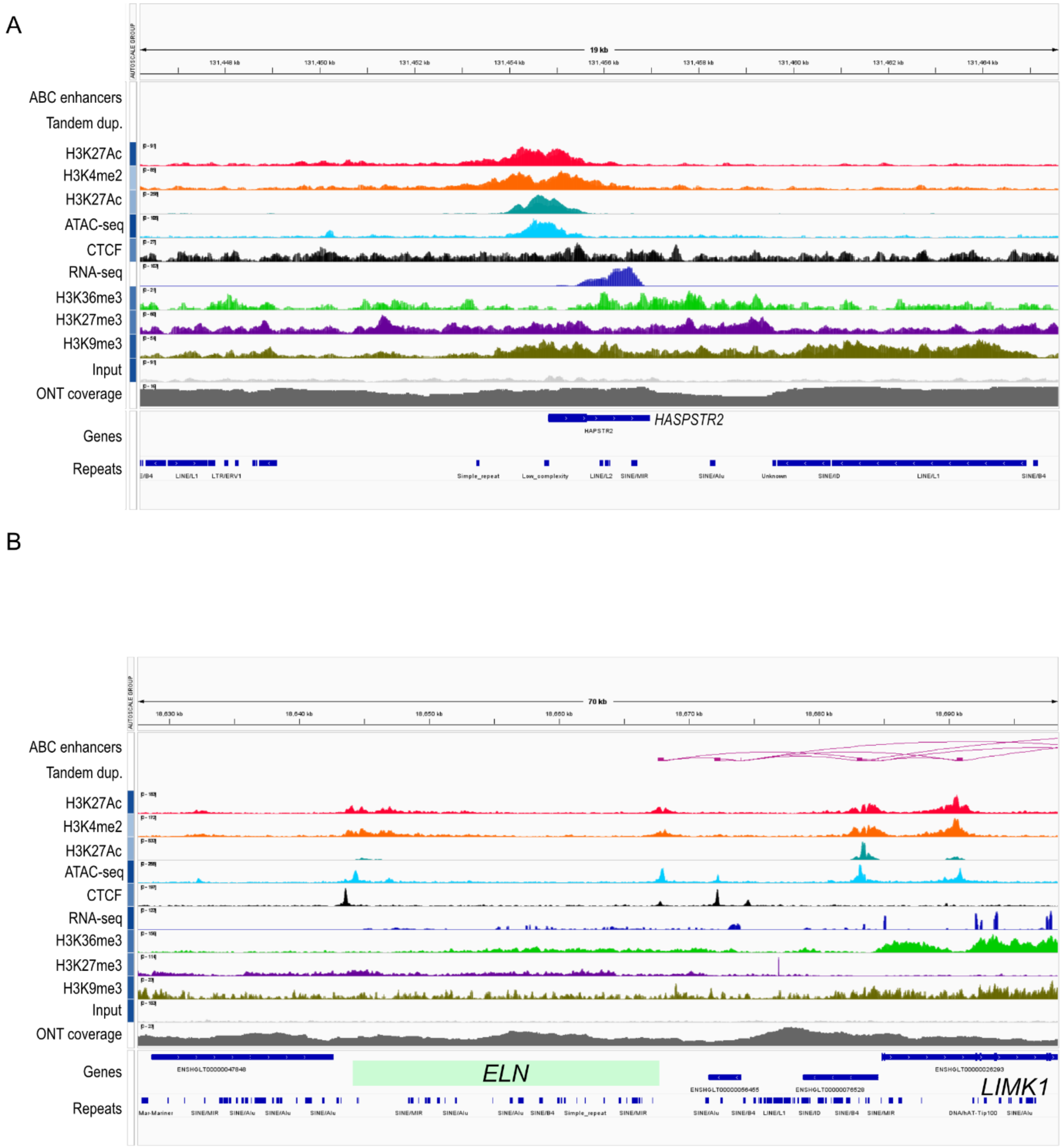


***Figure S8.*** *Interactive Genome Viewer screenshot of regions we identified as mis-annotated in mHetGlaV3.* A) Screenshot of the *HAPSTR2* gene, which was not identified with a gene symbol before the incorporation of the gene annotations in Ensembl 111. B) Screenshot of the locus containing the elastin (*ELN*) gene. The *ELN* gene was not annotated with a gene structure despite the presence of alignments in our RNA-seq and epigenetic data.

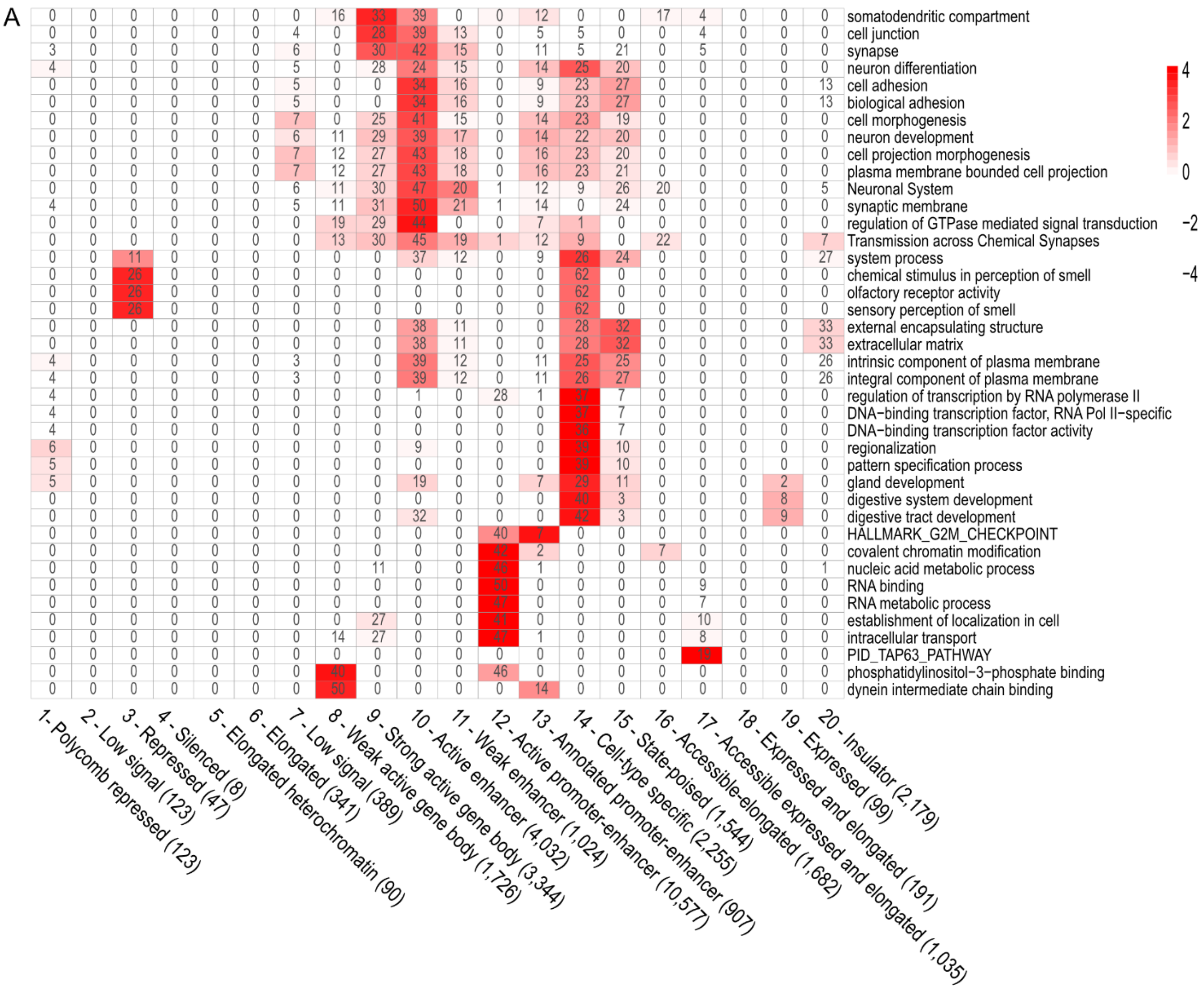
A
somatodendritic compartment
cell junction
synapse
neuron differentiation
cell adhesion
biological adhesion
cell morphogenesis
neuron development
cell projection morphogenesis
plasma membrane bounded cell projection
Neuronal System
synaptic membrane
regulation of GTPase mediated signal transduction
Transmission across Chemical Synapses
system process
chemical stimulus in perception of smell
olfactory receptor activity
sensory perception of smell
external encapsulating structure
extracellular matrix
intrinsic component of plasma membrane
integral component of plasma membrane
regulation of transcription by RNA polymerase II
DNA−binding transcription factor, RNA Pol II−specific
DNA−binding transcription factor activity
regionalization
pattern specification process
gland development
digestive system development
digestive tract development
HALLMARK_G2M_CHECKPOINT
covalent chromatin modification
nucleic acid metabolic process
RNA binding
RNA metabolic process
establishment of localization in cell
intracellular transport
PID_TAP63_PATHWAY
phosphatidylinositol−3−phosphate binding
dynein intermediate chain binding
4
2
0
−2
−4
1- Polycomb repressed (123)
2 - Low signal (123)
3 - Repressed (47)
4 - Silenced (8)
5 - Elongated heterochromatin (90)
6 - Elongated (341)
7 - Low signal (389)
8 - Weak active gene body (1,726)
9 - Strong active gene body (3,344)
10 - Active enhancer (4,032)
11 - Weak enhancer (1,024)
12 - Active promoter-enhancer (10,577)
13 - Annotated promoter-enhancer (907)
14 - Cell-type specific (2,255)
15 - State-poised (1,544)
16 - Accessible-elongated (1,682)
17 - Accessible expressed and elongated (1,035)
18 - Expressed and elongated (191)
19 - Expressed (99)
20 - Insulator (2,179)

B

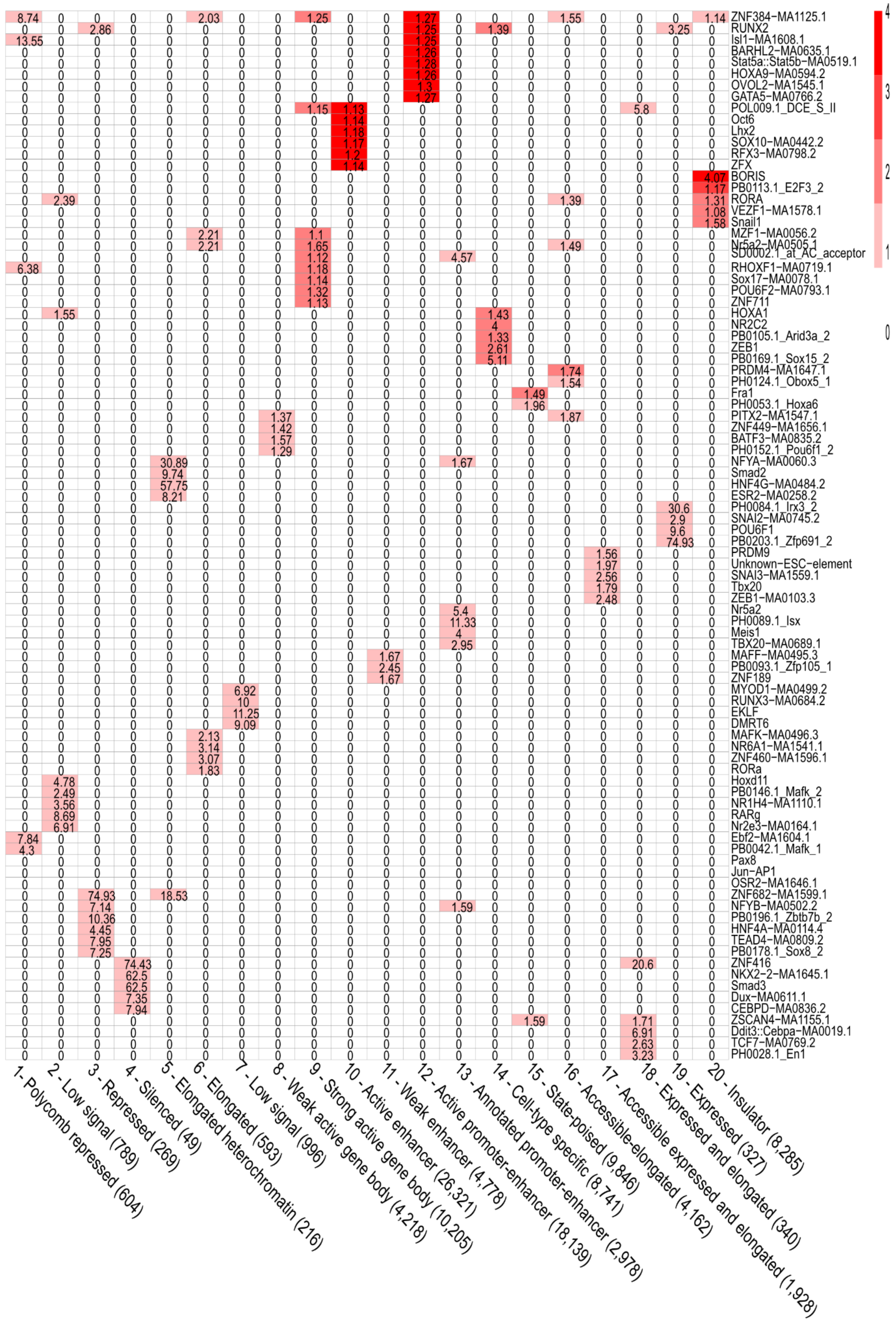
ZNF384-MA1125.1
RUNX2
Isl1-MA1608.1
BARHL2-MA0635.1
Stat5a::Stat5b-MA0519.1
HOXA9-MA0594.2
OVOL2-MA1545.1
GATA5-MA0766.2
POL009.1_DCE_S_II
Oct6
Lhx2
SOX10-MA0442.2
RFX3-MA0798.2
ZFX
BORIS
PB0113.1_E2F3_2
RORA
VEZF1-MA1578.1
Snail1
MZF1-MA0056.2
Nr5a2-MA0505.1
SD0002.1_at_AC_acceptor
RHOXF1-MA0719.1
Sox17-MA0078.1
POU6F2-MA0793.1
ZNF711
HOXA1
NR2C2
PB0105.1_Arid3a_2
ZEB1
PB0169.1_Sox15_2
PRDM4-MA1647.1
PH0124.1_Obox5_1
Fra1
PH0053.1_Hoxa6
PITX2-MA1547.1
ZNF449-MA1656.1
BATF3-MA0835.2
PH0152.1_Pou6f1_2
NFYA-MA0060.3
Smad2
HNF4G-MA0484.2
ESR2-MA0258.2
PH0084.1_Irx3_2
SNAI2-MA0745.2
POU6F1
PB0203.1_Zfp691_2
PRDM9
Unknown-ESC-element
SNAI3-MA1559.1
Tbx20
ZEB1-MA0103.3
Nr5a2
PH0089.1_Isx
Meis1
TBX20-MA0689.1
MAFF-MA0495.3
PB0093.1_Zfp105_1
ZNF189
MYOD1-MA0499.2
RUNX3-MA0684.2
EKLF
DMRT6
MAFK-MA0496.3
NR6A1-MA1541.1
ZNF460-MA1596.1
RORa
Hoxd11
PB0146.1_Mafk_2
NR1H4-MA1110.1
RARg
Nr2e3-MA0164.1
Ebf2-MA1604.1
PB0042.1_Mafk_1
Pax8
Jun-AP1
OSR2-MA1646.1
ZNF682-MA1599.1
NFYB-MA0502.2
PB0196.1_Zbtb7b_2
HNF4A-MA0114.4
TEAD4-MA0809.2
PB0178.1_Sox8_2
ZNF416
NKX2-2-MA1645.1
Smad3
Dux-MA0611.1
CEBPD-MA0836.2
ZSCAN4-MA1155.1
Ddit3::Cebpa-MA0019.1
TCF7-MA0769.2
PH0028.1_En1
1- Polycomb repressed (604)
2 - Low signal (789)
3 - Repressed (269)
4 - Silenced (49)
5 - Elongated heterochromatin (216)
6 - Elongated (593)
7 - Low signal (996)
8 - Weak active gene body (4,218)
9 - Strong active gene body (10,205)
10 - Active enhancer (26,321)
11 - Weak enhancer (4,778)
12 - Active promoter-enhancer (18,139)
13 - Annotated promoter-enhancer (2,978)
14 - Cell-type specific (8,741)
15 - State-poised (9,846)
16 - Accessible-elongated (4,162)
17 - Accessible expressed and elongated (1,928)
18 - Expressed and elongated (340)
19 - Expressed (327)
20 - Insulator (8,285)
4
3
2
1
0

C

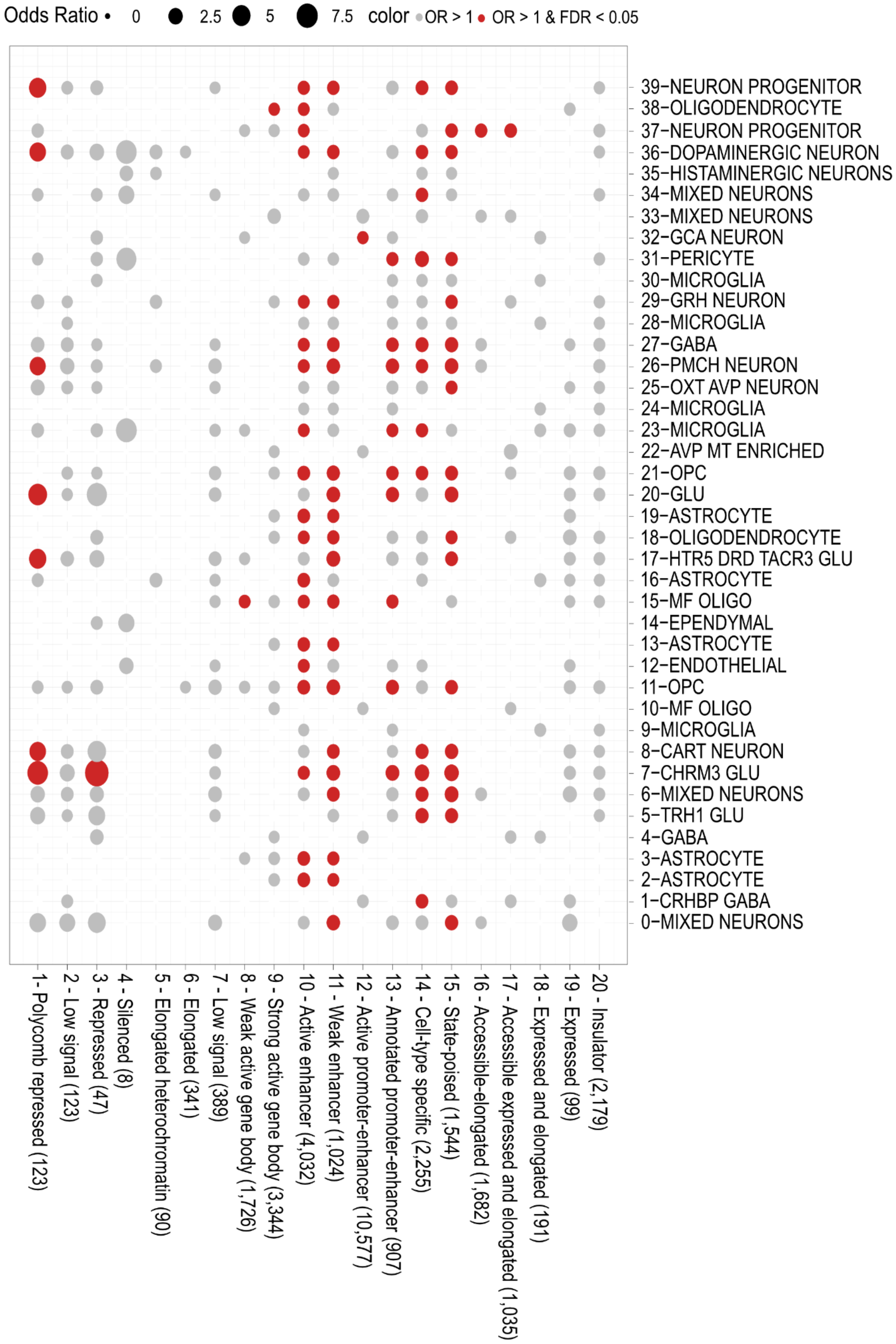

Odds Ratio • 0 2.5 5 7.5 color • OR > 1 • OR > 1 & FDR < 0.05
39-NEURON PROGENITOR
38-OLIGODENDROCYTE
37-NEURON PROGENITOR
36-DOPAMINERGIC NEURON
35-HISTAMINERGIC NEURONS
34-MIXED NEURONS
33-MIXED NEURONS
32-GCA NEURON
31-PERICYTE
30-MICROGLIA
29-GRH NEURON
28-MICROGLIA
27-GABA
26-PMCH NEURON
25-OXT AVP NEURON
24-MICROGLIA
23-MICROGLIA
22-AVP MT ENRICHED
21-OPC
20-GLU
19-ASTROCYTE
18-OLIGODENDROCYTE
17-HTR5 DRD TACR3 GLU
16-ASTROCYTE
15-MF OLIGO
14-EPENDYMAL
13-ASTROCYTE
12-ENDOTHELIAL
11-OPC
10-MF OLIGO
9-MICROGLIA
8-CART NEURON
7-CHRM3 GLU
6-MIXED NEURONS
5-TRH1 GLU
4-GABA
3-ASTROCYTE
2-ASTROCYTE
1-CRHBP GABA
0-MIXED NEURONS
1 - Polycomb repressed (123)
2 - Low signal (123)
3 - Repressed (47)
4 - Silenced (8)
5 - Elongated heterochromatin (90)
6 - Elongated (341)
7 - Low signal (389)
8 - Weak active gene body (1,726)
9 - Strong active gene body (3,344)
10 - Active enhancer (4,032)
11 - Weak enhancer (1,024)
12 - Active promoter-enhancer (10,577)
13 - Annotated promoter-enhancer (907)
14 - Cell-type specific (2,255)
15 - State-poised (1,544)
16 - Accessible-elongated (1,682)
17 - Accessible expressed and elongated (1,035)
18 - Expressed and elongated (191)
19 - Expressed (99)
20 - Insulator (2,179)

***Figure S9.*** *Peak set enrichment of hypothalamus ATAC-seq peaks binned into chromatin states in the naked mole-rat.* A) Heatmap of gene-set enrichment of ATAC-seq peaks annotated to non-intergenic (i.e., promoter, UTR, intron, or exonic) regions. Rows are gene sets that were in the top five most significant FDR-adjusted p-value of at least one chromatin state, and columns are chromatin states (# annotated genes per state). Heat represents the -log10(FDR-adjusted p-value of enrichment) normalized by each chromatin state (column) and the number inside the heatmap represents the number of genes that overlap with the chromatin state and gene set. B) Transcription factor enrichment and motif discovery using HOMER of ATAC-seq peaks mapped to each chromatin state. Rows are motifs that are in the top five most significant p-value of at least one chromatin state and columns are chromatin states. Heat represents the -log10(p-value) and the number within each heatmap represents the ratio of % motif in state-specific peak to % motif in all ATAC-seq peaks. C) Bubblegum plot of cell-type specific enrichment of peaks annotated to non-intergenic regions. Rows were cell-types in the hypothalamus identified using snRNA-seq in the female mouse hypothalamus and columns are chromatin states. Circle size represents the over-representation of cell-type markers in that cell-type compared to all other cell-types. Grey circles are not significantly over-represented (odds ratio > 1) and red circles are significantly over-represented (FDR < 0.05, odds ratio > 1).

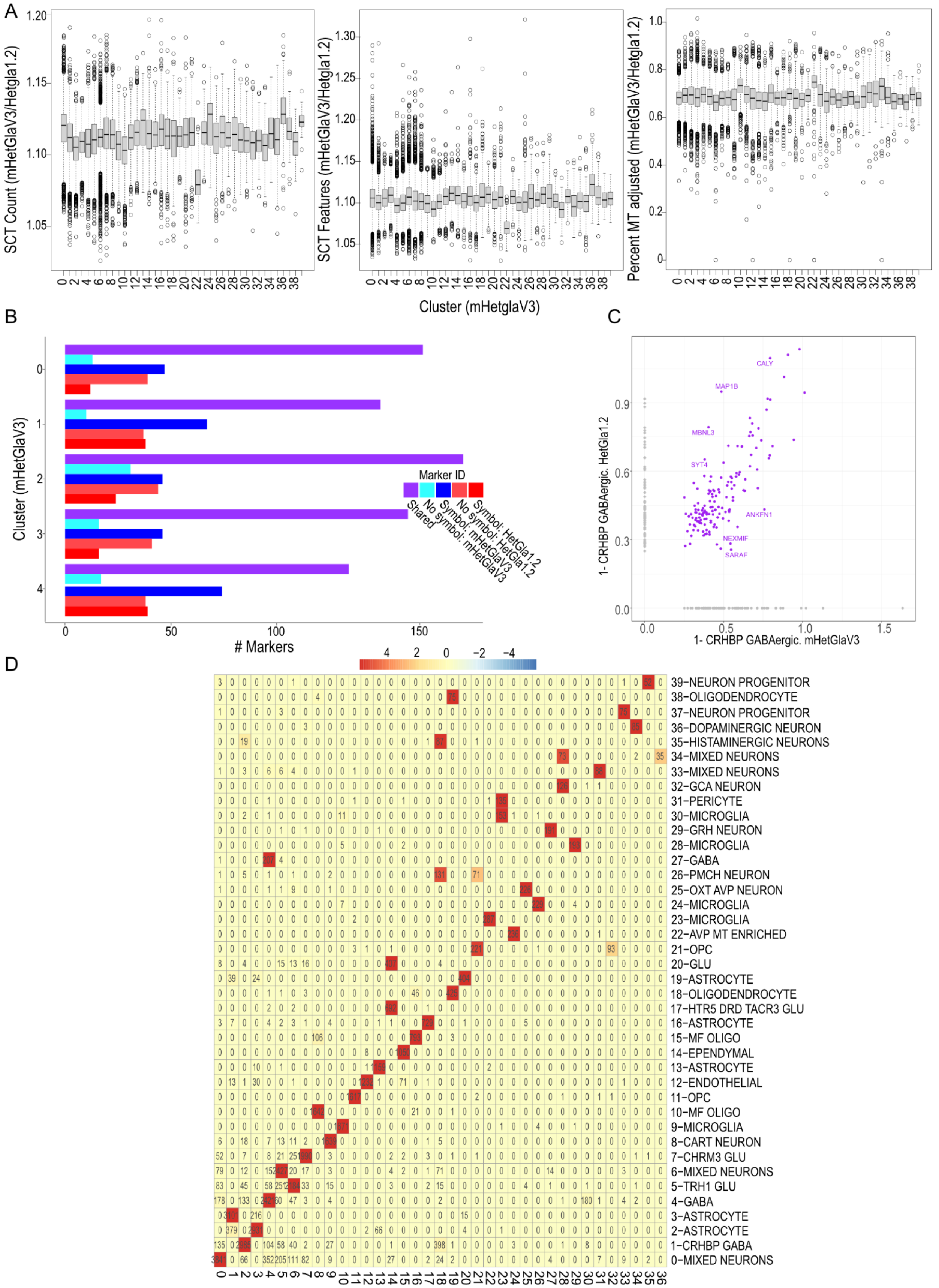
A
SCT Count (mHetGlaV3/Hetgla1.2)
SCT Features (mHetGlaV3/Hetgla1.2)
Percent MT adjusted (mHetGlaV3/Hetgla1.2)
Cluster (mHetglaV3)
B
Cluster (mHetGlaV3)
### Markers
Marker ID
Shared
No Symbol: mHetGlaV3
No symbol: HetGla1.2
No Symbol: mHetGlaV3
Symbol: HetGla1.2
C
1- CRHBP GABAergic. HetGla1.2
1- CRHBP GABAergic. mHetGlaV3
CALY
MAP1B
MBNL3
SYT4
ANKFN1
NEXMIF
SARAF
D
39−NEURON PROGENITOR
38−OLIGODENDROCYTE
37−NEURON PROGENITOR
36−DOPAMINERGIC NEURON
35−HISTAMINERGIC NEURONS
34−MIXED NEURONS
33−MIXED NEURONS
32−GCA NEURON
31−PERICYTE
30−MICROGLIA
29−GRH NEURON
28−MICROGLIA
27−GABA
26−PMCH NEURON
25−OXT AVP NEURON
24−MICROGLIA
23−MICROGLIA
22−AVP MT ENRICHED
21−OPC
20−GLU
19−ASTROCYTE
18−OLIGODENDROCYTE
17−HTR5 DRD TACR3 GLU
16−ASTROCYTE
15−MF OLIGO
14−EPENDYMAL
13−ASTROCYTE
12−ENDOTHELIAL
11−OPC
10−MF OLIGO
9−MICROGLIA
8−CART NEURON
7−CHRM3 GLU
6−MIXED NEURONS
5−TRH1 GLU
4−GABA
3−ASTROCYTE
2−ASTROCYTE
1−CRHBP GABA
0−MIXED NEURONS

***Figure S10.*** *Improvement of gene-cell and cell-type maker mapping of hypothalamus snRNA-seq data when aligned to HetGla1.2 and mHetGlaV3.* A) Boxplots of the ratio of single-cell transformed (SCT) counts (left) SCT features (middle) and % mitochondria (right) between mHetGlaV3 and HetGla1.2, separated by clusters found in mHetGlaV3. Each point is a cell. Values greater than 1 are greater in mHetGlaV3, and values less than 1 are greater in HetGla1.2. B) Barplot of the number of marker genes assigned to the four largest clusters where bar height represents the number of genes. Purple bars represent genes shared by both clusters, light blue represents mHetGlaV3 specific cluster with no gene symbol assigned, blue represents mHetGlaV3 specific cluster with a gene symbol assigned, light red represents HetGla1.2 specific cluster with no gene symbol assigned, and red represents HetGla1.2 specific cluster with a gene symbol assigned. C) Scatterplot of the fold-changes of cell-type markers for CRHBP+ GABAergic neurons when aligning data to mHetGlaV3 (x-axis) and HetGla1.2 (Y-axis). Points are genes. Points in purple represent shared markers (FDR < 1e-10, fold-change > 1.2), where labelled genes have a difference of more than three standard deviations of fold-change between shared markers. Grey points are markers specific to each genome assembly. D) Heatmap of cell-cluster assignment between mHetGlaV3 and HetGla1.2. Rows are clusters identified when analyzing snRNA-seq data aligned to mHetGlaV3 and columns are clusters identified when analyzing snRNA-seq data aligned to HetGla1.2. Heat represents the number of cells in each cluster pair, and the number of cells in each cluster pair is found in the heatmap.

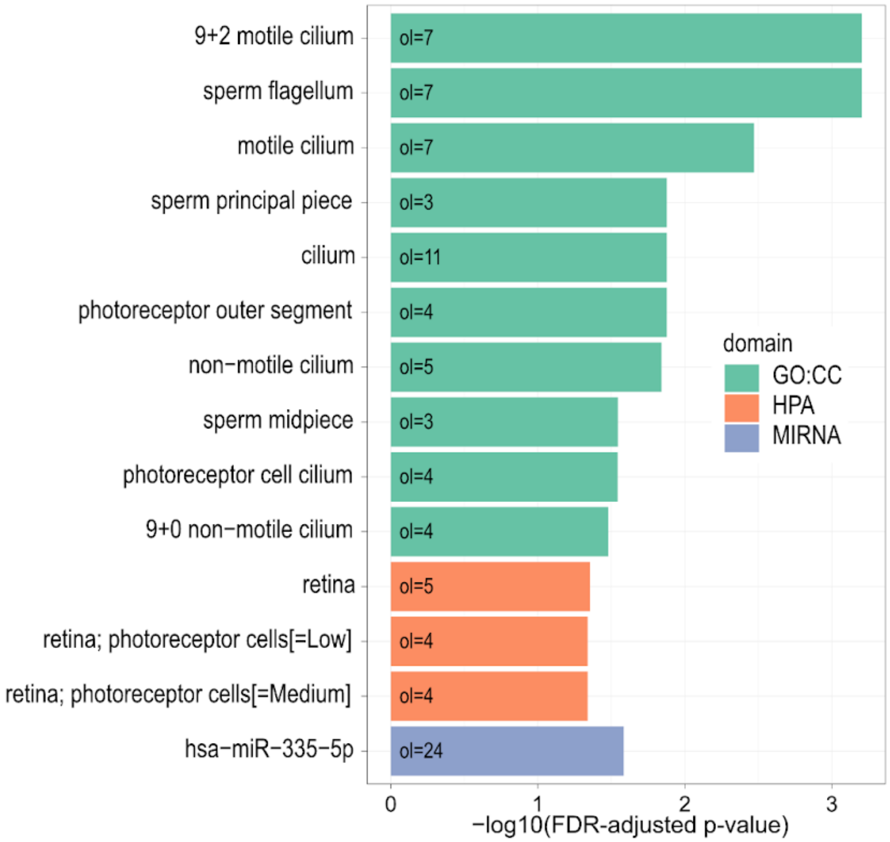


***Figure S11.*** *Summary of gene inactivation and an example of gene loss in the NMR.* A) Barplot of enriched gene sets of genes that were identified as fully inactivated in both mHetGlaV3 and pHetGlaV3 using the tool to infer orthologs from genome alignments (TOGA). We applied an additional gene expression filter to these genes, such that these genes needed less than 0.5 reads per kilobase per million gene expression in all tissues stored in Bens et al., 2018. Rows are gene sets, bar height is the -log10(FDR-adjusted p-value), and “ol” represents the number of genes in that gene set.

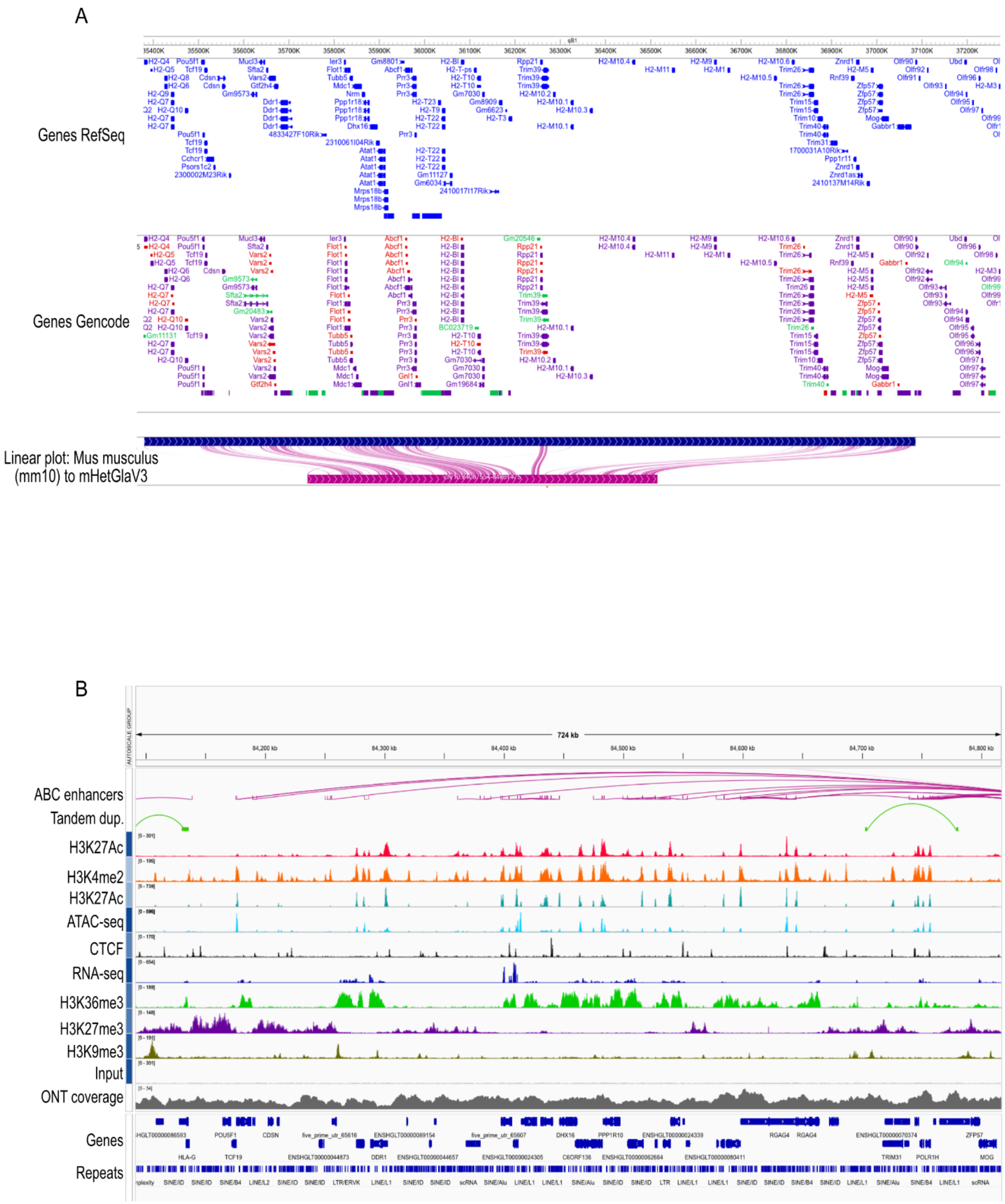


***Figure S12.*** *Example of the expanded Mhc gene family in Mus musculus (mm10) that is not present in the naked mole-rat (NMR) (mHetGlaV3).* A) Washu epigenome browser of the mm10 assembly at the *Mhc* gene family region, showing a linear plot between the mm10 and mHetGlaV3 assemblies. B) IGV plot of the surrounding genomic regions of what would be the *Mhc* gene family in *Mus musculus*, in the NMR (mHetGlaV3).

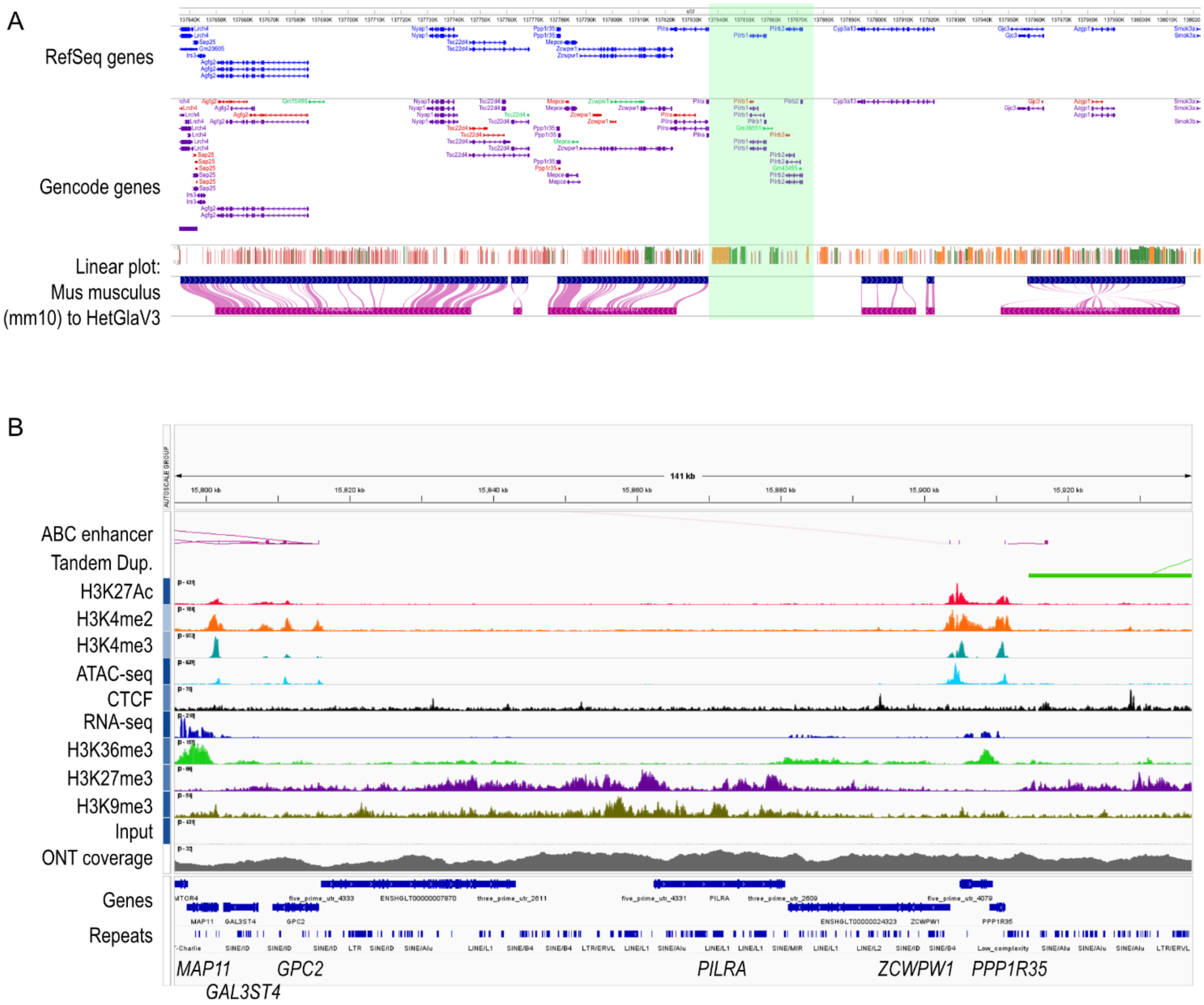


***Figure S13.*** *Example of a gene loss event of PILRB genes in the naked mole-rat.* A) WashU epigenome browser of the paired immunoglobulin-like receptor (*PILR*) locus in *Mus musculus* (mm10), with a linear plot showing alignments to the NMR (mHetGlaV3). B) IGV screenshot of the *PILR* locus in the NMR.

HetGlaV4. IGKV: Chr6: 67,936,558-68,906,787

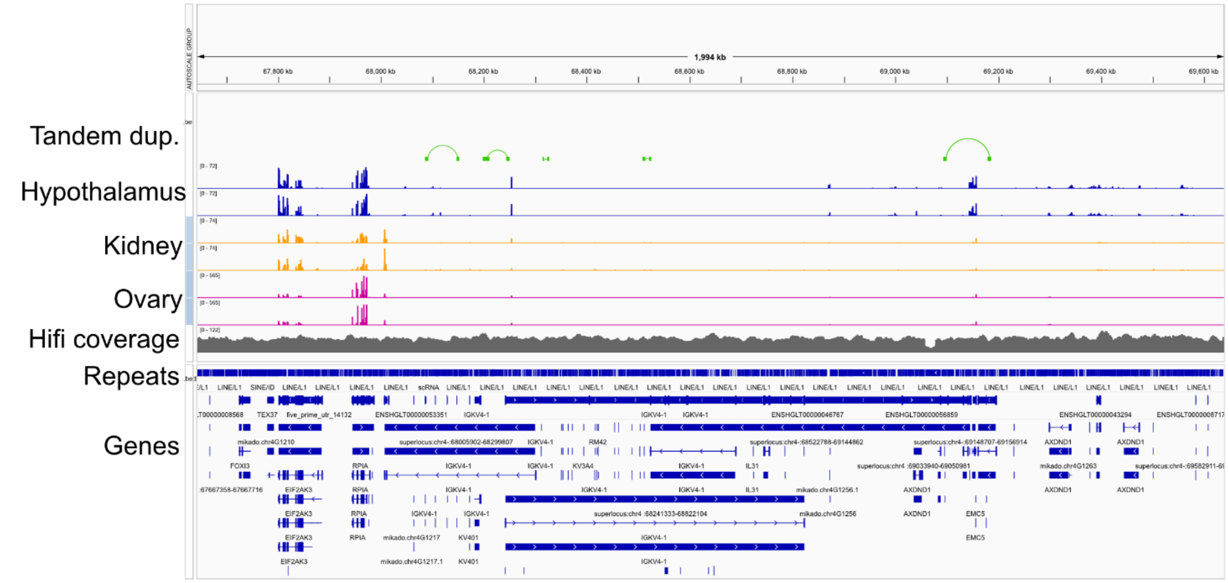


mHetGlaV3. IGKV: Chr4: 55,877,651-57,249,372

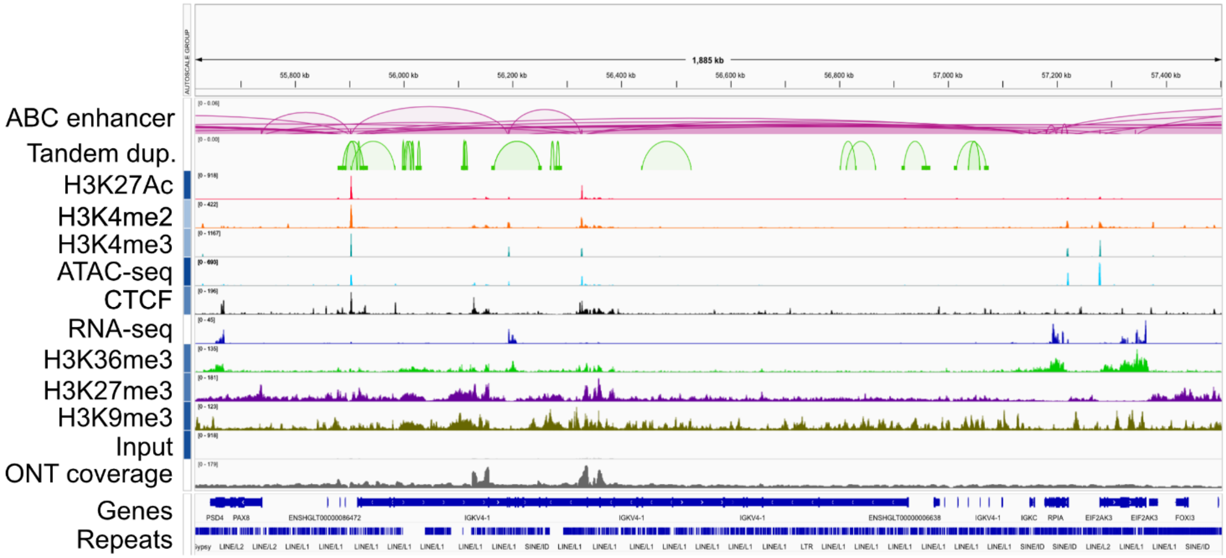

HetGlaV4. OLFR: Chr12: 19,051,455-37,537,326

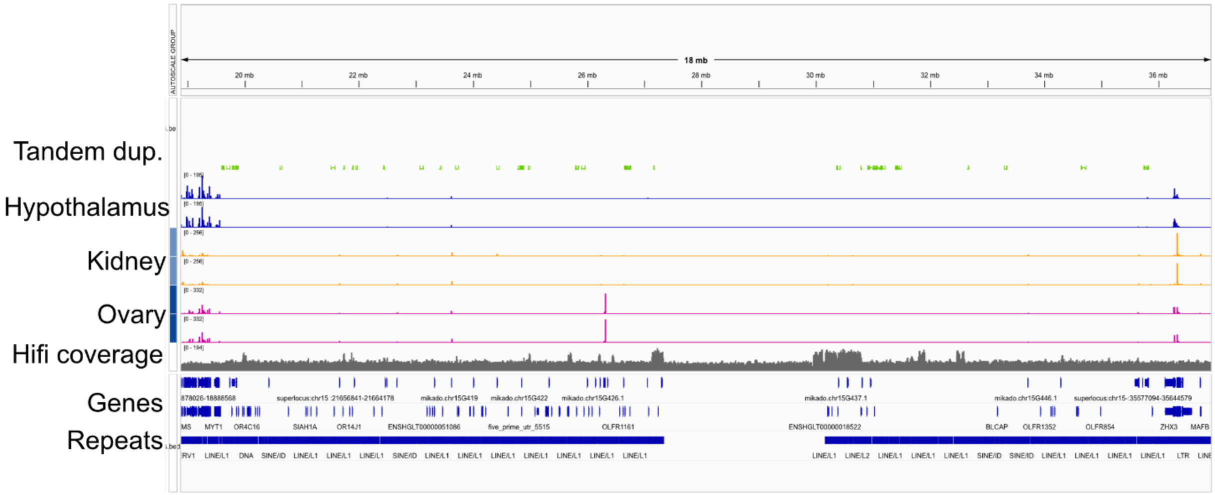


mHetGlaV3 OLFR: Chr2: 37,998,924-50,068,501

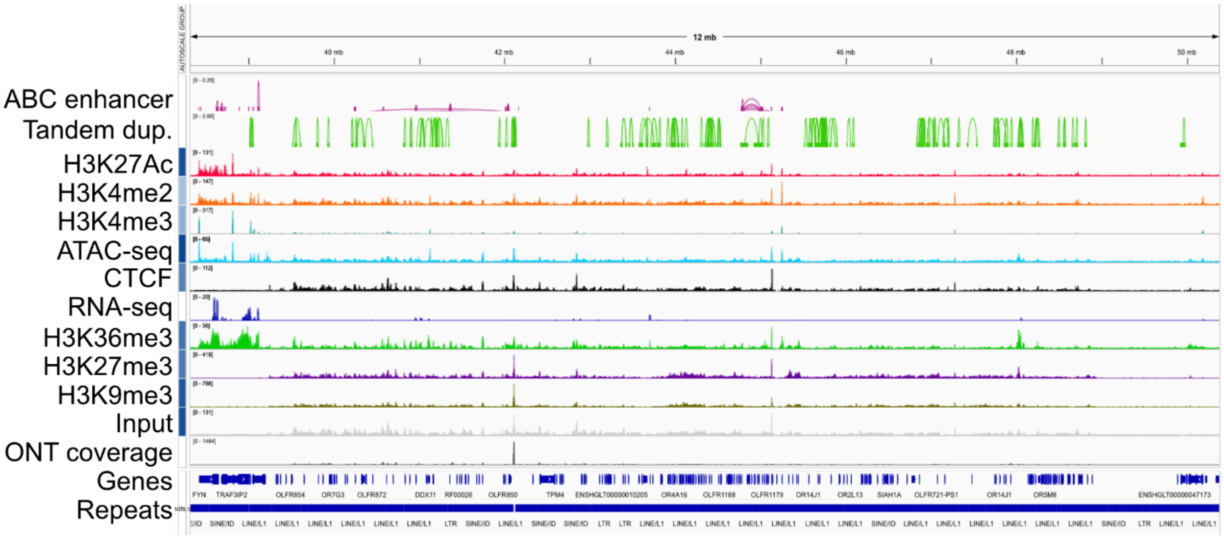

HetGlaV4. *SLC25A2*: Chr11: 88,620,294-88,625,676

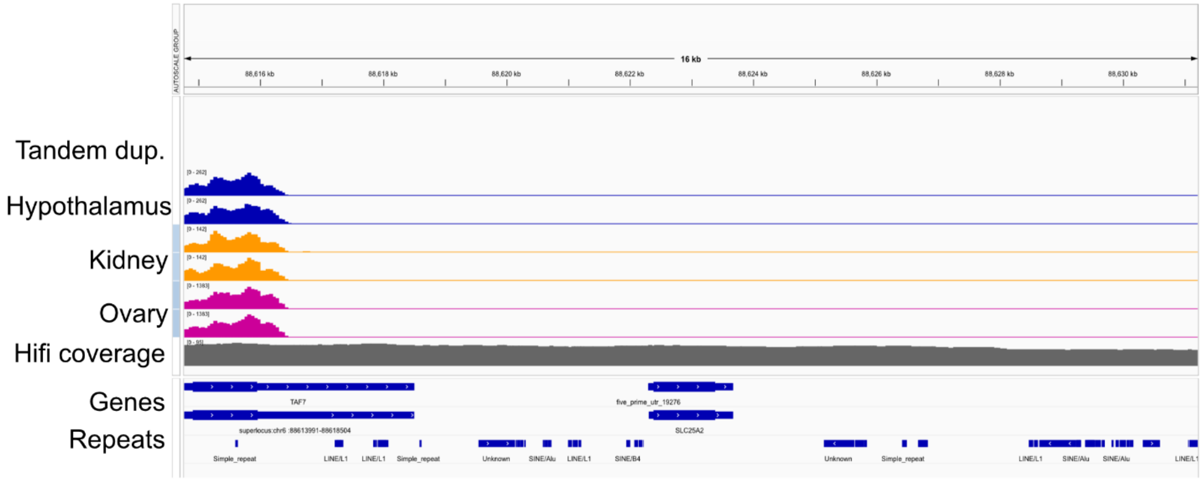


mHetGlaV3.*SLC25A2*: Chr11: 19,061,867-19,077,013

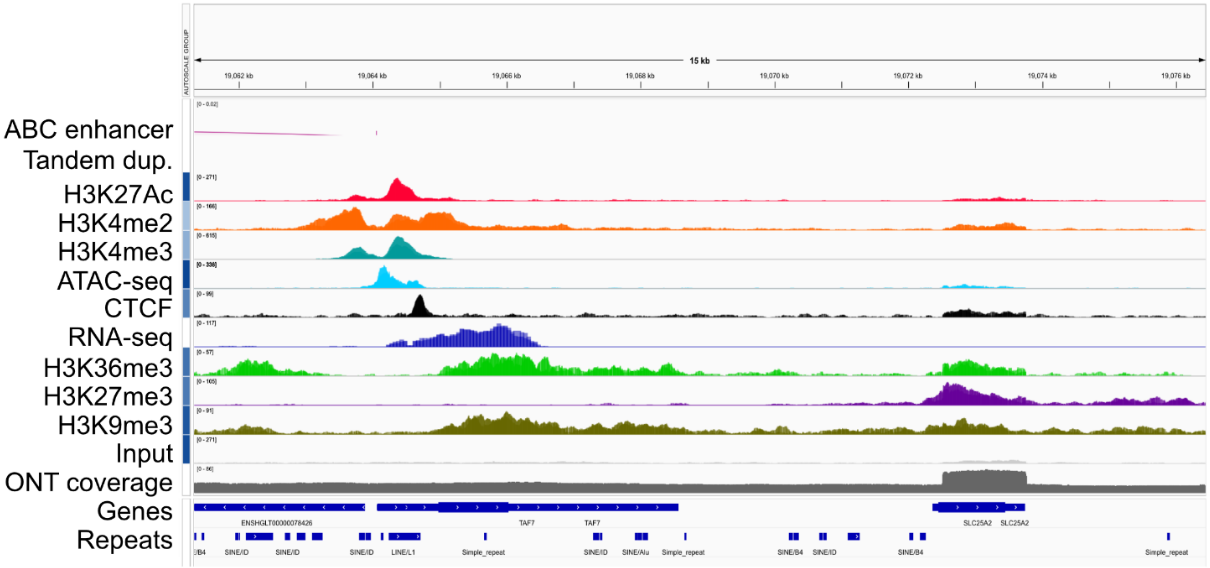

HetGlaV4. *TSSK4, CHMP4A, MDP1, NEDD8, GMPR2, TINF2*: Chr1: 18,529,596-18,625,443

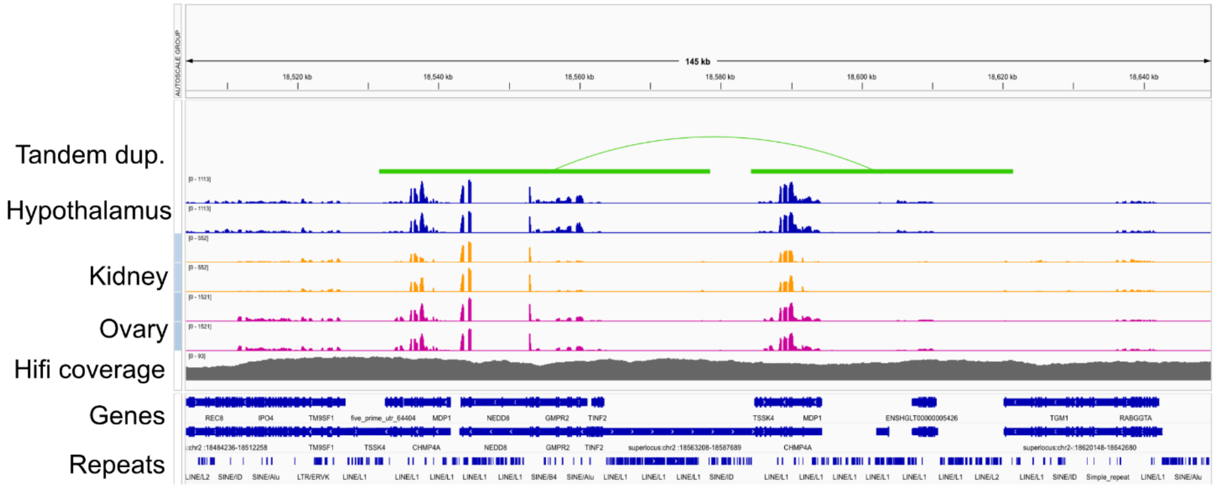


mHetGlaV3. *TSSK4, CHMP4A, MDP1, NEDD8, GMPR2, TINF2*: Chr1: 37,353,228-37,497,191

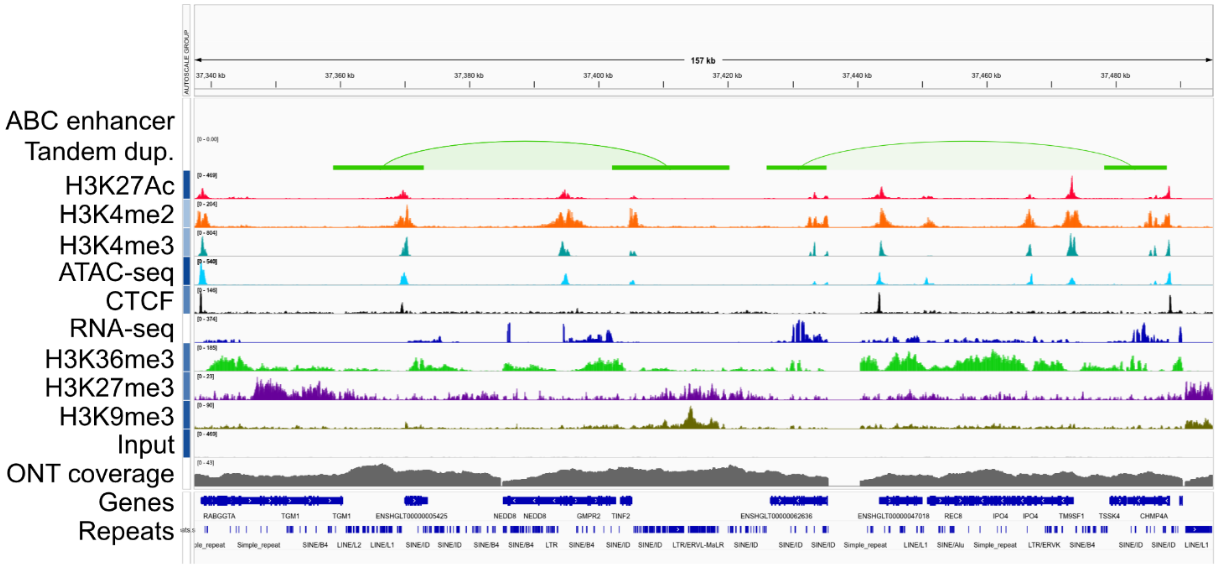


***Figure S14.*** *Four examples of where a naked mole-rat genome assembly using PacBio Hifi and ONT ultra-long reads (HetGlaV4) could resolve regions that were misassembled in mHetGlaV3.* These regions are shown on the interactive gene viewer screenshots (HetGlaV4 top, mHetGlaV4 bottom). In HetGlaV4, tracks represent RNA-seq data from various tissues while mHetGlaV3 displays the entire epigenome annotation of the NMR hypothalamus.
