## Supplementary File S2 for "An updated reference genome sequence and annotation reveals gene losses and gains underlying naked mole-rat biology"

*ARF4*: chr6:85,003,219-85,055,214

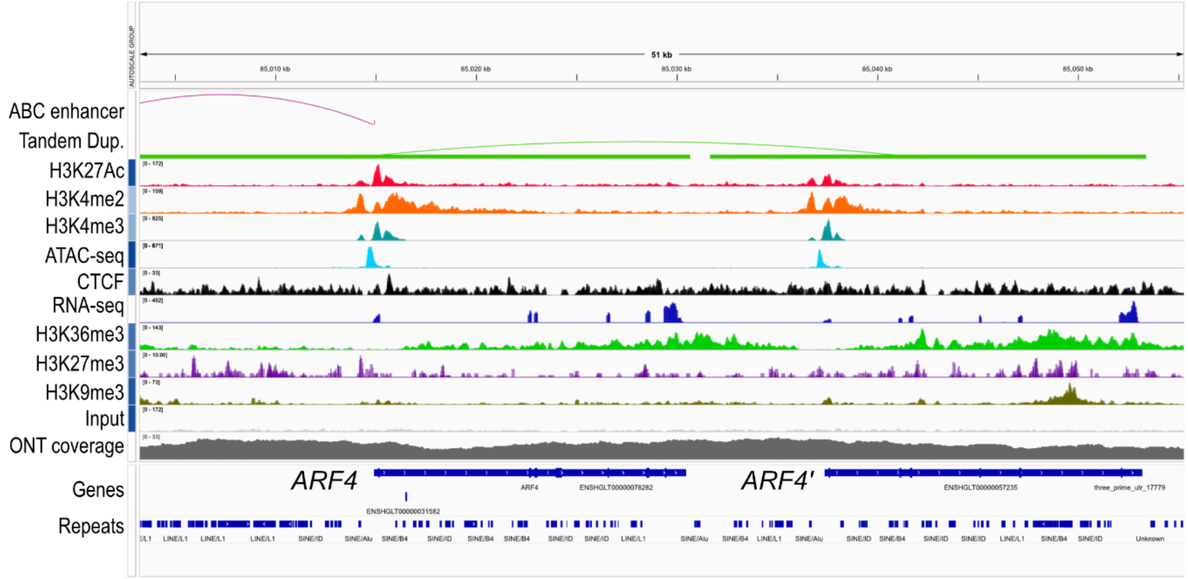


*CEL*: chr13:3,321,953-3,345,504

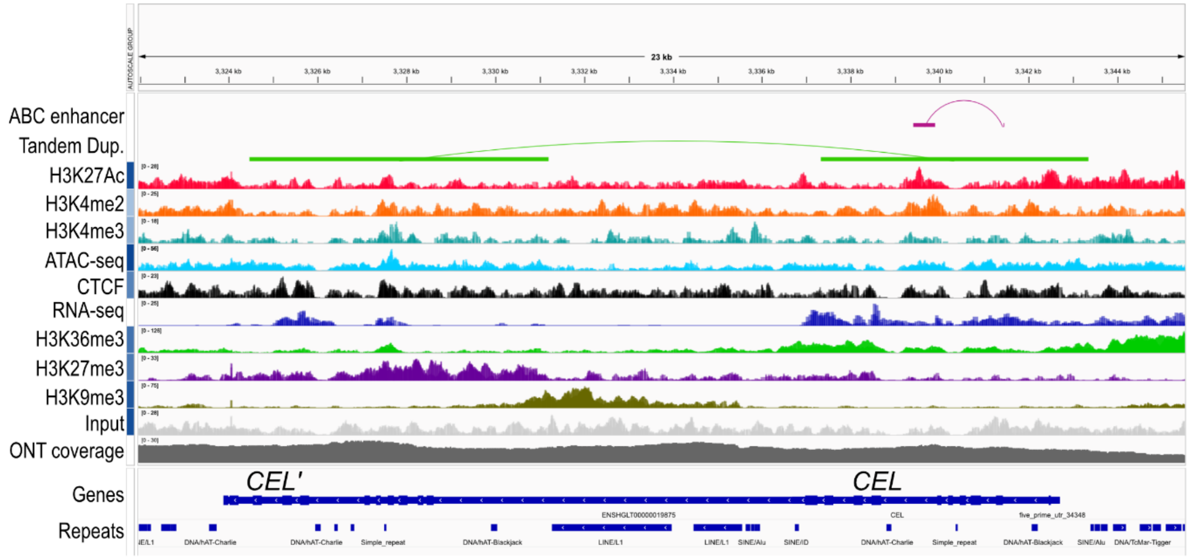

*COG1*: chr21:48,098,201-48,200,79

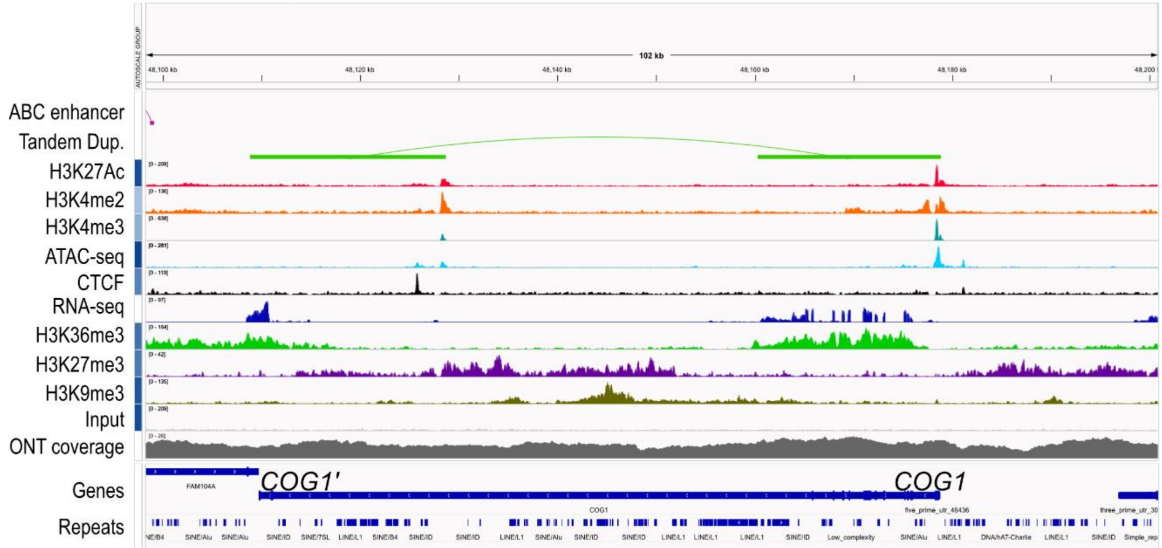


*CST6*: chr13:3,321,953-3,345,504

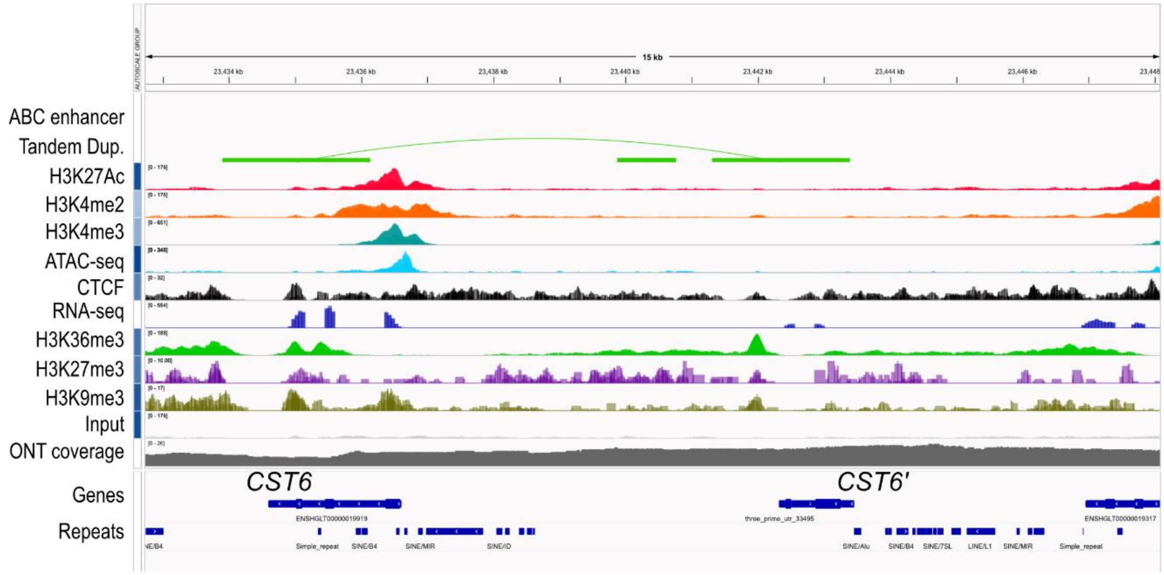

*CTRB1*: chr7:91,436,127-91,492,524

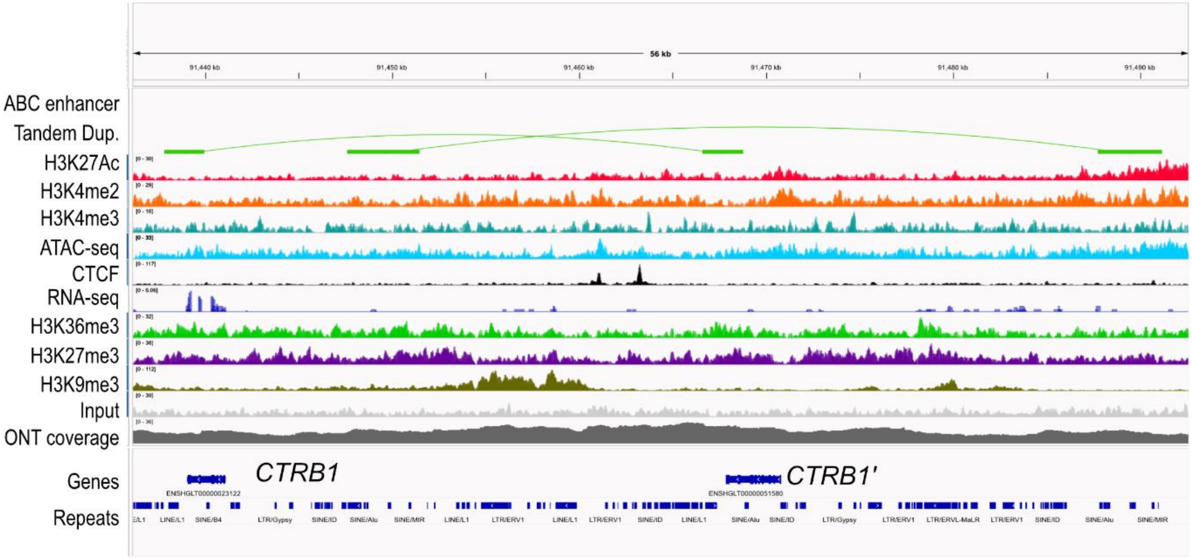


*DDT*: chr23:38,197,873-38,313,130

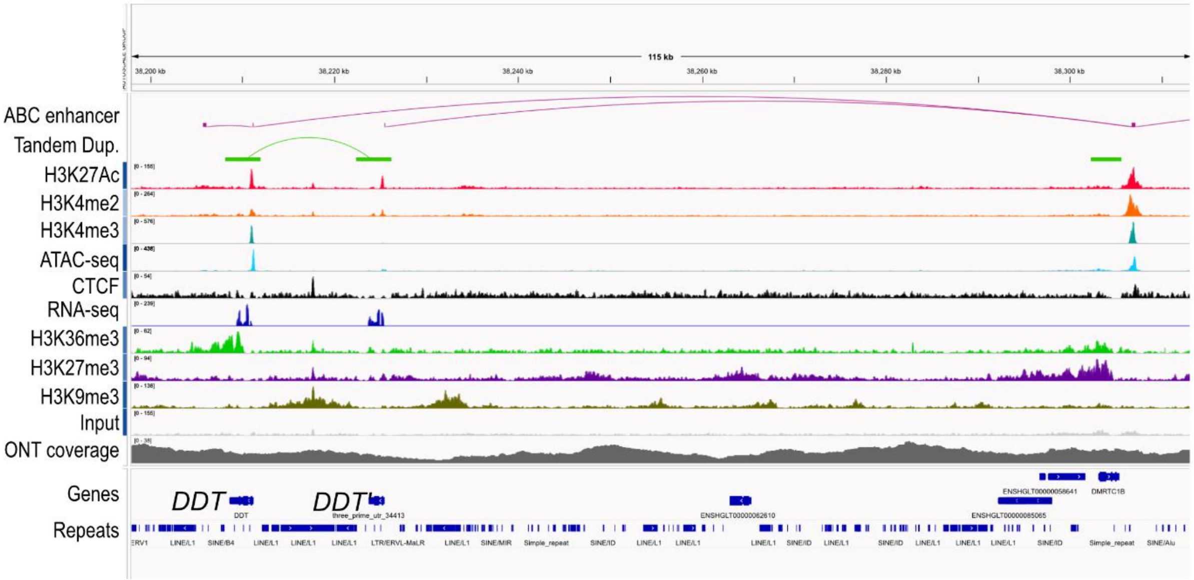

*EPHB6 TRPV5 TRPV6 LLCFC1 KEL*: chr20:46,648,036-46,963,435

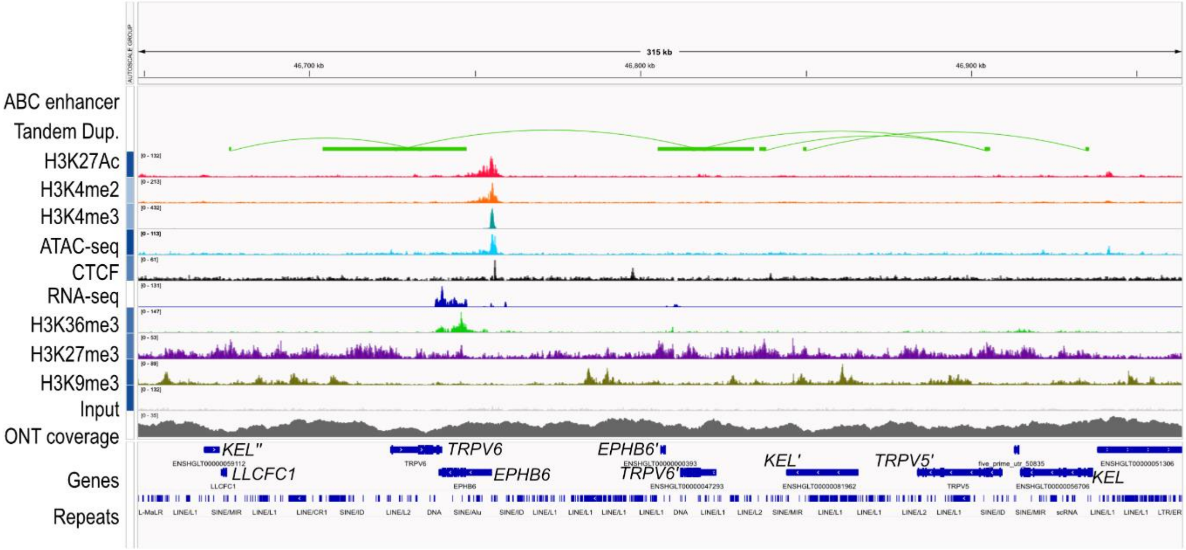


*FABP1*: chr4:57,701,358-57,769,171

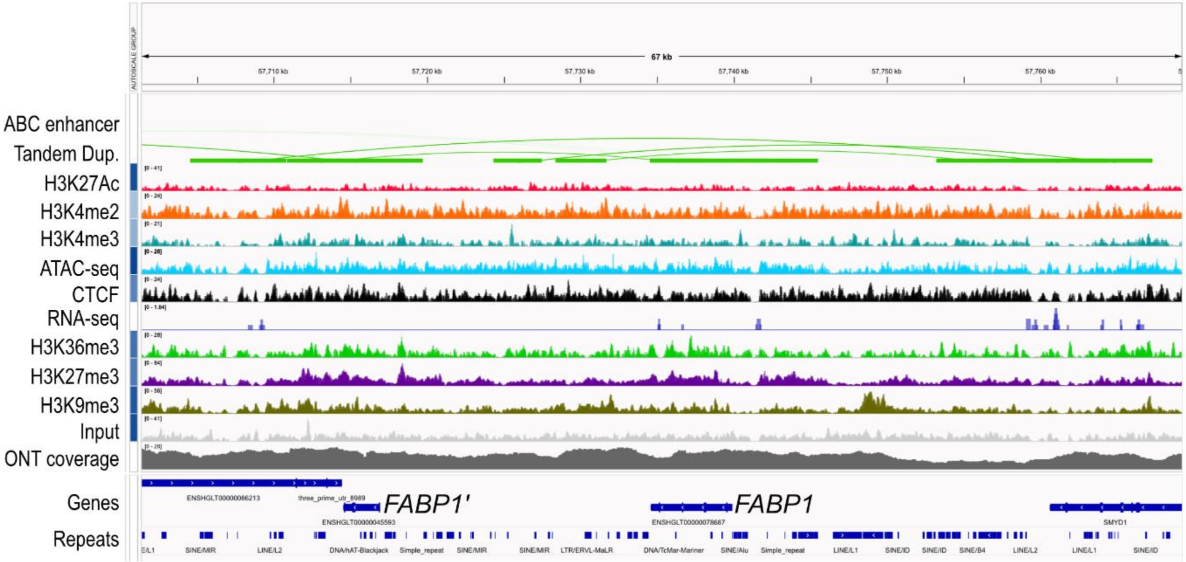

*FMO4*: chr24:15,893,506-16,009,869

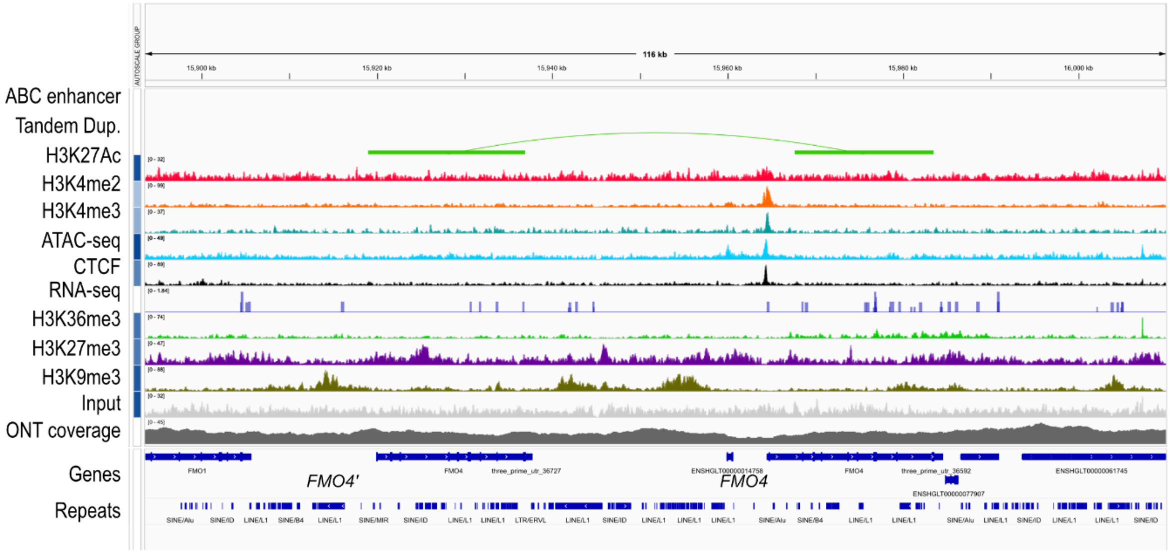


*KCTD14*: chr5:89,138,875-89,170,998

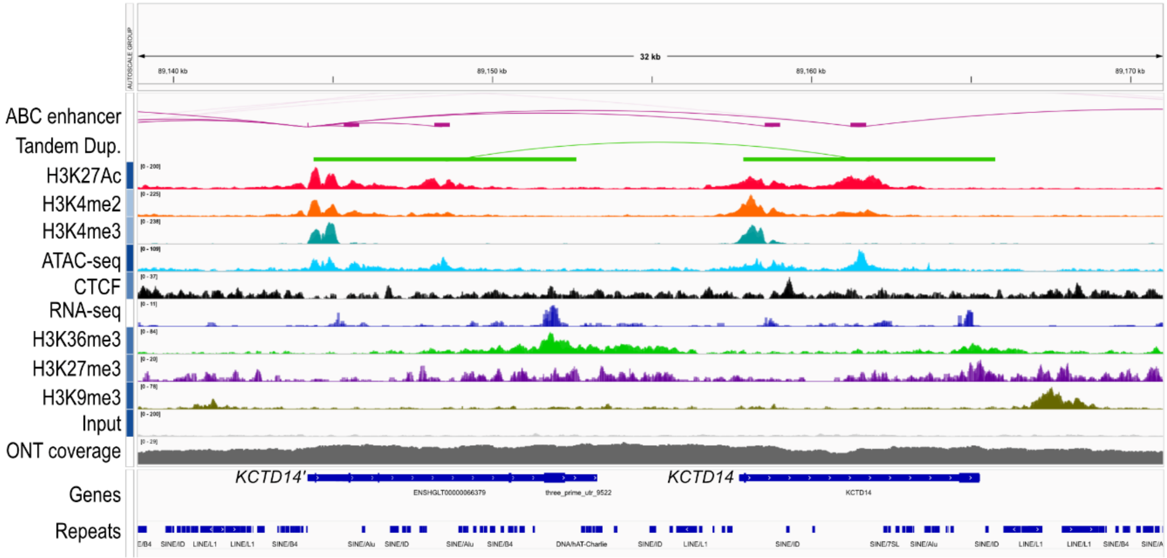

*KYAT1*: chr13:6,086,859-6,133,961

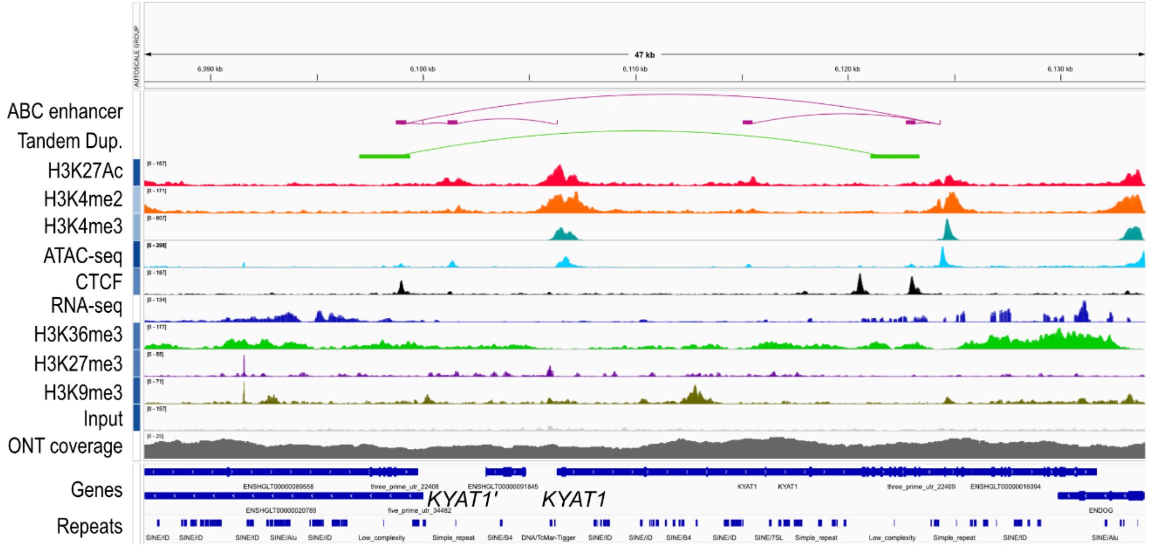


*LCN2*: chr13:7,349,575-7,373,773

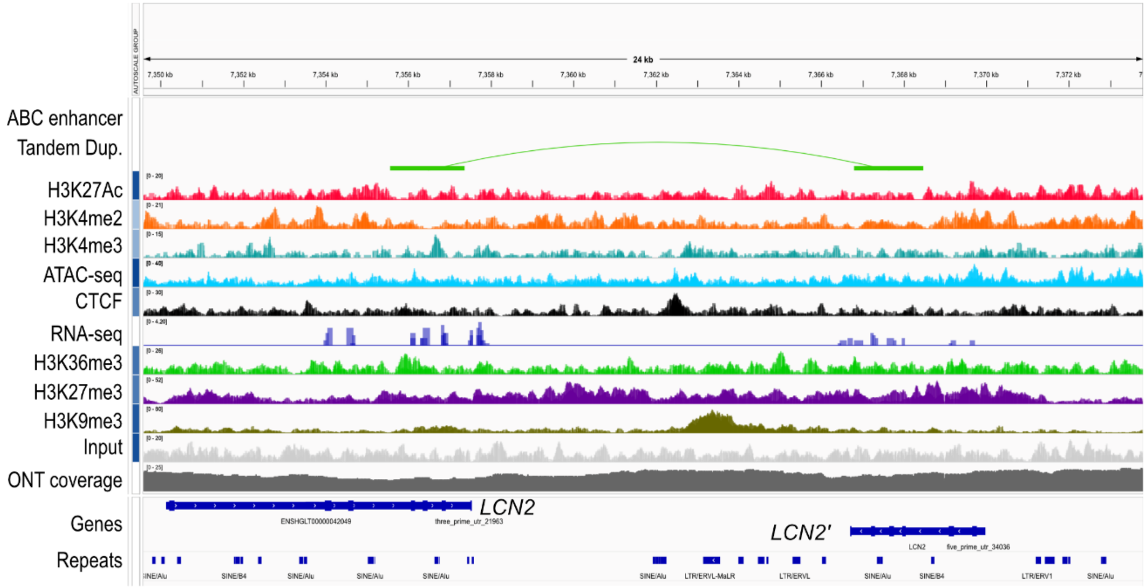

*NUDT16*: chr11:32,214,427-32,268,393

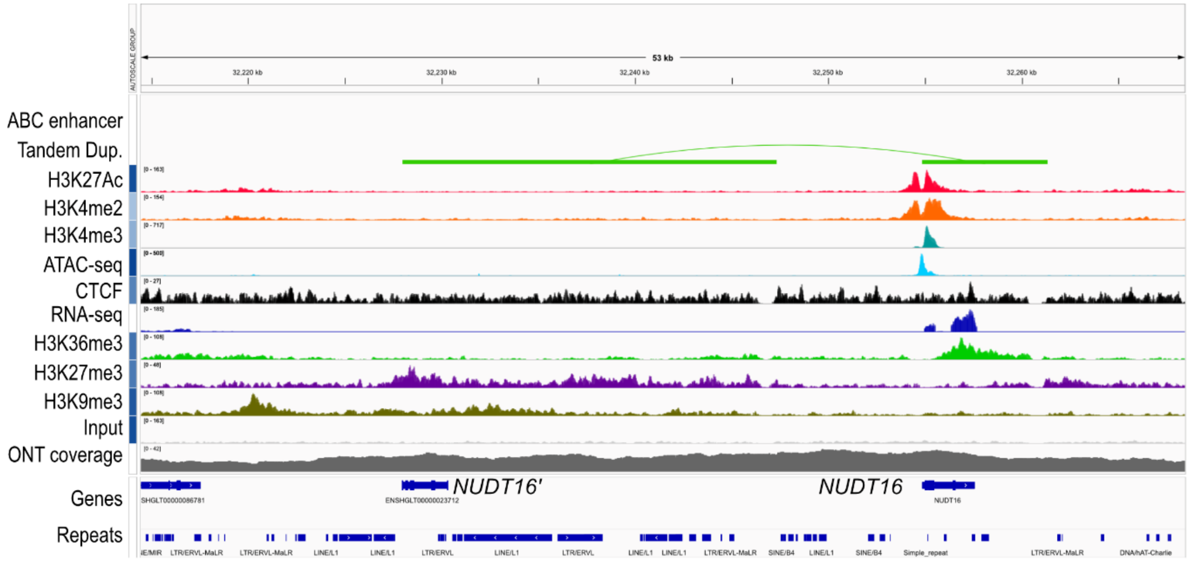


*SPIN2B*: chrX:62,619,182-62,657,958

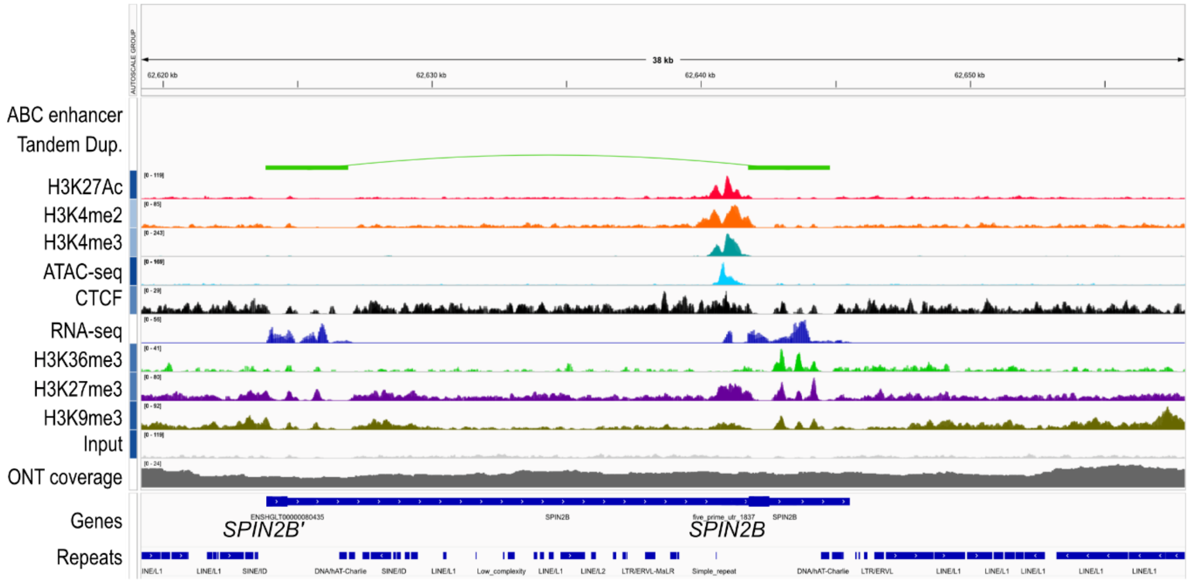

*TCP1*: chr2:87,715,383-87,860,850

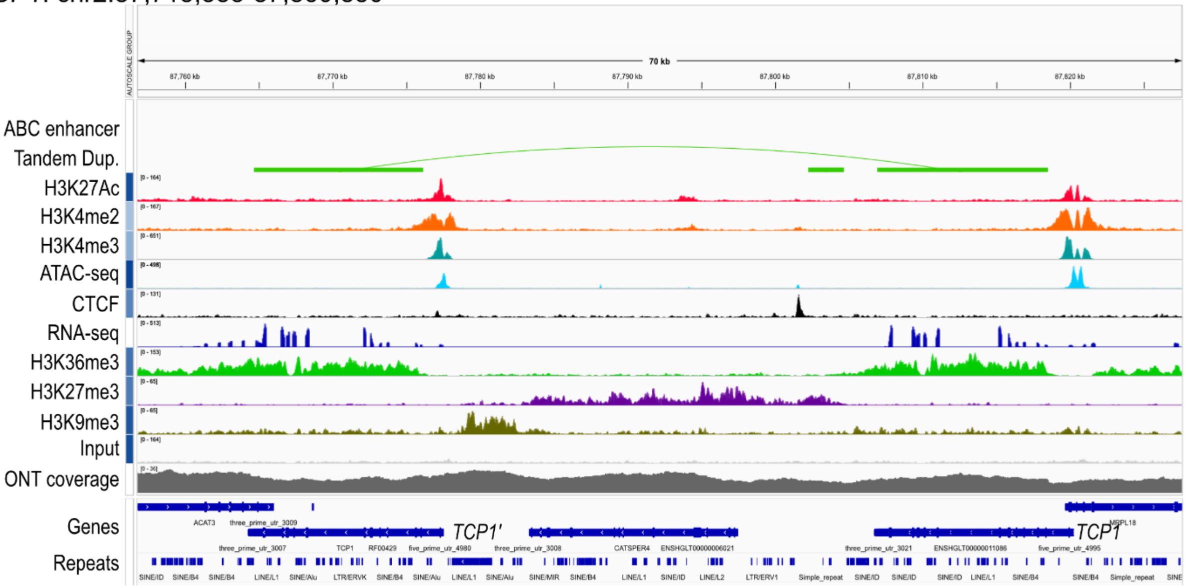


*TINF2*: chr1:37,352,142-37,425,908

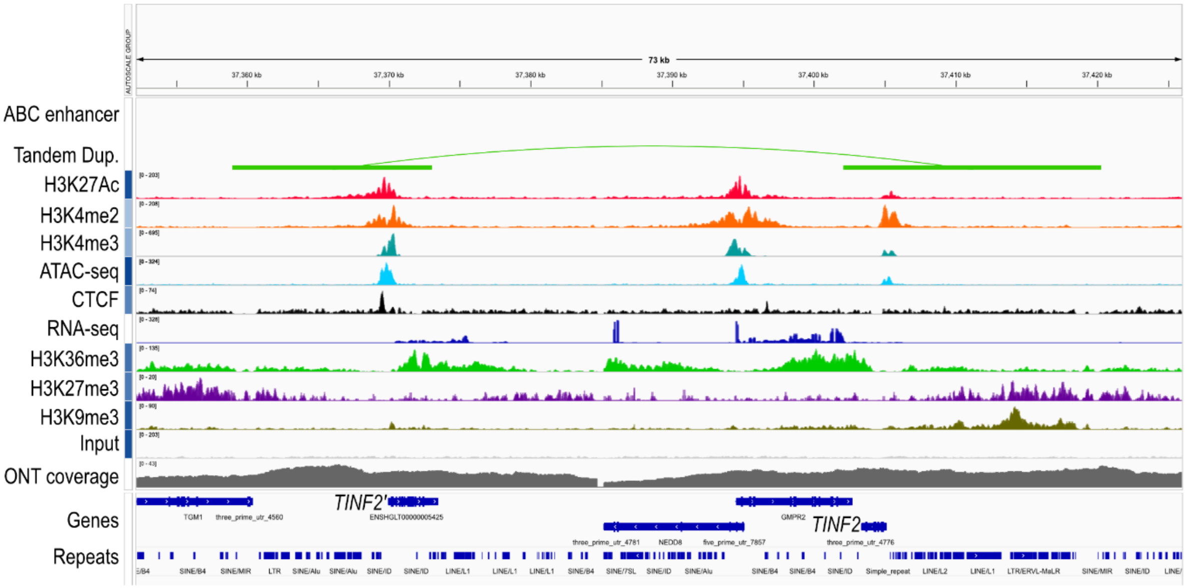
